# Profiling and modulating astrocyte borders at injected biomaterials in mice

**DOI:** 10.64898/2026.08.26.747354

**Authors:** Eric M. DuBois, Kunyu Li, Patrick Kulaga, Laboni F. Hassan, Honour O. Adewumi, Isaac C. Herrick, Kayla Dunson, Timothy M. O’Shea

## Abstract

Astrocyte border formation is a conserved neuroprotective response to neural tissue disruption, yet astrocyte border states at implanted biomaterials remain less well characterized than injury responses. Here, we developed the Astrocyte Border Characterization (ABC) Tool, which leverages a shear-thinning, injectable biomaterial to locally deliver astrocyte-specific RiboTag AAVs and small molecule regulators in the mouse striatum, enabling molecular profiling and phenotypic modulation of astrocyte border (AB) cells. Spatially precise delivery of AAV using the ABC Tool yielded enhanced specificity and robust RiboTag expression in AB cells from 7-70 days post injection. Temporal transcriptomic profiling of AB cells revealed predominantly acute, transient changes in genes governing dedifferentiation, proliferation, metabolic reprogramming, and inflammation regulation. Persistent changes accounted for only 14% of regulated genes but involved critical gain of functions in immune regulation and host defense that mirrored astrocyte border responses at chronic CNS injuries. Local delivery of indiscriminate or astrocyte-selective ablation molecules delayed, rather than prevented, border formation, ultimately yielding thicker astrocytes borders with increased inflammation and fibrosis at the biomaterial-tissue interface. Conversely, local delivery of β-hydroxybutyrate (BHB) from the ABC Tool altered key aspects of the transcriptional reprogramming to attenuate chronic astrocyte reactivity and prevent biomaterial contraction without exacerbating inflammation or fibrosis. Our findings establish the ABC Tool as a bioassay for studying and manipulating astrocyte borders at implanted biomaterials and identify focal metabolic regulation as a strategy to modulate AB cell phenotypes and enhance the CNS biocompatibility of biomaterials.

## Introduction

As the predominant glial cells in the mammalian central nervous system (CNS), astrocytes play essential roles in maintaining neural circuit activity and tissue homeostasis^1^. Astrocytes accomplish this by tiling neural parenchyma in a non-overlapping manner and interacting directly with neurons to regulate synaptic plasticity and transmission, waste recycling, and metabolism^2–5^. Specialized astrocyte populations at large blood vessels and the outermost parenchymal layers form selective barriers that regulate cell and molecular interactions with adjacent non-neural compartments^6^. These astrocytes assemble into interconnected networks of overlapping cells to form astrocyte borders termed the *glia limitans*^7^. Reprogramming of non-overlapping neuron-interacting astrocytes into interconnected *glia limitans*-like astrocyte borders is an evolutionarily conserved, adaptive host response to trauma^6,8^, ischemia^9^, infection^10^, neoplasm^11^, autoimmunity^12^, and neurodegeneration^13^. To transition into border states, astrocytes must transiently re-enter the cell cycle and proliferate^14^, reduce regular neuronal support capabilities^15,16^, increase immune regulatory functions^17^, and conduct major shifts in metabolic pathway usage^18,19^. Targeted loss-of-function manipulations that disrupt these reprogramming processes lead to unrestricted spread of destructive inflammation or protein aggregates that exacerbate neural tissue damage, demonstrating the essential neuroprotective role of astrocyte borders ^8,9,12,13,20–23^.

Beyond CNS disorders, implantation of medical devices such as neural electrodes or drug delivery systems into the CNS elicits a conserved foreign body response (FBR) that also causes astrocytes to adaptively reprogram into border states^24–26^. The scale of the astrocyte reprogramming dictates the extent of neural tissue disrupted around implanted devices, ultimately determining neural device performance and longevity. Moreover, device micromotion, latent infection, or other persistent device-derived stimuli can further intensify astrocyte border formation and accelerate degradation of device function^24,27–29^. Although astrocyte border formation is a common feature of CNS disorders and a centerpiece of device FBR, conserved and context-specific functions of these astrocyte borders remain poorly understood. Furthermore, while we and others have extensively studied astrocyte transcriptomic changes across many different disorders where astrocyte border formation is prominent ^29–34^, precisely decoupling the transcriptional profiles of border astrocytes from other non-proliferative astrocyte reactivity states has been challenging, since transgenic and viral-based approaches indiscriminately label both astrocyte populations. The advent of single-cell sequencing and spatial transcriptomic technologies has improved discrimination of border astrocytes from other astrocyte states^29,31^, but limited read depth and processing artifacts with these techniques still hinders thorough molecular characterization^35,36^.

Improving our understanding of astrocyte border biology requires new approaches to selectively analyze and manipulate astrocyte border (AB) cells. Previously, we demonstrated that focal injection of non-resorbable, soft biomaterials into mouse striatum triggers astrocyte reprogramming into border states within a narrow zone of tissue adjacent to the biomaterial surface^24,37^. A consequential finding from these prior studies was that molecular cargo released from specific injected biomaterial formulations was restricted to a defined zone adjacent to the biomaterial surface, corresponding to the tissue region occupied by newly formed border astrocytes^24,37^. These observations motivated us to explore whether we could use biomaterials delivering AAVs and small molecule regulators to directly label, manipulate, and analyze striatal astrocytes undergoing transcriptional reprogramming into border states in response to biomaterial injection.

Here, we developed and validated the Astrocyte Border Characterization (ABC) Tool, which involved delivering AAV5 GfaABC1D-Rpl22HA from non-resorbable, viscoelastic, methylcellulose (MC)- based biomaterials to enable astrocyte-specific expression of a ribosomal protein-hemagglutinin tag (RiboTag) in a spatially restricted population of striatal astrocytes undergoing adaptive reprogramming into border states. We assessed RiboTag-labeled astrocyte borders morphologically by immunohistochemistry and conducted translational profiling of AB cells by RiboTag immunoprecipitation (IP)-enriched RNA sequencing at timepoints up to 70 days post-injection. We found that biomaterial-adjacent astrocytes undergo temporally dependent morphological changes that are associated with transient transcriptional alterations in genes regulating dedifferentiation, proliferation, metabolic reprogramming, and inflammation, as well as persistent changes in a smaller subset of genes that are conserved in astrocyte borders at chronic CNS injuries. Local delivery of indiscriminate or astrocyte-selective ablation molecules from biomaterials disrupted astrocyte border formation, leading to increased inflammation and fibrosis. Meanwhile, delivery of astrocyte-targeted metabolic regulators attenuated border formation without exacerbating peripherally derived FBR processes. Our findings demonstrate that the ABC Tool can be used to study and regulate astrocyte borders and establish an effective bioassay for future screening of new therapies to enhance border formation to fortify neural tissues against disease or to attenuate borders for improved neural device integration.

## Results

### AAV delivery from injected biomaterials enhances specificity of astrocyte border labeling

Biomaterials injected into the CNS stimulate local astrocytes to transition into border states. For this study, we focused on characterizing these astrocyte responses in the striatum, because it is an easily accessible anatomic site for consistent and reproducible viral and biomaterial injections. Additionally, striatal astrocytes predominantly exhibit neuron-interacting, non-overlapping territorial domains, meaning that border state transitions involve pronounced morphological and molecular changes^24^. To determine whether the ABC Tool, a formulation of MC biomaterial delivering RiboTag AAV under the astrocyte promoter GfaABC1D, specifically labels induced astrocyte borders, we first conducted immunohistochemistry (IHC) evaluations to characterize the spatial distribution of viral expression via the detection of hemagglutinin (HA)-positive ribosomes (**Fig. 1**). The selectivity of viral expression in striatal astrocytes afforded by the AAV5 serotype and GfaABC1D promoter is well established, and our labeling of striatal astrocytes by AAV solution injections performed comparably to these previous reports^38–40^. The astrocyte viral expression by the ABC Tool was benchmarked against two experimental control groups receiving: (i) an injection of RiboTag AAV solution only (AAV), or (ii) an initial striatal injection of RiboTag AAV solution followed by an injection of MC biomaterial without loaded virus 14 days later (AAV/MC) (**Fig. 1a**). All groups were evaluated 28 days after the final surgical injection and received the same total amount of AAV (1µL, 1×10^12^ gc/ml AAV). MC biomaterials used in the ABC Tool and AAV/MC conditions were prepared at 5 wt%, representing the highest MC concentration in which formulations could be both loaded and readily extruded through the pulled glass micropipettes used for surgical injections (**Supplementary Fig. 1**). MC was selected as the biomaterial for this work because it is readily commercially available, non-resorbable *in vivo,* and elicits a FBR with a minimal non-neural compartment but a well-defined astrocyte border^24,37^. Formulated MC biomaterials demonstrated reproducible mechanical properties across multiple independent preparations, with a mean storage modulus (G’) of 129 ± 22 Pa (95% CI), tan(ẟ) of 1.96 ± 0.13 (95% CI) and stress relaxation time constant (*τ*) of 0.018 seconds, resulting in a soft, viscoelastic biomaterial (**Fig. 1b, Supplementary Fig. 1**).

**Fig. 1.**
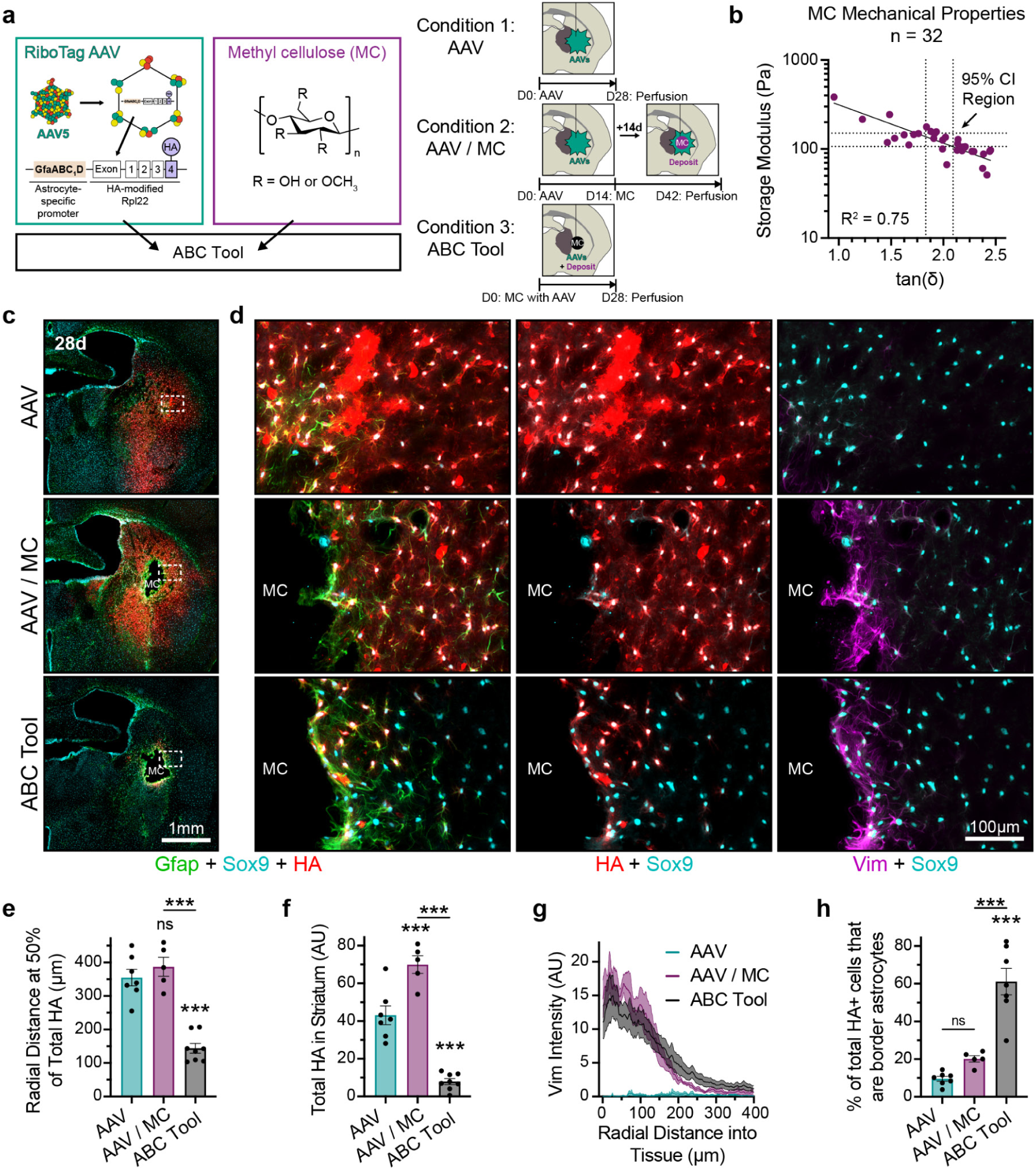
AAV delivery from injected biomaterials enhances specificity of astrocyte border labeling. **a.** Schematic of the ABC Tool, which incorporates the GfaABC_1_D-RiboTag AAV in methylcellulose (MC) biomaterials and experimental timelines for conditions 1 (AAV), 2 (AAV/MC), and 3 (ABC Tool). **b.** Plot showing storage modulus (Pa) vs tan(δ) for 5wt% MC demonstrating reproducibility of material properties across formulations (n = 32). Dashed lines show 95% Confidence Interval (CI) region. **c.** Survey images showing distribution of HA expression in striatal astrocytes at 28 days after injection of AAV solution or biomaterial. Dashed box highlights region for detail images. **d.** Detail images showing Vimentin (Vim)-positive astrocyte border formation and HA labeling for the three conditions at 28d. **e.** Radial distance at 50% of total HA expression measured from the start of the tissue. **f.** Quantification of total HA expressed by astrocytes in striatum. **g.** Plots of average vimentin intensity measured radially from the center of injection or edge of the biomaterial deposit. **h.** Percent of total HA-positive astrocytes located within the Vimentin-positive astrocyte border. Bar charts in **e.-f., h.** show not significant (ns), and ***P < 0.0006 compared to AAV or as designated, via one-way ANOVA with Tukey’s multiple comparison test. Graph shows mean ± s.e.m. with individual data points showing AAV (n = 7), AAV / MC (n = 5), and ABC Tool (n = 8 (**e.-f.**) or n=7 (**h.**)).

MC biomaterials in both the ABC Tool and AAV/MC condition persisted as focal deposits in the striatum when evaluated at 28 days post injection (**Fig. 1c-d**). Infusion of RiboTag AAV solution caused widespread transduction and HA expression in Sox9-positive astrocytes across essentially the entire striatum (**Fig. 1c)**. Conversely, the ABC Tool showed spatially restricted delivery of AAV such that only astrocytes immediately adjacent to the biomaterial surface had detectable HA expression, with the 50th percentile of HA-labelled astrocytes located less than 150µm from the biomaterial surface (**Fig. 1d-e**). The spatially restricted delivery of AAV using the ABC Tool resulted in 86% less total HA detected within the striatum compared to both the AAV only and AAV/MC controls (**Fig. 1f**). Astrocyte border state transitions involve the formation of non-overlapping domains and persistent upregulation of the intermediate filament protein, Vimentin, which we used to identify astrocyte border (AB) cells generated around injected MC biomaterials^29^. Both the ABC Tool and AAV/MC groups, but not the AAV solution only condition, stimulated the formation of Vimentin-positive astrocyte borders that were wholly contained within a discrete zone, 250 µm from the biomaterial surface (**Fig. 1d,g**). Using the ABC Tool, approximately 80% of the total detected HA was constrained within the 250 µm border region (**Supplementary Fig. 2**). By contrast, only 30% of total detected HA in the AAV/MC condition was contained within the border zone, with most HA expression being in Vimentin-negative, striatal astrocytes maintaining non-overlapping cellular domains within the bulk of the preserved striatal neuropil. Using the ABC tool, 80% of HA- and Sox9-positive cells within the 250 µm border zone expressed Vimentin and essentially all labeled cells displayed overlapping domains consistent with a border astrocyte phenotype (**Supplementary Fig. 2**). Notably, less than 60% of HA-positive cells in the border zone of AAV/MC mice were Vimentin positive and at least 25% of HA-positive cells were single-domain astrocytes (**Supplementary Fig. 2**). Thus, of all the HA-positive cells labeled by the ABC Tool at 28 days post injection (dpi), we conservatively estimated that at least 60% were border astrocytes, representing over a 3-fold enrichment for astrocyte border cells compared to the AAV/MC condition (**Fig. 1h**). ABC Tool injections caused no change in mouse behavior or activity levels compared to AAV solution only injections, suggesting that the striatum-persisting biomaterials induced highly localized tissue effects and did not broadly impact mouse brain functions (**Supplementary Fig. 3**).

These data show that the ABC Tool generates discrete astrocyte borders that form around the persistent biomaterial deposit to permit enhanced astrocyte border-specific RiboTag labeling compared to viral solution injections by spatially restricting AAV delivery to the astrocyte border region.

### The ABC Tool enables translational profiling of astrocyte border states at injected biomaterials

Since the ABC Tool permitted highly specific expression of RiboTag in AB cells, we next used the tool to conduct translational profiling of AB cells around MC biomaterials at 28 dpi. RiboTag immunoprecipitation (IP) applied to homogenized striatal tissue biopsied from around MC deposits was used to isolate HA-positive ribosomes from border astrocytes. Subsequently, mRNA that was actively being translated within these ribosomes was purified and sequenced (**Fig. 2a**). To determine the extent of the adaptive reprogramming involved in forming astrocyte borders, we also performed RiboTag IP and RNA sequencing on animals receiving: (i) the AAV solution only condition to profile healthy striatal astrocytes, and (ii) the AAV/MC condition to account for global reactive changes following biomaterial injection including those derived from border and non-overlapping, domain persisting astrocytes in the neuropil. Sequencing of mRNA from each sample’s flow-through solution after RiboTag IP was used to characterize gene expression by non-astrocyte striatal cells contained within the biopsied tissue region and to assess astrocyte enrichment afforded by RiboTag IP. We recovered approximately 50ng of astrocyte border mRNA on average from a single brain using the ABC Tool, which was approximately 33% of the total mRNA extracted from the AAV solution-injected striatum and 12% of the total mRNA recovered from the AAV/MC condition, which was consistent with the magnitude of the differences in total HA detected by IHC (**Fig.1f, 2b**). The greater amounts of mRNA recovered from the AAV/MC condition likely reflects elevated mRNA synthesis levels associated with reactive changes following biomaterial injection (**Fig. 2b**). The total RiboTag IP mRNA extracted using the ABC Tool was less than 2.5% of the total mRNA recovered from the flow-through samples, demonstrating the significant selectivity of the IP method. Across all three conditions, there was no difference in the amount of mRNA recovered in flow-through samples, reflecting consistencies in the volumes of striatal tissue being biopsied (**Supplementary Fig. 4**). All RiboTag IP samples showed highly enriched expression of the 3xHA sequence transcripts associated with the virally expressed Rpl22-HA, whereas the flow through samples had no (ABC Tool) or very limited (AAV and AAV/MC) detectable 3xHA transcripts (**Fig. 2c, Supplementary Fig. 4**).

**Fig. 2.**
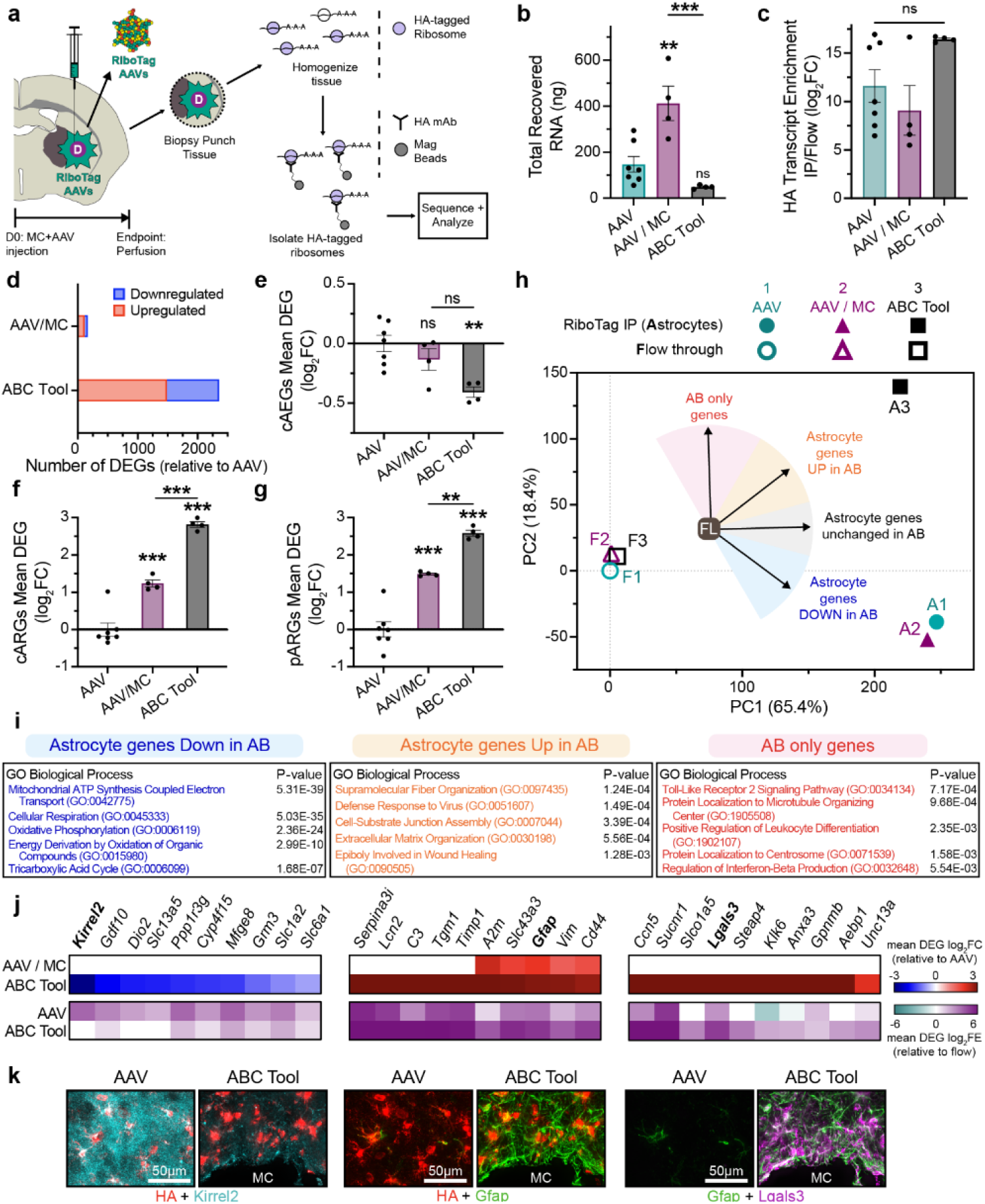
The ABC Tool enables translational profiling of astrocyte border states at injected biomaterials. **a.** Schematic showing RiboTag immunoprecipitation (IP) and RNA sequencing workflow for evaluating mRNA expression in Astrocyte Border (AB) cells. **b.** Total RNA recovered from the RiboTag-IP process for all conditions. **c.** Relative enrichment of 3xHA in RiboTag-IP samples relative to flow samples. **d.** Bar chart showing the number of differentially expressed genes (DEGs) relative to AAV (IP) for conditions involving MC biomaterial injection. **e-g.** Mean differential expression (log_2_FC) across conditions for curated gene lists for: **e.** consensus healthy astrocyte-enriched genes (cAEGs) **f.** consensus astrocyte reactivity genes (cARGs) and **g.** persisting astrocyte reactivity genes (pARGs). **h.** PCA of 20931 DEGs across RiboTag-IP and Flow through samples across the three conditions, with the 4 categories of AB gene lists derived from the factor analysis based on Factor Loading Vector Cartesian Angle (FLVCA) shown. **i.** Table showing significant GO-BP terms for astrocyte genes down in AB, astrocyte genes up in AB, and AB-only genes. **j.** Top heatmap shows select DEGs from the astrocyte genes down in AB, astrocyte genes up in AB, and AB-only gene categories for the AAV/MC and ABC Tool condition. Bottom heatmap shows mean enrichment in IP samples relative to Flow for the AAV and and ABC Tool conditions **k.** Detail images showing colocalization of HA-tagged ribosomes in astrocytes with Kirrel2 and Gfap, as well as Gfap and Lgals3, in healthy and border astrocytes. All bar charts in this figure (**b.-c., e.-g.**) show not significant (ns), **P < 0.005, and ***P < 0.0007 compared to AAV or as designated via a one-way ANOVA with Tukey’s multiple comparison test. Graph shows mean ± s.e.m. with individual data points showing AAV (n = 7), AAV / MC (n = 4), and ABC Tool (n = 4). All transcriptomic data shown in this figure is derived from the following sequenced sample set: IP: AAV (n = 7), AAV / MC (n = 4), and ABC Tool (n = 4). Flow: AAV (n = 7), AAV / MC (n = 4), and ABC Tool (n = 4).

Using the ABC Tool, we identified 2357 unique differentially expressed genes (DEGs) in astrocyte border cells compared to healthy striatal astrocytes, with 1489 upregulated and 868 downregulated DEGs. Notably, we detected relatively few DEGs in the AAV/MC condition compared to healthy astrocytes, with only 114 upregulated and 51 downregulated DEGs, representing just 7% of the total DEGs detected by the ABC Tool (**Fig. 2d**). The increased number of detected DEGs with the ABC Tool is consistent with three results: (i) that the ABC Tool is permitting the sequencing of the AB cell population with enhanced enrichment compared to AAV/MC condition; (ii) that AB cells at injected MC biomaterials are persistently transcriptionally distinct from astrocytes remaining within the striatal neuropil; and (iii) that reactive changes in neuropil astrocytes following a single perturbing event (i.e. a biomaterial injection) are transient, such that this predominant astrocyte population labeled in the AAV/MC condition has mostly returned to near pre-injection states by the 28 dpi evaluation timepoint, consistent with previous findings^41^.

To characterize the nature of AB cell DEGs identified by the ABC tool at 28 dpi, we first examined the expression levels for gene panels of consensus healthy astrocyte-enriched genes (cAEGs) and astrocyte reactivity genes (cARGs), which are lists of genes that we previously derived using meta-analysis of archival astrocyte specific mRNA datasets^29,42^, and which have been adopted by others in the field to characterize astrocyte responses^43^. We also collated a list of persisting astrocyte reactivity genes (pARGs) containing transcripts that we have noted to be chronically elevated and highly enriched in astrocyte borders at CNS injuries^29^ (**Fig. 2e-g**). In the AB cells labelled by the ABC tool, over 30% of cAEGs (132 out of 429) were detected as DEGs, with the majority (85%) being downregulated, leading to a significant reduction in mean expression of cAEGs in AB cells compared to healthy astrocytes (**Fig. 2e**). Downregulated cAEGs in AB cells included molecules involved in fatty acid metabolism (*Acaa2*, *Gstm1, Acsbg1, Acsl6, Slc27a1, Slc25a34, Elovl2*), synapse modulation (*Kirrel2, Gpc5, Gpc6*), and astrocyte maturation (*Cntfr, Fzd2, Cables1, Nrarp*) (**Supplementary Fig. 5**). Strikingly, there were many established canonical striatal region-specific astrocyte genes not featured on the cAEG list that were also prominently downregulated in AB cells including *Crym*, *Prss56, Ppp1r1b, Aldh5a1, Slc25a3,* and *Slc6a3*^39,44^, indicating a pronounced loss in regionally specialized astrocyte functions as part of the transition into AB cell states (**Supplementary Fig. 5**). Less than 2% of cAEGs (8 out of 429) were detected as DEGs in the AAV/MC condition with mean expression of cAEGs being no different than healthy astrocytes, demonstrating that AB cells undergo more pronounced astrocyte dedifferentiation than domain persisting reactive astrocytes.

Astrocytes from both the ABC Tool and the AAV/MC condition had elevated expression of cARGs and pARGs compared to healthy striatal astrocytes. However, we detected at least 5-fold more DEGs using the ABC Tool for both the cARGs (83 vs 12 out of 170) and pARGs (118 vs 22 out of 259) and a mean DEG expression level for both gene lists that was essentially double that of the AAV/MC condition (**Fig. 2f-g, Supplementary Table 1**). Notable upregulated cARGs expressed by AB cells included well studied astrocyte reactivity-associated molecules (*Gfap, Vim, Serpina3n, Timp1, Lcn2*)^29,37^ as well as molecules involved in regulating inflammatory responses (*Cd44, Trem2, Ptx3, Tlr4, Cd14, Fcgr2b, Cd180*), cytokine signaling (*Ccl3, Icam1, Osmr, Anxa2, Cxcl16*), and complement pathway activation (*Cd4b, C1qa, C1qb, C1qc, C3ar1*) (**Supplementary Fig. 5**). Additionally, pARGs highly expressed in AB cells included several molecules that have been proposed as potential border markers in the context of CNS injury including *C3, A2m, Tgm1, Cdsn, H2-ab1,* and *Cd74*^29^.

To identify the gene expression profiles that distinguish AB cells from other neural cell types in addition to healthy and reactive astrocyte states, we next performed principal component analysis (PCA) on an ensemble of 20,931 genes differentially regulated across RiboTag IP and Flow-through samples from the three conditions. PCA revealed a clear separation of samples based on astrocyte enrichment along PC1 and the AB cell state along PC2 (**Fig. 2h, Supplementary Fig. 4**). Flow-through samples for all three conditions were closely associated together and overlapped in Euclidean PCA space suggesting that the population of sampled non-astrocyte cells within the striatum were mostly unaltered by the biomaterial implantation, at least when assessed at 28 dpi. By contrast, RiboTag IP samples from the ABC Tool clearly separated from the AAV and AAV/MC samples which both clustered together (**Fig. 2h**). Applying an unsupervised analysis of cell type representation using the genes with a PC1 factor loading (FL) >0.9 resulted in identification of astrocytes with high confidence using the curated Cell Marker database^45^, while genes with FL <-0.9 identified other neural cells using the same database, supporting effective enrichment of astrocyte mRNA and de-enrichment of mRNA from other cell types using the RiboTag IP as noted in our prior CNS injury studies (**Supplementary Fig. 4**)^29^.

We applied a detailed factor analysis on the PCA plot by profiling the genes with FL vector magnitudes >0.8 with different relative contributions in PC1 and PC2 directions to reveal 4 main categories of genes defining the translatome of AB cells based on their FL Vector Cartesian Angle (FLVCA): (i) astrocyte enriched genes that did not change in AB cells compared to healthy astrocytes (2604 genes; FLVCA = ±15°); (ii) astrocyte enriched genes downregulated in AB cells relative to healthy astrocytes (1521 genes; FLVCA = −15 to −60°); (iii) astrocyte enriched genes that were further upregulated in AB cells relative to healthy astrocytes (1936 genes; FLVCA = +15 to +60°); and (iv) genes that were only upregulated and enriched in AB cells (1659 genes; FLVCA = +60 to 120°) (**Fig. 2h, Supplementary Table 2**). Unsupervised analysis of Gene Ontology Biological Processes (GO-BPs) revealed that astrocyte enriched genes downregulated in AB cells (*Astrocyte genes Down in AB*) pertained to several oxidative metabolic processes including molecules critical to the oxidative phosphorylation, Tricarboxylic Acid (TCA) cycle, and electron transport chain (ETC) (**Fig. 2i**).

Additionally, astrocyte enriched genes downregulated in AB cells included transporters for glutamate (*Slc1a2*), gamma-aminobutyric acid (GABA) (*Slc6a1*), citrate (*Slc13a5*), and mitochondrial carrier family (*Slc25a4, Slc25a17, Slc25a19*), while molecules that facilitate synapse elimination (*Mfge8*) and formation (*Gdf10*), thyroid hormone availability (*Dio2*), and glutamate uptake (*Grm3*) were also prominently downregulated (**Fig. 2j, Supplementary Fig. 6**). Notably, none of these genes were detected as significantly altered in the AAV/MC condition.

Genes identified to be astrocyte enriched and upregulated in AB cells by the FLVCA analysis (*Astrocyte genes Up in AB*) included some cARGs (28 of 170) and pARGs (100 out of 259) such as canonical reactivity molecules like *Lcn2*, *Tgm1, Timp1, A2m, Vim, Cd44, Fcgr2b,* and *Gfap* (**Fig. 2j**). Other notable astrocyte enriched genes upregulated in AB cells included numerous serine protease inhibitors *Serpina3-i,m,h,n*, the nucleoside transporter *Slc43a3* which is involved in purine salvage and metabolism, cytoskeletal organizing adaptor protein *Sorbs2,* tight junction regulator *Mpdz,* and the monocarboxylate transporter *Slc16a1,* among others (**Supplementary Fig. 6**). Some of these DEGs were also upregulated in the AAV/MC condition but were consistently at lower magnitudes compared to the ABC Tool. GO-BPs identified from this list of genes included processes related to cytoskeletal and ECM organization as well as inflammation and cytokine signaling, microbial defense, and wound healing (**Fig. 2i**).

Of the 1659 genes identified to be uniquely upregulated in AB cells (*AB only genes*), some were found on the cARG (23 out of 170) and pARG (40 of 259) lists, including molecules already established to be newly expressed in astrocyte border cells following CNS injury or biomaterial implantation like *Nes* and *Lglas3*^24^. GO-BPs identified from the AB cell only gene list prominently featured numerous processes related to ciliogenesis, including both primary and motile cilium formation, which are hallmarks of G0 state quiescence and barrier functions respectively^29^ (**Fig. 2i**). Upregulated ciliogenesis genes included centrosomal molecules (*C2cd3*, *Cep55*, *Cep70*, *Cep131, Cep162, Cep192, Cep250, Cep295, Cep350, Cenpf, Spice1, Incenp*), cilia and flagella related molecules (Cfap46, Cfap54, Cfap126, *Clxn, Ccdc66, Ccdc74a, Ccdc77, Ccdc88c, Ccdc146, Ccdc191, Kif9, Spef2*), and axonemal dyneins (*Dnai4, Dnah5, Dnah6, Dnah9*), while certain transcriptional regulators of G0 state quiescence and proliferation arrest (*Ccn5, Foxn3, Rbl1, Atf3*) were also uniquely upregulated in AB cells (**Fig. 2j**). AB cells also uniquely expressed high levels of certain pro-inflammatory (*Sucnr1, Klk6, Pycard*) and anti-inflammatory regulators (*Gpnmb, Steap4),* toll-like receptor signaling molecules *(Tnip2, Lgals9, Tlr2, Tlr4, Rnf216),* molecules regulating interferon production or induced by interferon signaling^46^ *(Irf7, Relb, Isg15, Il21, Setd2, Bst2, Ifi47, Ifi208, Oas1a, Oas1h, Oasl1),* ECM remodeling *(Ecm2, Aebp1, Col2a1, Col12a1, Matn4*), and canonical brain barrier transporters^47^ (*Slco1a5, Slc22a2, Slc22a5*) (**Fig. 2j**). Protein expression changes in example downregulated (Kirrel2) and upregulated molecules (Gfap and Lgals3) in AB cells relative to healthy astrocytes were corroborated with IHC (**Fig. 2k**).

These data show that the ABC Tool enables more effective characterization of the AB cell translatome at implanted biomaterials than methods involving pre-labeling of local fields of astrocytes. The improved specificity in AB cell analysis using the ABC tool enabled identification of many hundreds of genes uniquely regulated in AB cells that would otherwise go undetected due to contamination from distinct astrocyte populations in the adjacent neuropil. Furthermore, the observed AB cell translatome at 28 dpi shows a profile consistent with a major re-prioritization of core functions in astrocytes upon adopting border states, with the expression of many mature and region-specific neural circuit-maintaining molecules diminished, so that new border specific functions can be adopted and maintained.

### Astrocyte border formation involves temporally regulated adaptive reprogramming that contracts MC biomaterials

Astrocytes undergo temporally dependent transcriptional changes after CNS injury to form borders around lesions to protect adjacent viable neural tissue, but whether similar reprogramming processes are conserved at injected biomaterials is unknown^29^. To address this knowledge gap, we first assessed the changes in astrocyte morphology at multiple timepoints during border state transitions at injected MC biomaterials and characterized AAV delivery and RiboTag-HA expression timelines to establish the interval in which the ABC Tool could be used for subsequent translational profiling. We performed IHC evaluations on striatal tissue sections taken at 4, 7, 14, 28, and 70 dpi of ABC Tool formulations to capture a range of acute and chronic timepoints (**Fig. 3**). At 4 dpi, uniformly spherical deposits of MC biomaterials in the center of the striatum were surrounded by preserved neural tissue containing fields of hypertrophic but non-overlapping, single domain Gfap- and Vimentin-positive astrocytes. Astrocytes surrounding the biomaterial showed negligible HA-RiboTag expression at 4dpi (**Fig. 3a-b**). By 7 dpi, Vimentin-positive astrocytes had undergone pronounced morphological transformations including territorial area reduction, elongation, and re-organization into extensively overlapping, higher density populations confined to a tissue zone located within 100-200µm of the biomaterial surface (**Fig. 3b, g-i, Supplementary Fig. 7**). Most, but not all, border forming astrocytes expressed HA-RiboTag by 7 dpi, while total and spatial distribution of HA-RiboTag expression as measured by IHC was not significantly altered in AB cells from 7 to 70 dpi, suggesting robust and stable expression of Rpl22-HA in these cells (**Fig. 3b-c, Supplementary Fig. 7**). The amount of astrocyte specific mRNA recovered by RiboTag IP increased from 7 days (∼20ng) to 28 days (∼50ng) consistent with enhanced incorporation of the Rpl22-HA into actively translating ribosomes over that time, but there was no difference in total recovered mRNA by RiboTag IP for 28 and 70 dpi samples suggesting the presence of a stable total number of HA-positive cells at chronic astrocyte borders (**Fig. 3d**).

**Fig. 3.**
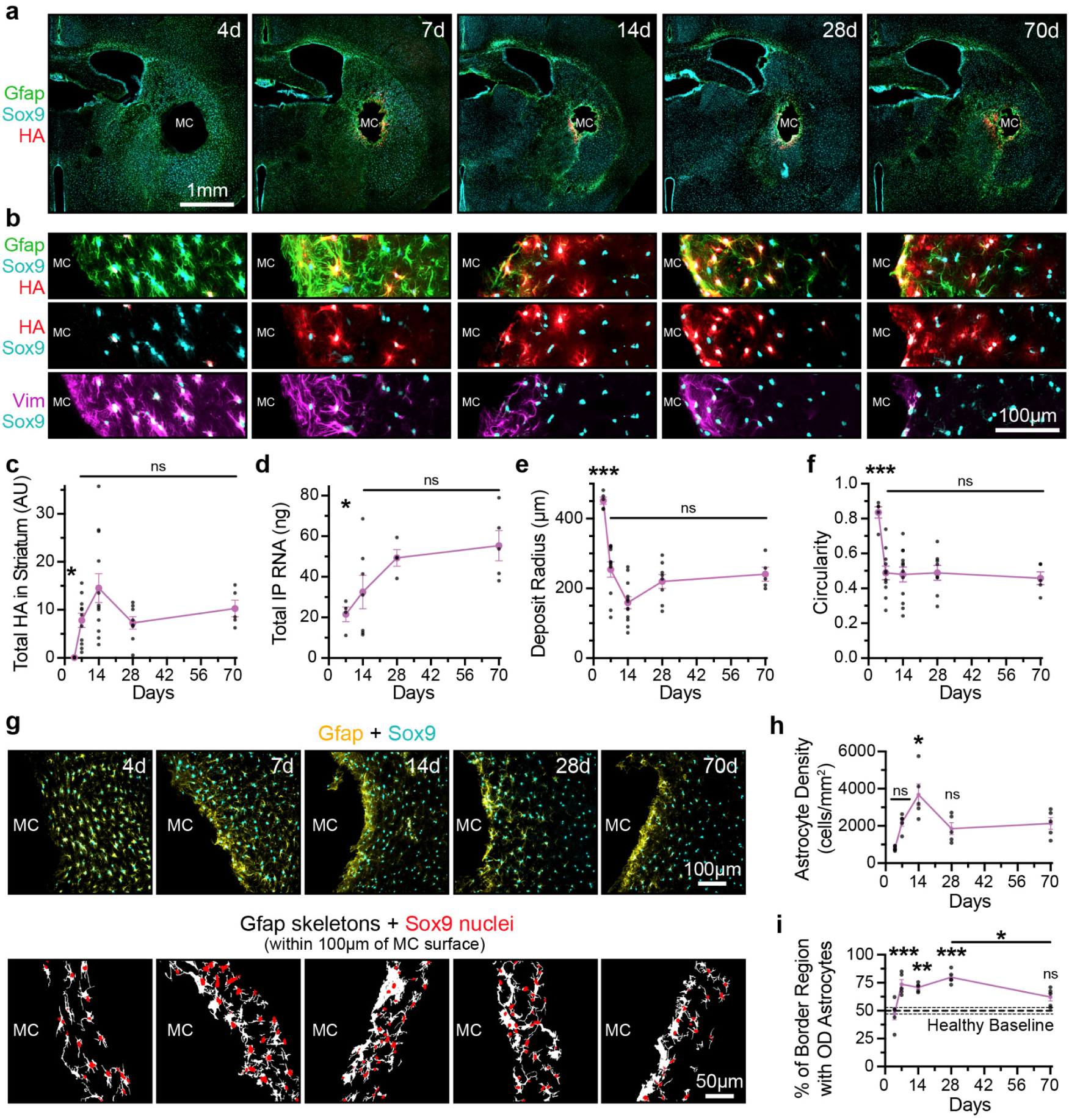
MC biomaterial injections provoke temporally dependent changes in astrocyte morphology and anisotropic contraction of deposits. **a.-b.** Overview (**a.**) and detail (**b.**) images showing changes in astrocyte morphology and HA expression at MC biomaterials from 4-70 dpi. **c.** Quantification of total HA expressed by astrocytes at biomaterials from 4-70 dpi. **d.** Total RNA recovered by RiboTag-IP from 4-70 dpi. **e.-f**. Quantification of change in biomaterial radius (**e.**) and circularity (**f.**) from 4-70 dpi. **g.** Representative images of biomaterial tissue interfaces from 4-70dpi and examples of extracted astrocyte skeletons used for border morphology and cell density analysis. **h.** Quantification of Sox9-positive astrocyte density up to 100µm from the MC biomaterial surface from 4-70 dpi. **i.** Quantification of proportion of overlapping-domain (OD) astrocytes at MC biomaterials from 4-70 dpi. Dashed line shows mean ± s.e.m. for Healthy Baseline (AAV in solution control at 28d, n=7). All x-y plots in this figure (**c.-f., h.-i.**) show not significant (ns), *P < 0.04, **P < 0.003, and ***P < 0.0008 compared to 70d or as designated, except for (**i.**), which is compared to 4d via either a Brown-Forsythe and Welch ANOVA with a Dunnett’s T3 multiple comparisons test (**c.**) or a one-way ANOVA with Tukey’s multiple comparison test (**d.-f., h.-i.**). Graph shows mean ± s.e.m. with individual data points showing: (**c., e.-f.**) 4d (n = 5), 7d (n = 11), 14d (n = 12), 28d (n = 8), and 70d (n = 5). (**d.**) 7d (n = 4), 14d (n = 7), 28d (n = 4), and 70d (n = 5). (**h.-i.**) n=5 for all timepoints.

Wound closure and lesion contraction are critical functions of astrocyte borders formed at CNS injuries. This process is driven by upregulation of *Gfap* and *Vim* and numerous other cytoskeletal remodeling genes involved in epithelial-to-mesenchymal transition (EMT) including *Acta2* and *Myh9,* which enable astrocytes to generate traction forces to contract the margins of lesion cores^29,48–50^. In line with these gain-of-functions, we noted marked volume contraction of MC biomaterials between 4 and 7 dpi, coinciding with the period of most dynamic change in astrocyte morphologies around the biomaterial deposit. During this critical 3-day interval, MC biomaterials showed a significant reduction in average deposit area (∼0.8 to 0.35mm^2^), radius (451 to 254 µm), and circularity (0.76 to 0.47), consistent with anisotropic contraction (**Fig. 3a-b,e-f, Supplementary Fig. 7**). Notably, there was no further change in biomaterial geometry after 7dpi, with deposits persisting essentially unaltered through 70dpi, consistent with contractile processes being acutely executed and MC biomaterials showing negligible biodegradation.

Given HA-positive astrocytes were detected as early as 7 dpi and persisted through 70 dpi, we next performed translational profiling of AB cells using the ABC Tool at 7, 14, 28, and 70 dpi, comparing these samples to healthy striatal astrocytes using the AAV solution control as before. All RiboTag IP samples showed consistent FPKM levels and extent of enrichment for the virally expressed 3xHA transcripts, indicating effective AB cell mRNA isolation across the different timepoints (**Supplementary Fig. 8**). To identify the temporally regulated gene expression profiles in AB cells we performed PCA on 6,981 DEGs that were detected as differentially regulated for at least one of the timepoints compared to healthy striatal astrocytes (**Fig. 4a, Supplementary Fig. 8**). PCA on this dataset revealed a clear temporal progression in AB cells with PC1 indicating acute and transient changes in a manner similar to the temporally dependent changes previously noted in astrocytes responding to CNS injury^29^.

**Fig. 4.**
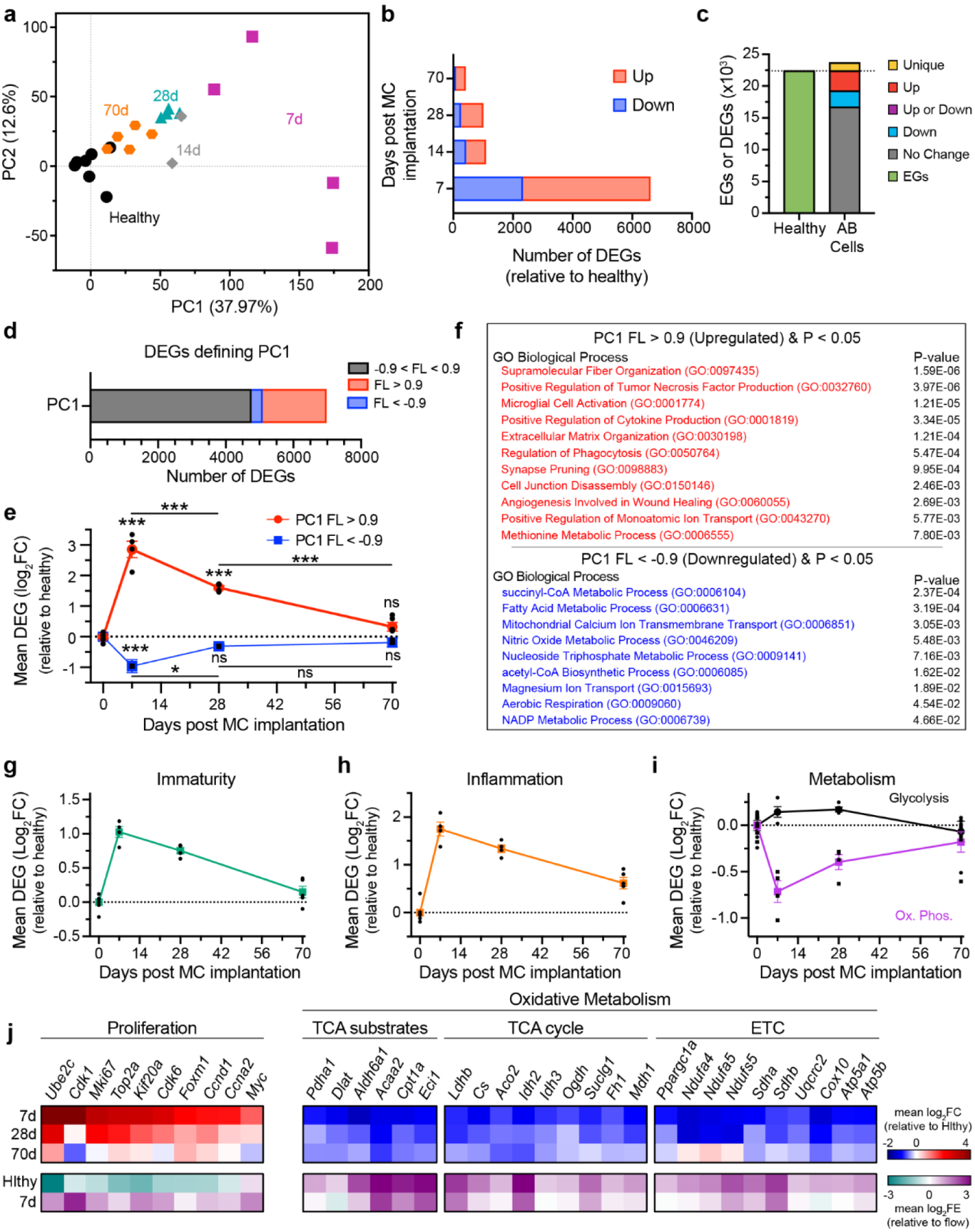
Astrocyte border formation involves temporally regulated transcriptional reprogramming. **a.** PCA of all temporally regulated DEGs. **b.** Number of DEGs at each post biomaterial injection timepoint relative to healthy astrocytes (AAV IP). **c.** Number of genes regulated in healthy astrocytes compared to AB cells at all timepoints. **d.** Number of DEGs contributing to defining transiently upregulated (PC1 FL > 0.9) and downregulated DEGs (PC1 FL < −0.9). **e.** Mean log_2_(FC) of genes defining transiently upregulated (PC1 FL > 0.9) and downregulated DEGs (PC1 FL < −0.9) at different times after MC biomaterial injection. Plot shows not significant (ns), *P < 0.03, and ***P < 0.0005 compared to 0d or as designated via one-way ANOVA with Tukey’s multiple comparison test. **f.** GO-BPs significantly upregulated (red) or downregulated (blue) as identified by unbiased evaluations of DEGs defined by PC1 FL > 0.9 or FL < −0.9. **g.-i.** Mean log_2_FC at different times after MC biomaterial injection for curated list of genes defining immaturity (**g.**), inflammation (**h.**), as well as glycolysis and oxidative phosphorylation (Ox. Phos.) metabolism (**i.**). **j.** Heatmaps of log_2_FC and log_2_FE for proliferation and metabolism-related genes at designated timepoints post MC biomaterial injection. (**e., g.-i.**) show mean ± s.e.m. with individual data points showing Healthy (n = 7), 7d (n = 4), 28d (n = 4), and 70d (n = 5).

However, unlike those prior CNS injury evaluations^29^, PC2 did not reveal any significant persistent gene expression changes across the 70-day time course but rather accounted for inter-sample variability defined by the 7 dpi samples. The transient nature of the gene regulation post biomaterial injection was also reflected in the total number of DEGs detected across the different timepoints, with 6,610 genes differentially regulated at 7 dpi but less than 430 genes altered at 70 dpi compared to healthy striatal astrocytes (**Fig. 4b**). Of the total number of detected healthy astrocyte expressed genes (EGs), nearly 75% (16,676) were not significantly changed at any time during AB cell formation, 3,164 were upregulated, 2,536 were downregulated, and 1,273 were defined as newly expressed genes, since they were not produced at detectable levels in healthy astrocytes (**Fig. 4c**). Of the newly expressed genes only 20 (1.6%) maintained elevated expression through 70dpi but included chemokines *Ccl3*, *Ccl9*, the cytokine inactivator *Mmp12*, and established astrocyte reactivity transcriptional regulator *Runx1*^30^.

Over 2,200 genes (∼32% of all DEGs) had a PC1 FL >0.9 which indicated prominent transient upregulation that peaked at 7dpi (**Fig. 4d-e**). There were nearly 6-fold more DEGs with a FL >0.9 compared to FL <-0.9, which represented genes transiently downregulated. This is consistent with there being more temporally dependent gain-of-function changes than loss-of-function changes associated with AB formation (**Fig. 4d-e**). Unsupervised analysis of GO-BPs for transiently upregulated DEGs (PC1 FL>0.9) revealed processes associated with the activation of inflammation (cytokine production, phagocytosis, and microglia activation), ECM and supramolecular fiber organization, angiogenesis, synapse pruning, ion transport, and methionine-related metabolism (**Fig. 4f**). By contrast, transiently downregulated DEGs (PC1 FL<-0.9) were associated with numerous metabolic processes including aerobic respiration, NADP reactions, acetyl-CoA and succinyl CoA synthesis, fatty acid metabolism, and nucleoside triphosphate metabolism among others (**Fig. 4f**).

Analysis of curated astrocyte- and biological process-specific genes lists revealed notable temporal dependent changes in AB cells (**Fig. 4g-j**). In a manner similar to CNS injury responses, astrocytes responding to MC biomaterial injections transiently upregulated astrocyte immaturity genes including *Ube2c, Haspin, E2f7, Bub1,* and *Foxd3* while concurrently downregulating astrocyte maturity genes such as *Kcnj14*, *Id4*, *Ntm*, *Gpc5,* and *Slc7a10* as well as many cAEGs (**Fig. 4g, Supplementary Fig. 8-9**). Canonical genes regulating proliferation including *Cdk1*, *Mki67*, *Top2a*, *Kif20a*, *Ccnd1* were also upregulated acutely and peaked at 7dpi (**Fig. 4j**, **Supplementary Fig. 8**). For both immaturity and proliferative gene programs, there was a 2-fold or greater increase in mean differential expression at 7 dpi but a return to healthy baseline levels by 70dpi. The peak of proliferative gene expression likely occurred sometime before the earliest transcriptomically evaluable 7dpi timepoint given that histological staining showed numerous Top2a- and Sox9-positive astrocytes at forming astrocyte borders at 4dpi but relatively fewer proliferative cells at 7dpi (**Supplementary Fig. 8**). AB cells also showed a more than 6-fold upregulation of cARGs (**Supplementary Fig. 8**) and 2.5-fold upregulation of inflammation genes at 7 dpi (**Fig. 4h**). The expression of these reactive programs decreased steadily after 7dpi but, unlike the immaturity and proliferative genes, did not fully return to baseline healthy levels at 70dpi. Evaluation of the FLVCA analysis derived gene lists from before also showed temporally dependent regulation, with *Astrocyte genes Down in AB cells* being the most depressed at 7dpi while mean DEGs levels for *Astrocyte genes Up in AB* cells and *AB only genes* peaking at 7dpi (**Supplementary Fig. 8**). Given the noted MC biomaterial contraction from 4 to 7 dpi by IHC, we also evaluated changes in expression of genes associated with EMT, since it is a major driver of contractile cell phenotypes necessary for tissue wound closure. Prototypical EMT genes, including *Vim*, *Fn1*, *Myh9, Cd44, Cnn1, Timp1, Ecm1, Emp3, Fmod,* and *Tgm2*, were all upregulated over 2-fold by 7 dpi and mostly persisted at these elevated levels through 28dpi, whereas other canonical EMT genes like *Acta2*, *Tagln*, *Postn*, and *Snai1*, were not significantly differentially regulated at any post-injection timepoints (**Supplementary Fig. 9**).

Unsupervised GO analysis applied to PC1-defining genes revealed that astrocytes undergo several pronounced, temporally regulated metabolic changes in response to MC biomaterial injections (**Fig. 4f**). This prompted us to conduct further hypothesis-driven analysis of metabolism changes by evaluating curated hallmark gene sets for glycolysis and oxidative phosphorylation taken from the Molecular Signatures Database (MSigDB). These metabolic programs are known to be reciprocally regulated and altered during proliferative, reactive, and contractile responses. AB cells showed a modest increase in expression of glycolysis genes including master transcription factors *Myc* and *Foxm1* which were upregulated by over 4- and 2-fold respectively at 7dpi (**Fig. 4i-j**). These transcriptional regulators are key drivers of aerobic glycolysis metabolism which is essential for fueling cell proliferation. Genes associated with oxidative phosphorylation and oxidative metabolic processes including TCA substrate production (*Pdha1, Dlat, Acaa2,* and others), TCA cycle (*Ldhb, Cs, Idh2, Fh1*, and others), and ETC (*Ppargc1a, Ndufa4, Sdha*, and others) were comprehensively downregulated acutely at 7dpi and failed to fully recover to pre-injection levels in the AB cells profiled at 70dpi (**Fig. 4i-j**). In healthy striatal tissue, oxidative metabolism genes are highly enriched in RiboTagIP samples compared to flow-through, indicating their contribution to essential and specialized metabolic functions present more so in astrocytes than other cell types (**Fig. 4j**). There was significant de-enrichment of these oxidative metabolism genes following MC biomaterial injection, consistent with a loss of specialized astrocyte metabolism during AB cell state transition.

The nature and magnitude of the temporally regulated transcriptional reprogramming associated with AB cell formation at injected MC biomaterial mimicked astrocyte wound responses to ischemic strokes made in the same striatal location by focal L-N5-(1-Iminoethyl)ornithine (LNIO) injection (**Supplementary Fig. 10**). Following stroke, much like upon MC biomaterial injection, astrocytes underwent transient dedifferentiation, short-lived increased expression of proliferation and immaturity genes, diminished oxidative metabolism, and activation of inflammation and astrocyte reactivity with maximally altered gene expression detected at 2 days post injury. Across all regulated gene programs examined, the magnitude of differential expression and temporal dynamics of these expression changes in astrocytes were essentially equivalent in stroke versus MC biomaterials, such that the total number of DEGs detected at 7- (6283) and 28- (1243) days post stroke were comparable to the magnitude of changes detected at MC biomaterials at the same timepoints (6610 at 7dpi and 1007 at 28dpi) (**Supplementary Fig. 10**).

These data show that AB cell state transitions stimulated by MC biomaterial injections involve temporally regulated transcriptional reprogramming in local astrocytes, resulting in transient periods of dedifferentiation, immaturity, proliferation, and inflammation regulation as well as activation of EMT leading to marked contraction of biomaterial deposits. To support these essential border forming processes, AB cells remodel their metabolism by upregulating glycolysis and concurrently reducing highly astrocyte-specialized oxidative processes. Biomaterial FBR-induced transcriptional reprogramming mimics the essential facets of astrocyte wound responses to ischemic and hemorrhagic injuries, suggesting that the processes involved in astrocyte border formation are biologically conserved and essential to restoring critical barrier functions at any localized tissue perturbation.

### Chronic astrocyte border cells persist at MC biomaterials with progressive immunoregulatory gain-of-function

To identify features of AB cells that might emerge or consolidate after the acute period of astrocyte state transition, we examined DEGs regulated after 7 dpi but persisting through 70 dpi. Of the 6610 DEGs detected at 7dpi, 40% of genes (2671) were regulated back towards pre-injection healthy astrocyte levels (1764 downregulated from 7dpi and 907 upregulated from 7dpi). There were 793 genes regulated from 7-70dpi that were not changed from 0-7dpi, making them delayed regulated genes (**Fig. 5a**). Of the 3464 total genes that were altered from 7 to 70 dpi, there were only 18 DEGs detected when making comparisons between the 28 and 70 dpi timepoints, suggesting these later two timepoints contain AB cells that were essentially transcriptionally equivalent (**Fig. 5a**). Based on this insight, we pooled the 28 and 70 dpi samples for a subsequent analysis, designating the pooled group as chronic AB cells. Upon reanalyzing differentially expressed genes relative to healthy we detected 6648 total DEGs across the acute (7 dpi) and chronic AB cell designations. Of those DEGs, 5745 (∼ 86%) genes exhibited transient regulation (either up or downregulated acutely and returning to baseline levels chronically). The remaining 903 DEGs were chronically altered by being persistently up- or down-regulated from acute to chronic timepoints (795 DEGs) or by being regulated in a delayed manner in the chronic state (108 DEGs) (**Fig. 5b**). Of the 903 chronically regulated genes, 648 (72%) continued to be upregulated above healthy levels, maintaining expression levels that were greater than 3-fold higher on average than the healthy astrocyte baseline at 70dpi. By contrast, 255 genes (28% of chronic genes) persisted as downregulated DEGs and on average had approximately a 2-fold lower level of expression than healthy astrocytes (**Fig. 5c**). Notably, 477 of 903 (53%) of the chronically regulated genes were found on the gene lists derived from the prior FLVCA analysis applied to the 28dpi samples, including 198 DEGs from the *Astrocyte genes Up in AB cells* list, 181 DEGs from the *AB only genes* list, and 98 DEGs from the *Astrocyte genes Down in AB cells* list (**Supplementary Table 3-4**). Furthermore, chronically upregulated DEGs accounted for 74 of 259 pARGs that are chronically elevated and highly enriched in astrocyte borders at CNS injuries^29^ (**Supplementary Table 5**).

**Fig. 5.**
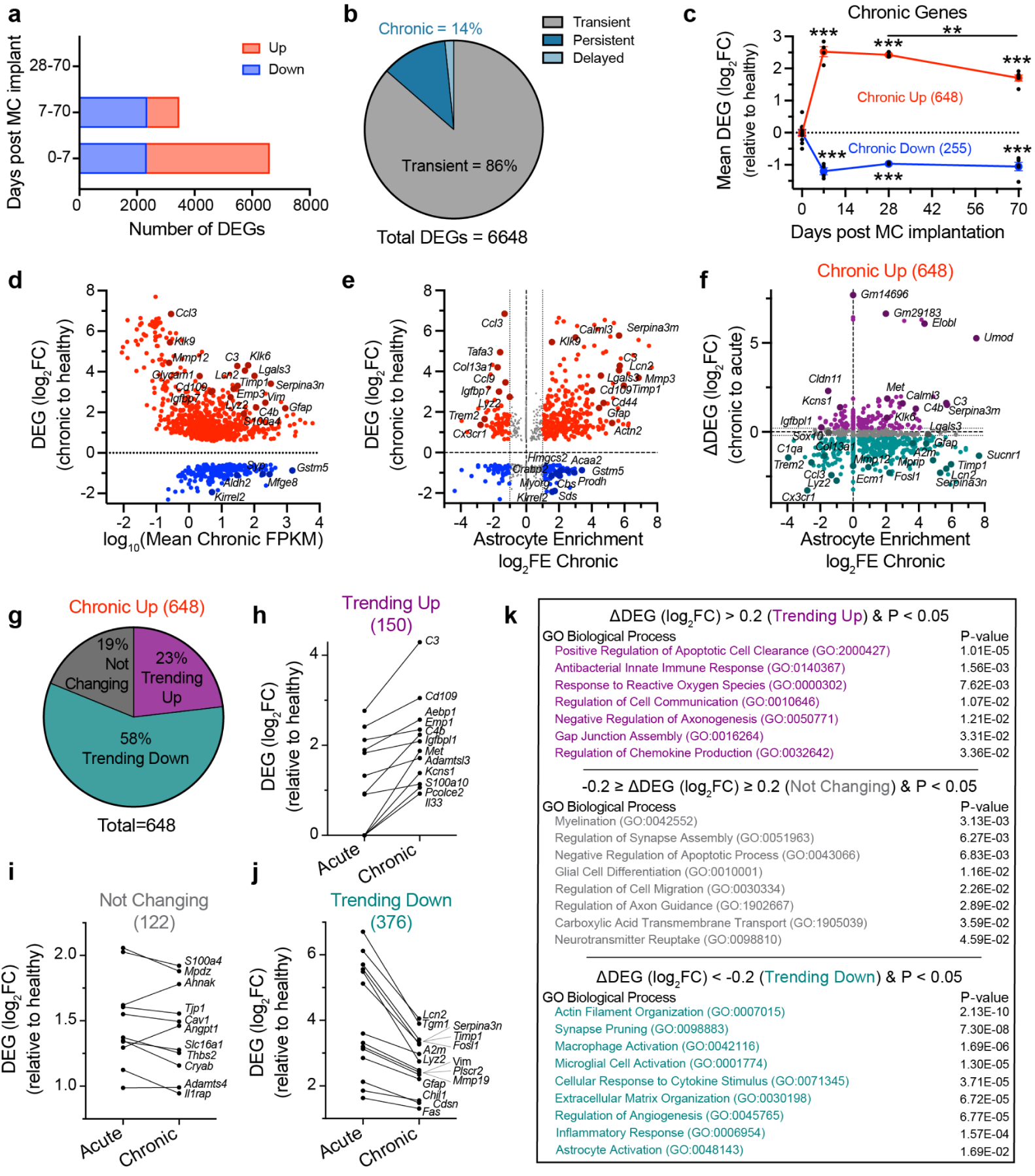
Chronic astrocyte border cells at MC biomaterials persist with increasing gain of functions in host defense and immune regulation. **a.** Number of astrocyte DEGs significantly changed between different timepoints. **b.** Relative proportion of transiently and chronically regulated DEGs in AB cells. **c.** Mean log_2_(FC) of genes defining chronically upregulated and downregulated DEGs. Graph shows mean ± s.e.m. with individual data points showing Healthy (n = 7), 7d (n = 4), 28d (n = 4), and 70d (n = 5). Plot shows **P < 0.004 and ***P < 0.0001 compared to 0d or as designated via one-way ANOVA with Tukey’s multiple comparison test. Comparisons of 7d to 28d for Chronic Up, and both 7d to 28d and 28d to 70d for Chronic Down are not significant. **d.** Scatter plot showing mean log_2_FC and FPKM of 903 chronically regulated DEGs. Upregulated genes are colored in red and downregulated genes are colored in blue. Genes that are of special interest are labeled. **e.** Scatter plot showing comparison of relative enrichment and expression levels of significant chronically regulated DEGs. Upregulated genes are colored in red and downregulated genes are colored in blue. Genes with minimal astrocyte enrichment or de-enrichment (−1 ≤ log_2_FE ≤ 1) are labeled in gray. Genes that are of special interest are labeled. **f.** Scatterplot showing change in differential expression between acute and chronic timepoints (ΔDEG), and log_2_FE values for significant, chronically upregulated genes. Genes that are trending up (ΔDEG > 0.2) are colored in purple, genes trending down (ΔDEG < −0.2) are colored in turquoise, and genes that are not changing (−0.2 < ΔDEG < 0.2) are colored in gray. Genes that are of special interest are labeled. **g.** Relative proportion of chronically upregulated genes with ΔDEG that is trending up, down, or not changing. **h.-j.** Mean DEG values at acute and chronic timepoints for select genes that are **h.** trending up, **i.** not changing, and **j.** trending down. **k.** Table showing significant GO-BPs associated with upregulated genes categorized according to ΔDEGs that are trending up, down, or not changing.

Many chronically regulated DEGs were uniquely expressed by AB cells, with 45% of all chronically upregulated DEGs being enriched by more than two-fold in these astrocytes compared to other cell types (**Fig. 5d-e**). There were also many prominent astrocyte enriched genes persistently downregulated in chronic borders, including molecules involved in fatty acid metabolism and transport (*Elovl2*, *Fads1, Crabp2, Mfsd2a, Acaa2, Acot11*), amino acid metabolism *(Sds*, *Cbs*, *Ivd, Prodh, Gldc, Sardh*), ketone body biosynthesis (*Hmgcs2*), and established astrocyte-specific molecules like the glycoside hydrolase *Myorg,* and the growth factor receptor *Fgfr3*. Collectively, the chronically downregulated gene expression data suggest certain metabolic processes, in particular how astrocytes use and produce lipid-based oxidative substrates, are persistently dysregulated in AB cells.

For the 648 chronically upregulated DEGs, we compared the change in gene expression during the acute (7dpi) to chronic transition (28/70dpi) to identify three distinct gene classifications based on the nature of the change: (i) genes trending up (150 genes), genes not changing (122 genes), and genes trending down (376 genes) (**Fig. 5f-k**). Upregulated genes trending down represented the majority (58%) of chronically upregulated genes and GO-BP analysis of these genes revealed many of the same processes deemed to be transiently upregulated in the prior analysis including synapse pruning, ECM organization, angiogenesis, and inflammatory responses (**Fig. 5k**). Molecules assigned into this category included canonical, astrocyte-enriched reactivity genes such as *Tgm1*, *Timp1*, *Lcn2*, *Serpina3n, Cd44*, and *A2m*, which all contribute immune regulatory functions. Other immune regulating molecules chronically elevated but trending down over time in AB cells included *Trem2*, *Ccl3*, *C1qa*, *Cxcr1* and *Lyz2*, but these were not enriched in AB cells relative to the other cell types present around the implanted MC biomaterials that were processed in the flow-through samples, likely because recruited immune cells and microglia also highly express these molecules (**Fig. 5e-f,j**). Upregulated and astrocyte enriched genes that persisted essentially unchanged during the acute to chronic transition included molecules involved in formation and maintenance of tight junctions and barrier functions that all happen to be regulated by the transcription factor Runx1 (*Angpt1, Tjp1, Mpdz, Lgals3*) as well as transporters for monocarboxylates, ergothioneine, and urea (*Slc16a1, Slc22a4, Slc14a1*) among others (**Fig. 5i**).

GO-BP analysis of astrocyte enriched genes that trended up chronically revealed processes critical to antibacterial defense, apoptotic cell clearance, response to reactive oxygen species, and regulation of chemokine production (**Fig. 5k**). Molecules in this categorization included the antimicrobial defense factor, *C3*, which increased its expression nearly 3-fold during the acute to chronic transition in AB cells to result in a 20-fold increase in expression compared to healthy astrocytes (**Fig. 5h**). It is notable that a similar chronic increase in *C3* expression is observed as part of astrocyte wound responses to CNS injuries^29^. Expression of other complement system molecules, *C4a*, C*4b,* and *Serping1*, which coordinate opsonization processes in conjunction with *C3*, were also detected in the trending up gene list, consistent with a more prominent role chronically as part of ongoing immune surveillance at the astrocyte border. Other notable immunoregulatory molecules involved in host defense that showed increased expression in chronic AB cells included innate antimicrobial effectors (*Trl4*, *Ly96*, *Umod*, *Trf*, *Il33*, and *Serpinb1a*), antiviral responses (*Plscr1, Stx11*), and immune modulatory factors (*Cd109, Emp1, Igfbpl1, Adamtsl3, Klk6* and *Klk9*) (**Fig. 5h**).

Collectively, the translational profiling of chronic AB cells shows that only a small subset of DEGs persist in AB cells beyond the acute transition phase, but these molecules impart gain-of-function in astrocyte borders, most notably in immune regulation and host defense capabilities, that are likely vital to protecting spared neural tissue adjacent to the implanted biomaterial.

### Delivery of ablation molecules from injected biomaterials does not prevent astrocyte border formation and exacerbates immune recruitment and fibrosis

A potential strategy to improve the long-term performance of implanted biomaterials is to attenuate the astrocyte border component of the FBR by either minimizing the number of locally recruited astrocytes or modulating their phenotypic and functional state transitions. Ablation of wound repair astrocyte responses at CNS injury lesions via transgenic loss of function manipulations effectively reduces the cellular density and thickness of astrocyte borders but is coupled with increased inflammation and fibrotic lesion volume expansion^20,30,51,52^. To test how disrupting astrocyte border formation alters the FBR to MC biomaterials and the capacity to characterize AB cells, we evaluated delivering two distinct ablation molecules from the ABC Tool: (i) the vasoconstrictive agent, LNIO, which induces focal ischemia and ablates neurons and glia indiscriminately (**Fig. 6a, Supplementary Fig. 11**)^24,53^; and (ii) an astrocyte-specific glia toxin, L-2-Aminoadipic Acid (L2AA), which promotes astrocyte-selective ablation with minimal neuron or microglia loss by acting as a competitive inhibitor of glutamate transporters, which are highly and uniquely expressed by astrocytes^54–57^ (**Fig. 6a, Supplementary Fig. 11-12**). The effects of both molecules on striatal neural tissue in the mouse brain were confirmed through solution injections into a sperate cohort of mice prior to incorporation into biomaterials (**Supplementary Fig. 11**).

**Fig. 6.**
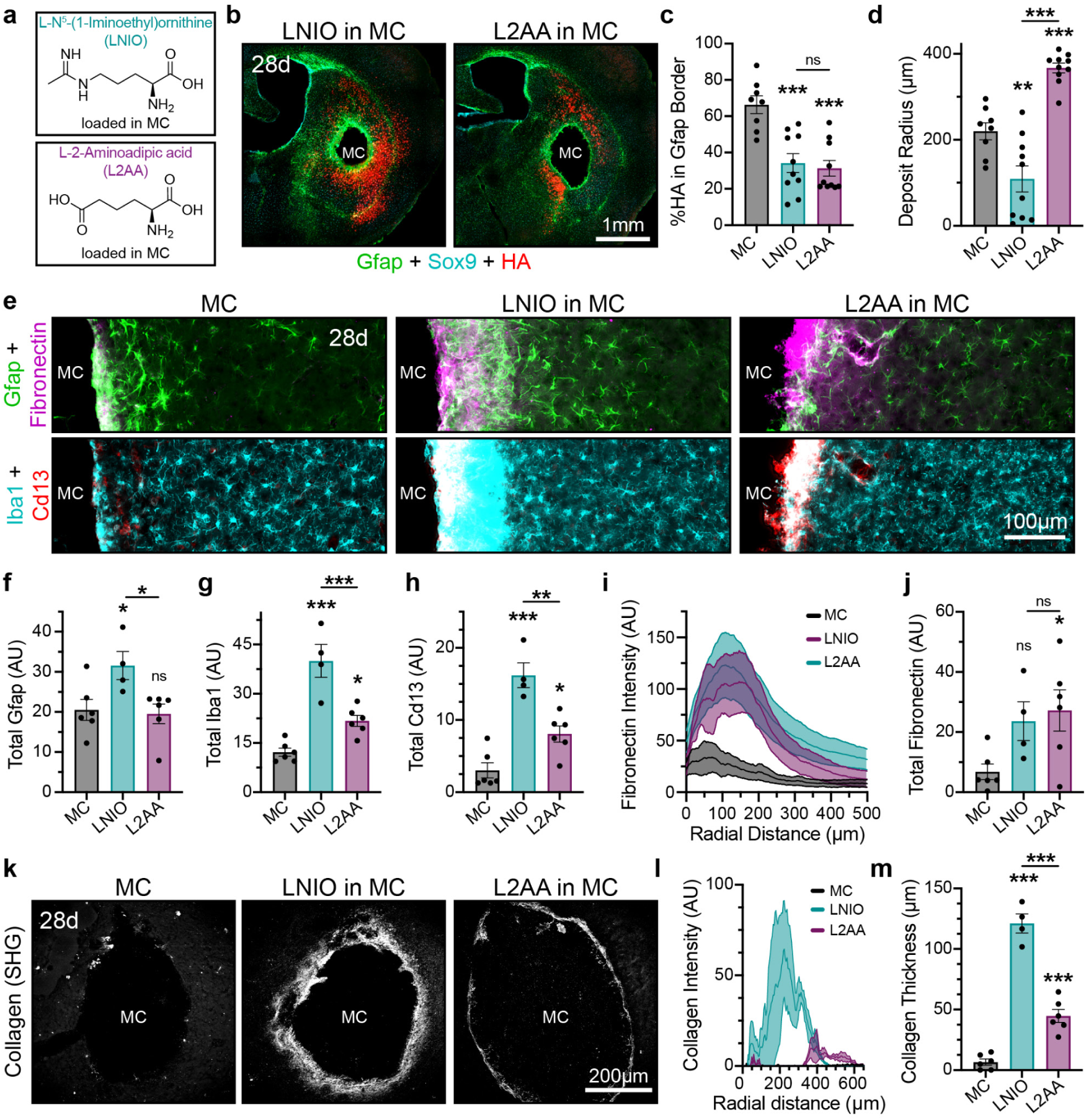
Delivery of ablation molecules from injected biomaterials temporarily disrupts astrocyte border formation leading to altered RiboTag AAV delivery and FBR exacerbation. **a.** Schematic showing chemical structures of L-N^5^-(1-Iminioethyl)ornithine (LNIO) and L-2-Aminoadipic Acid (L2AA). **b.** Overview images showing morphology of astrocyte borders and distribution of HA expression in striatal neuropil at 28 dpi for LNIO in MC and L2AA in MC. **c.** Percent of total HA expression contained within Gfap-positive astrocyte borders for MC biomaterials formulated with and without ablation molecules. **d.** Quantification of MC biomaterial deposit radius at 28 dpi for formulations with and without ablation molecules. **e.** Detail images of the FBR response to MC biomaterials formulated with and without ablation molecules showing stromal (Fibronectin) and inflammatory (Iba-1, Cd13) cells. **f.-h.** Quantification of total Gfap (**f.**), Iba1 (**g.**), and Cd13 cells (**h.**) within 300µm of the biomaterial-tissue interface. **i.** Fibronectin intensity measured radially from the deposit surface. **j.** Quantification of total Fibronectin within 300µm of the biomaterial-tissue interface. **k.** Overview images showing extent of fibrillar collagen deposition as measured by SHG microscopy. **l.** Collagen intensity measured radially from the center of injection. **m.** Quantification of fibrotic capsule thickness for MC biomaterials formulated with and without ablation molecules. All bar charts in this figure show not significant (ns), *P < 0.05, **P < 0.006, and ***P < 0.0009 compared to MC or as designated via one-way ANOVA with Tukey’s multiple comparison test. Plots **c.-d.** in this figure show mean ± s.e.m. with individual data points showing MC (n = 8), LNIO (n = 10), and L2AA (n = 10). Plots **f.-h.**, **j.**, and **m.** show mean ± s.e.m. with individual data points showing MC (n = 6), LNIO (n = 4), and L2AA (n = 6). Intensity traces in **i** and **l** show mean ± s.e.m., with s.e.m. represented as light shaded areas.

Local delivery of LNIO and L2AA from injected MC biomaterials disrupted a narrow tissue region immediately adjacent to the biomaterial surface, leading to more severe FBRs, altered astrocyte borders, and changes to the spatial distribution of viral transduction as indicated by HA expression (**Fig. 6**, **Supplementary Fig. 13**). Comparing mouse behavior prior to biomaterial injection and 3 dpi by cylinder test showed no significant change in either activity levels or motor performance for the LNIO- or L2AA-loaded MC biomaterial treated mice, confirming the localized effects of the molecule-based ablation (**Supplementary Fig. 14**). Delivering LNIO or L2AA from MC biomaterials caused a 50% decrease in the expression of HA in AB cells by IHC, a 75% reduction in mRNA recovered by RiboTag IP, and ∼12-fold fewer sequenced 3xHA transcripts, suggesting that formed astrocyte borders were insufficiently labelled for effective translational profiling (**Fig. 6c, Supplementary Fig. 13**). There was noticeably increased detectable HA-expression in neuropil astrocytes away from biomaterial deposits, with sparse HA-positive cells extending throughout the entire striatum for both LNIO- and L2AA-loaded biomaterials. This is consistent with increased tissue dispersal of delivered RiboTag-AAV as a result of the small molecule-induced tissue disruption, which inhibits astrocyte-based partitioning and subsequent restriction of molecular delivery, in a manner similar to that noted before for biomaterials that provoke equivalent wound-like FBRs^24^. Focally delivering LNIO markedly reduced MC deposit size at 28 dpi, whereas delivering L2AA significantly increased deposit size at the same timepoint relative to MC-only, indicating molecule dependent effects on contractile functions of astrocyte borders (**Fig. 6b,d**). Notably, neither ablation molecule prevented astrocyte border formation around the biomaterial by 28dpi, and in fact, LNIO delivery led to the formation of significantly thicker Gfap-positive borders compared to L2AA or MC-only (**Fig. 6e-f**). Thus, collectively, these data are consistent with small-molecule ablation causing acute and focal loss of astrocytes immediately adjacent to biomaterial deposits that contributes to only a temporary delay in astrocyte border formation and its critical barrier functions, rather than persistently altering these responses.

Molecule-based ablation of astrocyte borders caused increased localized inflammation with significantly greater numbers of Iba-1-positive microglia and Cd13-positive macrophages noted at biomaterials delivering each of the molecules but was markedly more severe for the LNIO formulation (**Fig. 6e, g-h**). Molecule-based ablation of astrocyte borders also resulted in the increased synthesis of collagen and formation of thick fibrotic capsules circumscribing MC biomaterials, which otherwise showed negligible fibrosis at the 28 dpi timepoint (**Fig. 6k**). Quantification of the extent of fibrillar collagen deposition produced by fibronectin-positive stromal cells (**Fig. 6i-j**) using second harmonic generation (SHG) 2-Photon (2P) microscopy^58^ showed that LNIO and L2AA delivery led to fibrotic capsules that were approximately 120µm and 50µm thick respectively, which were more than 17 and 2.6 fold greater than the minimal capsules detected at MC-only formulations (**Fig. 6k-m**).

These findings show that small molecule-based ablation of local astrocytes acting either indiscriminately or selectively on astrocytes merely delays rather than prevent astrocyte border formation at injected MC biomaterials. Notably, the ablation molecule-driven delay in astrocyte border formation exacerbates FBR at MC biomaterials resulting in significantly increased immune recruitment and fibrosis. These data suggest that preventing astrocyte border formation through cell ablation is not an effective strategy for improving the biocompatibility of implanted biomaterials in the CNS.

### β-hydroxybutyrate (BHB) attenuates the metabolic reprogramming essential to astrocyte proliferation *in vitro*

Metabolic reprogramming emerged as a prominent, temporally dependent transcriptional change identified by the ABC tool, with AB cell state transitions requiring transient upregulation of glycolytic pathways and concurrent downregulation of specialized oxidative metabolism to fuel proliferative and reactivity responses (**Fig. 4, 5**)^18,19,59,60^. We hypothesized that delivering small molecule regulators counteracting this metabolic reprogramming would attenuate AB cell state transitions in biomaterial-adjacent astrocytes, preserving their viability and minimizing neural tissue disruption around the implant. We identified β-hydroxybutyrate (BHB), which is a ketone body produced in low glucose states^61^, as a potential candidate molecule to achieve this modulation. BHB is an established alternate energy substrate for astrocytes, promoting increased mitochondrial oxidative metabolism and inhibiting glycolytic energy pathways^61^. Healthy striatal astrocytes express the BHB transporter, Mct1 (*Slc16a1*) (**Supplementary Fig. 15**), and there is robust and selective upregulation of Mct1 in response to MC biomaterial injection (**Supplementary Fig. 15**). BHB is produced endogenously by Hmgcs2, the rate-limiting enzyme responsible for ketogenesis, and serves as a direct substrate for oxidative metabolism by being converted to acetyl-CoA which enters the TCA cycle. Expression of *Hmgcs2*, *Acaa2* (enzyme responsible for the final step in the conversion of BHB to acetyl-CoA), and *Ppargc1a* (a master regulator of BHB metabolism), were all detected as significantly downregulated in striatal astrocytes transitioning into AB cell states by the ABC Tool (**Fig. 4j, 5e**). Thus, we rationalized that delivery of exogenous BHB from MC biomaterials had the potential to rescue oxidative metabolism in AB cells by acting as a metabolic substrate or signaling molecule for mitochondrial biogenesis and other relevant processes^61,62^.

To characterize the effect of exogenous BHB (20mM) on astrocyte responses, we first employed an *in vitro* culture system using mouse quiescent astrocytes (qA), derived by differentiating RiboTag-expressing neural progenitor cells (NPC) with 2% fetal bovine serum (FBS). Consistent with our previous studies^42,63^, there was essentially complete conversion of NPC into astrocytes under FBS exposure, with greater than 84% of cells being Sox9-Hi-expressing qA cells after only 2 days of culture, irrespective of whether normal glucose (Glc) or BHB fuel was used in the media (**Supplementary Fig. 16**). qA cells generated in Glc media exhibited a 2-fold increase in both Gfap and Mct1 protein expression relative to qA derived in BHB media, consistent with increased reactivity with the Glc fuel (**Fig. 7a-b**, **Supplementary Fig. 16**). Cell survival was not apparently altered by fueling the qA with BHB fuel, although there was a small reduction in cell density compared to Glc fed qA after 2 days (**Fig. 7a**). To assess the differential effects of Glc and BHB on astrocyte proliferative responses, we conducted BrdU pulsing experiments after exposing qA to EGF and FGF (E/F) under serum free conditions for four days to stimulate mitosis (**Fig. 7c**). While over 65% of astrocytes grown in Glc + E/F media were BrdU-positive following a 6-hour BrdU (20µM) pulse, proliferative responses were significantly dampened under BHB + E/F media conditions, with only 21% of astrocytes being BrdU-positive (**Fig. 7c-d**). Proliferating astrocytes fed Glc showed increased expression of Gfap compared to the BHB, although both proliferating populations had reduced Gfap expression compared to qA (**Supplementary Fig. 16)**.

**Fig. 7.**
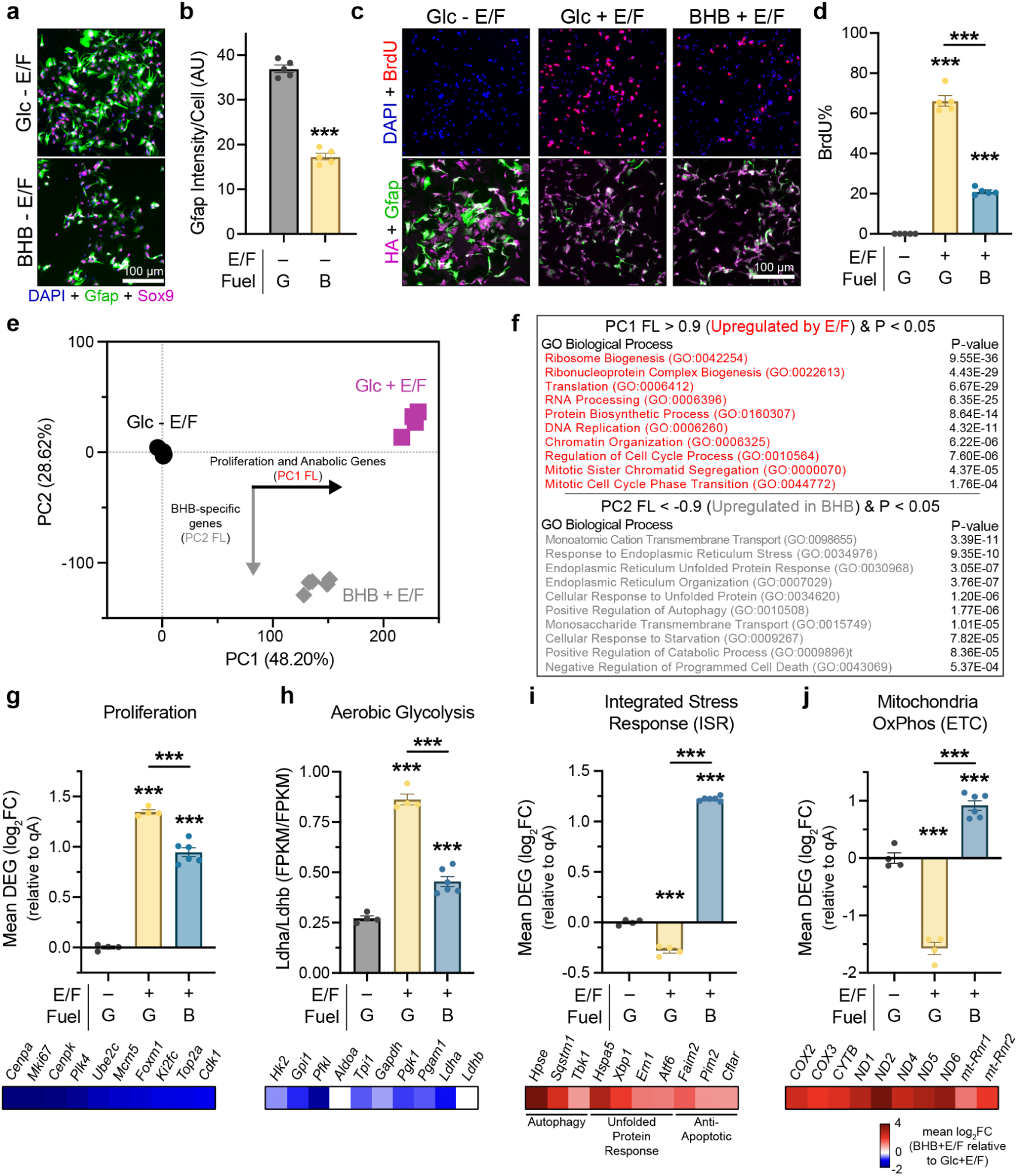
β-hydroxybutyrate (BHB) attenuates the metabolic reprogramming essential to astrocyte proliferation *in vitro.* **a.** ICC images for quiescent astrocytes (qAs) fueled with Glc- or BHB-media. **b.** Quantification of total Gfap intensity per cell for qAs fueled with Glc- or BHB-media. **c.** ICC images for qA and E/F mitogen-stimulated astrocytes in Glc- or BHB-media undergoing 6-hour BrdU pulse showing proliferation (BrdU) and HA RiboTag production. **d.** Quantification of percent BrdU-positive cells for qA and E/F mitogen-stimulated astrocytes in Glc- or BHB-media. **e.** PCA of all DEGs detected as differentially regulated between the 3 *in vitro* conditions. **f.** GO-BPs identified by unbiased evaluations of DEGs defined by PC1 FL > 0.9 (E/F mitogen-stimulated effects) or PC2 FL < −0.9 (BHB-specific effects). **g-j**. Mean differential expression (log_2_FC) across the 3 *in vitro* conditions and heatmaps showing representative fuel-dependent DEGs for: **g.** Proliferation, **h.** Aerobic Glycolysis (bar graph shows the ratio of *Ldha/Ldhb* expression), **i.** Integrated stress response (ISR), and **j.** Mitochondria Oxidative Phosphorylation. All bar charts (**b., d., g.-j.**) show not significant (ns) and ***P < 0.0006 compared to the first column of data or as designated with Welch’s t-test (**b.**) or one-way ANOVA with Tukey’s multiple comparison test (**d., g.-j.**). Plots in **b.** and **d.** show mean ± s.e.m. with individual data points showing ± E/F + G or B (n = 5). Data in **e., g.-j.** show mean ± s.e.m. with n=4 for ± E/F + G and n=6 for + E/F + B.

To identify BHB dependent effects on proliferative astrocyte gene expression we conducted RiboTag IP RNA-seq and performed PCA on 15,650 DEGs detected across 3 conditions: (i) qA cells on Glc media, and proliferative astrocytes exposed to E/F mitogens in (ii) Glc or (iii) BHB media (**Fig. 7e**). PCA revealed a clear separation of samples from the three conditions, with PC1 defining differences in proliferation and related anabolic programs such as translation, ribosome biogenesis, ribonucleoprotein complex biogenesis, and DNA replication, as identified by unsupervised GO analysis (**Fig. 7e-f**). While both conditions stimulated with E/F mitogens showed upregulation of PC1-defining gene programs, astrocytes fueled by BHB showed a significant attenuation of these programs compared to cells fueled by Glc (**Fig. 7f**, **Supplementary Fig. 17**). Most notably, BHB-fueled astrocytes showed a dramatic dampening of the E/F mitogen-stimulated upregulation of proliferation genes including the same canonical genes that were prominently upregulated in astrocytes responding to MC biomaterial injections *in vivo* such as Ube2c, *Cdk1*, *Mki67*, *Top2a*, and *Kif2c* (**Fig. 7g**). BHB-fed astrocytes also had a near two-fold reduction in expression of ribosome and ribonucleoprotein complex biosynthesis genes compared to Glc-fed cells, including diminished upregulation of essential ribosome subunit and assembly molecules such as *Lyar, Rps19, Nop56, Ran,* and *Rpl13a* among others, which are vital for sustaining cell proliferation (**Supplementary Fig. 17**). Consistent with these transcriptomic differences in ribosome synthesis was a reduction in HA-positive ribosome intensity by immunocytochemistry (ICC) in BHB-treated proliferating astrocytes compared to Glc-treated cells (**Supplementary Fig. 16**). Additionally, E/F mitogen-stimulated astrocytes grown in BHB media had lower *Ldha:Ldhb* ratios compared to cells in Glc media (**Fig. 7h**), representing a significant reduction in this key prognostic marker for aerobic glycolysis metabolism^64,65^. BHB-fed cells also had lower expression of master glycolysis transcription factor, *Foxm1*, as well as critical glycolytic enzymes such as *Hk2, Gpi1, Pfkl, Pgk1 and Pgam1* (**Fig. 7g-h**). These data are consistent with reduced aerobic glycolysis when mitogen stimulated astrocytes are fueled with BHB versus Glc.

The PC2 axis defined BHB-specific effects that were consistent with an adaptive integrated stress response (ISR) from Glc deprivation and alternate fuel presentation^66^, with GO analysis on PC2-defining genes revealing upregulation of several canonical stress response programs including autophagy (*Hpse*, *Sqstm1, Tbk1, Tsc2*), pro-survival endoplasmic reticulum unfolded protein sensors and chaperones (*Ern1, Atf6, Eif2ak3, Xbp1*), and anti-apoptotic signaling (*Cflar, Faim2, Pim2*) (**Fig. 7f**). Expression of genes associated with these adaptive ISR-related programs were dramatically increased in BHB-treated cells compared to both qA and proliferating astrocytes grown on Glc (**Fig. 7i**). BHB-fueled astrocytes also showed elevated expression of mitochondrial DNA-encoded electron transport chain (ETC) sub-units (COX2, COX3, CYTB, ND1, N2, ND3, ND4, ND5, ND6) and mitochondrial ribosomal RNAs (mt-Rnr1, mt-Rnr2), all of which are needed to augment mitochondrial protein synthesis and mitochondrial capacity for oxidative metabolism (**Fig. 7j**). Notably, BHB also directed increased expression of *Ppargc1a*, a master transcriptional regulator of mitochondrial biogenesis that is an established enabler of increased oxidative phosphorylation and fatty acid oxidation capacity^67^. *Foxo1* and *Foxo3* transcription factors, which form a complex with *Ppargc1a* to sustain the protective stress responses, and production of antioxidant defense molecules such as *Sod2, Mt2, Cat, Gpx1, Nfe2l2* that are essential to supporting increased oxidative phosphorylation capacity^68^, were also all upregulated in BHB treated cells (**Supplementary Fig. 17**). BHB-fueled astrocytes also showed prominent upregulation of a diversity of transmembrane transporters, another characteristic feature of the cellular stress response, including for metal cations (*Slc9a9, Slc11a2; Slc30a4,5,7; Slc31a1; Slc38a7,10; Slc39a6,7,9,14, Slc41a2*), anions (*Slc4a10, Slc12a2,5, Slc20a1,2*), amino acids (*Slc1a2, Slc1a4, Slc6a9, Slc7a11, Slc38a2*), nucleosides (*Slc29a2*), acetyl-coA (*Slc33a1*), sugar and vitamins (Slc2a6,10; Slc5a1, 3, 6, 9; Slc19a2; Slc23a2), and hormones (Slco3a1) (**Supplementary Fig. 17**).

Collectively, these *in vitro* data show that stimulating astrocytes with BHB can suppress mitogen-stimulated proliferative responses by attenuating aerobic glycolysis metabolism and augmenting adaptive stress response programs and oxidative metabolism.

### Local delivery of BHB attenuates MC biomaterial contraction and alters astrocyte borders *in vivo*

To test whether BHB could alter astrocyte border formation around biomaterials injected into the striatum, we incorporated BHB at 20mg/ml (∼160mM) into the ABC tool formulation. In a separate group, we loaded 2-deoxy-d-glucose (2DG) into the ABC Tool also at 20mg/ml (∼122mM) for comparison, since it is a non-metabolized glucose analogue that is a competitive inhibitor of glycolysis^69^ and has previously been shown to regulate FBR around subcutaneous implants^70,71^ (**Fig. 8a**). Incorporation of BHB or 2DG did not alter the stiffness or viscoelastic properties of the MC biomaterial, and BHB readily released from MC biomaterials by diffusion, resulting in near complete recovery after 7 days under simulated sink conditions *in vitro* (**Supplementary Fig. 18**). Striatal injections of the ABC Tool releasing either of these two metabolic regulators caused no acute changes in mouse behavior on the cylinder test (**Supplementary Fig. 19**). Discrete deposits were detected at 28 dpi for both BHB- and 2DG-loaded biomaterials with HA expression both prominent and restricted to the astrocyte border zone in a manner equivalent to MC only hydrogels (**Fig. 8b, Supplementary Fig. 18**). Markedly, BHB-loaded biomaterial deposits showed negligible contraction at either 7 or 28 dpi with deposit radius and circularity parameters being equivalent to that seen for MC-only prior to onset of contraction at 4dpi as well as for the astrocyte border ablating L2AA-releasing MC biomaterials at 28 dpi (**Fig. 3e-f, 8c-e**). By contrast, 2DG-loaded biomaterial deposits contracted considerably such that they were equivalent to MC-only formulations by 28 dpi. BHB-loaded biomaterials showed no difference in Vimentin expression at the formed astrocyte border at 28 dpi but did have a significant reduction in individual astrocyte size and total Gfap expression in AB cells at both 7 dpi and 28 dpi compared to MC-only formulations (**Fig. 8b,f**, **Supplementary Fig. 20**). Despite the apparent reduction in astrocyte size and Gfap expression, the number density of astrocytes at borders around BHB-loaded biomaterials at 7 dpi was nearly double that of MC-only formulations, reducing to equivalent levels by 28 dpi (**Supplementary Fig. 20)**. Unlike for the L2AA-loaded biomaterial, BHB driven modifications to astrocyte border formation did not provoke any changes in the recruitment of Cd13-positive immune cells or fibronectin-positive fibroblasts compared to the MC-only group (**Fig. 8g, Supplementary Fig. 20**), and the extent of collagen deposition at the biomaterial-tissue interface remained negligible (**Fig. 8h**).

**Fig. 8.**
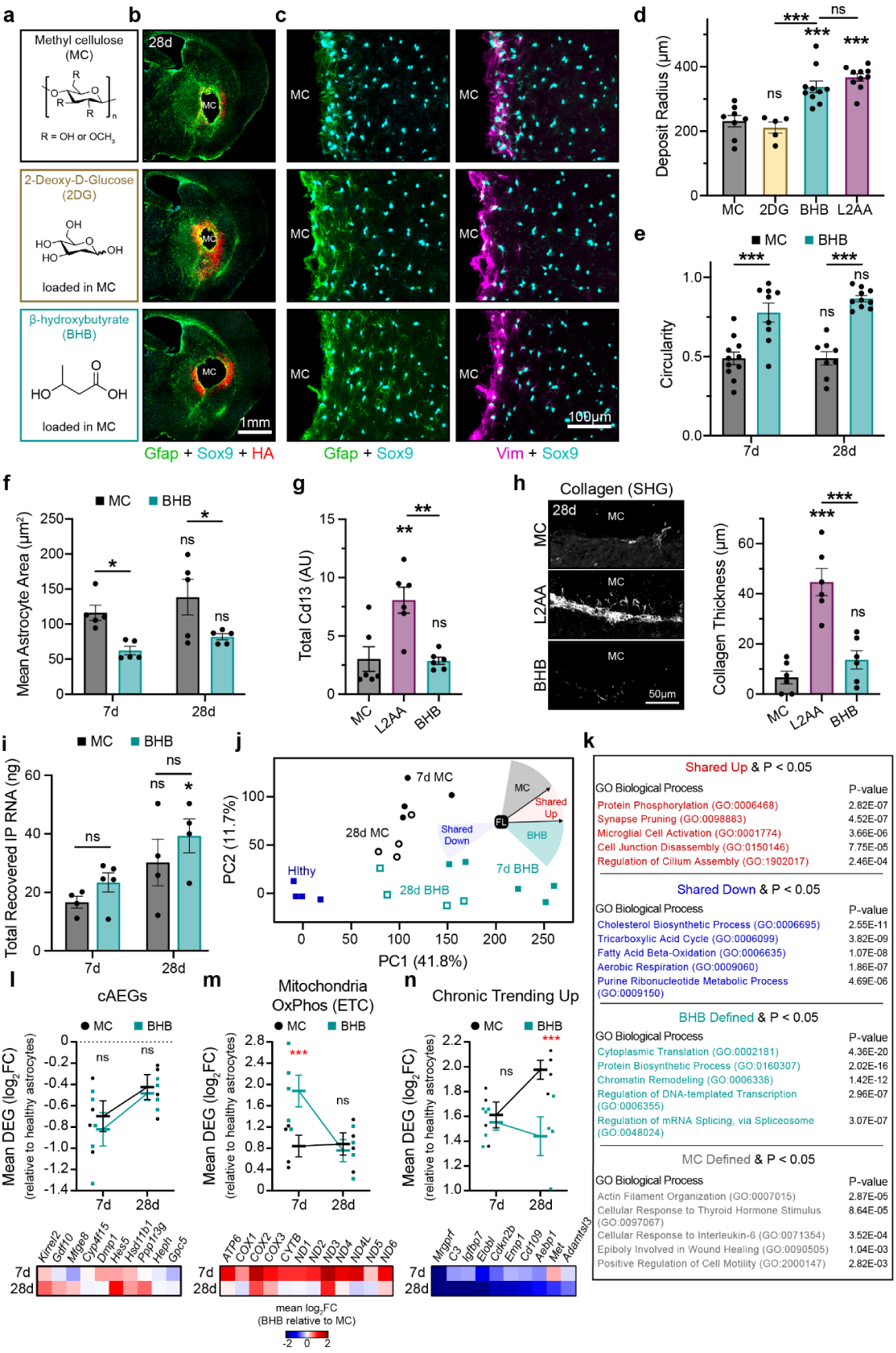
Local delivery of BHB attenuates MC biomaterial contraction and alters astrocyte border transcriptional reprogramming *in vivo*. **a.** Schematic showing chemical structures of MC, 2DG, and BHB. **b.** Overview images showing astrocyte distribution and HA expression at 28 dpi for MC only, and MC-loaded with 2DG or BHB. **c.** Detail images showing astrocyte border morphology for the three different biomaterial formulations. **d.** Quantification of biomaterial deposit radius at 28 dpi for MC only, and MC-loaded with 2DG, BHB or L2AA. **e.** Quantification of biomaterial deposit circularity at 7 and 28 dpi for MC-only and BHB-loaded biomaterials. **f.** Quantification of mean astrocyte area at astrocyte borders at 7 and 28 dpi for MC-only and BHB-loaded biomaterials. **g.** Quantification of total Cd13 within 300µm of the biomaterial-tissue interface. **h.** Overview image and quantification of fibrotic capsule thickness as measured by SHG microscopy. **i.** Total RNA recovered by RiboTag-IP for MC-only and BHB-loaded biomaterials at 7 and 28 dpi. **j.** PCA of DEGs detected as differentially regulated across MC-only and BHB-loaded biomaterials at 7 and 28 dpi relative to healthy astrocytes with the 4 categories of regulated gene lists derived from the FLVCA shown. **k.** Table showing significant GO-BPs associated with the 4 categories of regulated gene lists derived from the FLVCA. **l-n**. Mean differential expression (log_2_FC) and heatmaps across the MC-only and BHB-loaded biomaterials at 7 and 28 dpi showing DEGs for: **l.** cAEGs, **m.** Mitochondria Oxidative Phosphorylation, and **n.** Chronic Trending Up gene list. All graphs in this figure show not significant (ns), *P < 0.05, **P < 0.007, and ***P < 0.0003 either when compared to MC or as designated (**d., g.-h.**) via one-way ANOVA with Tukey’s multiple comparison test, or with comparisons made between timepoints within a group or as designated (**e.-f., i.**) via two-way ANOVA with Fisher’s LSD test, or with comparisons made between groups at a single timepoint (**l.-n.**) via two-way ANOVA with Tukey’s multiple comparison test. Graph shows mean ± s.e.m. with individual data points showing: MC (n = 8), 2DG (n = 5), L2AA (n = 10), and BHB (n = 10) (**d**); 7d MC (n = 11), 28d MC (n = 8), 7d BHB (n = 9), and 28d BHB (n = 10) (**e**); 7d/28d MC (n = 5), 7d/28d BHB (n = 5) (**f**); MC (n = 6), L2AA (n = 6), and BHB (n = 6) (**g-h**); or Healthy (n=4), 7d/28d MC (n = 4), 7d BHB (n = 5), and 28d BHB (n = 4) (**i-m**).

Given that releasing BHB altered astrocyte borders formed around MC biomaterials as assessed by IHC, we next performed translational profiling on the BHB driven changes to AB cells at acute (7dpi) and chronic (28dpi) timepoints, making comparisons with AB cells around unloaded MC-only biomaterials as well as healthy striatal astrocytes derived using the AAV solution as before. Consistent with the HA-expression detected by IHC, we recovered equivalent amounts of total mRNA after RiboTag IP from BHB-loaded biomaterials as that recovered from unloaded MC-only formulations, indicating BHB delivery did not induce astrocyte death like that observed with L2AA (**Fig. 8i**). We performed PCA on 13,516 DEGs that were detected as differentially regulated on at least one of the timepoints for either biomaterial formulation compared to healthy striatal astrocytes (**Fig. 8j**). PCA showed temporally dependent changes consistent with the prior evaluations (**Fig. 4a**), but BHB-loaded and MC-only biomaterials each showed distinct temporal reprogramming trajectories in PC space, with animals receiving MC-only biomaterial having a prominent PC2 component while this was negligible for mice receiving the BHB-loaded biomaterial (**Fig. 8j**). Performing factor analysis using the FLVCA methodology as outlined earlier, we identified genes that were up/downregulated in a nearly equivalent manner for the two biomaterial formulations, as well as genes that were more prominently regulated in each of the biomaterials such that it defined its unique reprogramming trajectory (**Fig. 8j**). GO-BPs identified from 970 genes that were equivalently downregulated in both biomaterials included essential metabolic processes previously identified, including TCA cycle, cholesterol biosynthesis, aerobic respiration, and purine ribonucleotide synthesis (**Fig. 8k**). Over 30% of cAEGs (150 out of 429) were detected as shared downregulated DEGs and there was no difference in the overall extent of the downregulation for the cAEGs between the two materials at either 7 dpi or 28 dpi (**Fig. 8l**). Notable shared downregulated cAEGs included many molecules regulating astrocyte-specific fatty acid metabolism (*Acaa2*, *Gstm1, Acsbg1, Acsl6, Slc27a1, Elovl2, Fads1*), as well as synapse maintenance and modulation (*Kirrel2, Gpc5, Sparcl1*) (**Supplementary Fig. 21**). These data suggest that BHB delivery did not alter the prominent and conserved de-differentiation and loss of essential astrocyte specific functions provoked following striatal biomaterial injection.

Shared upregulated genes for the two biomaterials determined by FLCVA (578 genes) identified GO-BPs for protein phosphorylation, synapse pruning, and microglia activation. This list of genes included 31 of 170 cARGs such as *Trem2, Tyrobp, Tgm1, C1qa, C1qb, C1qc, Fes,* and *Ptpn6* that contributed to the noted shared upregulated GO-BPs (**Fig. 8k**). Interestingly, there were 18 cARGs that were more prominently upregulated in mice receiving the MC-only biomaterial at both 7dpi and 28 dpi, including canonical reactivity genes *Serpina3n, Emp3, Gfap, Anxa3, Gpnmb,* among others, as well as genes that direct antigen processing, presentation, and phagocytosis like *Spi1, Fcg2b, Fcg3, Ctss, Ctsh*, and *Cd68*. By contrast there were only 3 cARGs (*Gpsm3, Hpgds, Csf3r*) that were more highly upregulated in AB cells around BHB-loaded biomaterials, but the expression was elevated above MC-only levels at 7dpi, but not at 28dpi. Additionally, while overall expression levels of cARGs at 7 dpi were no different between the two biomaterial groups, animals receiving the BHB-loaded formulation showed a significant reduction in mean cARGs by 28 dpi (**Supplementary Fig. 22**).

Both MC-only and BHB-loaded biomaterials evoked significantly elevated expression of canonical genes regulating proliferation and immaturity, with genes such as *Cdk1*, Igfbp2, *Ccnd1, Top2a*, *Mki67, Ube2c*, *Foxm1*, *Birc5, Kntc1* being equivalently upregulated in the two biomaterial groups at 7 dpi before returning to baseline, healthy astrocyte levels by 28 dpi (**Supplementary Fig. 23**). Thus, in contrast to our earlier *in vitro* findings, BHB delivery was insufficient to significantly attenuate pro-proliferative gene programs in border forming astrocytes *in vivo*. However, a key prognostic marker for aerobic glycolysis (the *Ldha*:*Ldhb* ratio) was significantly reduced at 7 dpi for the BHB-loaded biomaterial compared to the MC-only biomaterial, suggesting that BHB delivery may contribute to acutely attenuating the extent of metabolic reprogramming that is needed to fuel proliferative processes (**Supplementary Fig. 23).**

GO-BPs identified from the 275 genes that were determined to be more prominently upregulated in the MC-only reprogramming trajectory by FLVCA included actin filament organization, wound healing, and cell motility, which are all processes essential to reactivity and contractile functions in astrocyte borders (**Fig. 8k**). Notable molecules more elevated in mice receiving MC-only biomaterial included *Actn1, Myo1b, Gsn, Lcp1,* which serve critical actin-binding and cytoskeletal modifying functions that contribute to modifying cell shape and motility^50,72^ (**Supplementary Fig. 24**). By contrast, the BHB-loaded biomaterial reprogramming trajectory was defined by the elevated upregulation of 1132 genes by FLCVA, including molecules responsible for enabling translation and protein synthesis (*Eef1a1, Rps6kb2, Fau, Uba52*), chromatin remodeling (*Chd4, Smarca1, Smarcc1, Srcap*), and mRNA splicing (*Srrm1, Srrm4, Son, Ddx17*) (**Fig. 8k**, **Supplementary Fig. 24**). BHB is well established to act as a histone deacetylase (HDAC) inhibitor but can also modulate expression of other epigenetic machinery including histone and DNA demethylase enzymes^73,74^. AB cells around BHB-loaded biomaterials had elevated expression of such demethylase enzymes including *Kdm6b* and *Tet1* that contributed to the identified chromatin remodeling signature (**Supplementary Fig. 24**). Interestingly, key demethylase enzymes *Kdm4b* and *Kdm6b,* which remove histone methyl groups at different genomic loci, were regulated in opposing directions at 7 dpi in AB cells around BHB-loaded biomaterials. Downregulated *Kdm4b* and elevated *Kdm6b* expression like that noted in AB cells around BHB-loaded biomaterials is consistent with an epigenetic shift away from proliferative, immature, and reactive chromatin states and towards a more differentiated cell fate^75,76^. *Brd4*, another epigenetic modifying molecule essential for fatty acid metabolism^77^, was also notably increased in AB cells profiled around BHB-loaded biomaterials, suggesting a compensatory amplification of this metabolic program with BHB delivery (**Supplementary Fig. 24)**.

As before, oxidative phosphorylation and oxidative metabolic processes were significantly downregulated in AB cells around MC-only biomaterials, and local BHB delivery did not significantly prevent this altered metabolic state. In fact, the expression of hallmark oxidative phosphorylation genes on average were more downregulated in BHB-loaded biomaterials than MC-only biomaterials at 7 dpi, although there was no difference by 28 dpi (**Supplementary Fig. 25**). Closer inspection however revealed that local BHB delivery significantly lessened the acute downregulation of the master transcriptional regulator of oxidative metabolism, *Ppargc1a,* while also promoting comprehensive elevated expression of mitochondrial DNA-encoded ETC sub-units (ATP6, COX2, COX3, CYTB, ND1, N2, ND3, ND4, ND5, ND6) and mitochondrial ribosomal RNAs (mt-Rnr1, mt-Rnr2) (**Fig. 8m, Supplementary Fig. 25**). Additionally, other molecules enabling ETC processes not derived from mitochondrial DNA encoded genes returned to healthy baseline levels by 28dpi in mice receiving BHB-loaded biomaterials but remained persistently downregulated in AB cells around MC-only biomaterials (**Supplementary Fig. 25**). Thus, while BHB delivery did not substantially alleviate the apparent downregulation in the overall oxidative metabolism signature associated with the AB cell state transition, it did significantly augment the processes necessary for mitochondrial biogenesis and mitochondrial capacity to support oxidative metabolism, suggesting a partial rescue.

BHB delivery from MC biomaterials promoted a comprehensive attenuation of the expression of genes that define the chronically persisting AB cell state. In particular, mice receiving the BHB-loaded biomaterial had reduced expression of pARGs including canonical astrocyte enriched reactivity molecules like *Steap4*, *Ccn5*, *S1pr3*, *Lcn2*, *Cdsn*, A2m, *Slco1a5*, and *Fas* (**Supplementary Fig. 26**) as well as molecules assigned to the chronic gene lists from the earlier unsupervised analysis, including both the trending up and down lists (**Fig. 8n**, **Supplementary Fig. 26**). Markedly, expression of chronic trending up genes increased in MC-only biomaterials from 7 to 28 dpi as expected, but BHB-loaded biomaterials showed a significant reduction in the expression of these genes over the same time course, suggesting a critical attenuation in these delayed gain of functions (**Fig. 8n**). Notable chronic trending up genes showing this opposite temporal expression trend with BHB delivery included the complement system opsonization coordinating molecules (*C3*, *C4a*, C*4b,* and *Serping1*) as well as other critical immune modulatory factors such as *Cd109, Emp1, Aebp1, Igfbp7, Igfbpl1* (**Supplementary Fig. 26**).

Consistent with the potent immunoregulatory properties of BHB^78^, these data suggest that local BHB delivery may suppress chronic, pro-inflammatory signaling, thereby reducing the need for maintenance of defense capabilities in chronic AB cells for sustained protection of biomaterial adjacent neural tissue.

Collectively, these data show that local delivery of BHB from injected biomaterials can alter some, but not all, facets of the temporally regulated reprogramming associated with astrocyte border formation, leading to a modest reduction in the immune regulatory gain of functions in chronic AB cells and attenuated biomaterial contraction. Importantly, these findings provide proof-of-principle that local delivery of metabolic regulators from implanted biomaterials are viable candidate molecules for manipulating CNS FBRs.

## Discussion

In this work, we show that minimally invasive injection of a biomaterial into the CNS stimulates temporally regulated adaptive reprogramming and proliferative reactive changes in local astrocytes, producing transient and persistent functional changes reminiscent of astrocyte wound repair responses at CNS injuries^29,43^. Through local ablation studies, we demonstrated that astrocyte border formation is an indispensable facet of CNS FBRs, such that inhibiting the response exacerbates inflammation and fibrosis. Furthermore, by incorporating local delivery of molecular regulators targeting specific mechanisms of astrocyte reprogramming, we established proof-of-principle that astrocyte border phenotypes around biomaterials can be therapeutically modulated. These insights were enabled by the development and validation of the ABC Tool, a new platform technology that has the potential to transform our understanding of the important biomaterial properties that dictate FBRs in the CNS and other organ systems. These findings have broad implications for molecular profiling of FBRs at implanted biomaterials, advance our mechanistic understanding of astrocyte border functions, and provide a path forward for leveraging molecular regulators of astrocyte metabolism to manipulate AB cell states at implants.

Firstly, this work provided a new method for deriving detailed transcriptional information on the spatially defined AB cell population at injected biomaterials *in vivo*. Single cell (scRNA-seq)^79^ and spatial^80^ transcriptomics have become prevailing techniques in the field, since they provide insights into the phenotypic heterogeneity of cell populations and the nature of intercellular signaling networks. These techniques have been employed to profile the cellular complexity of FBRs to various biomaterials including at scaffolds implanted subcutaneously^81^ or intramuscularly^82^, at miniaturized breast implants^83^, and at implanted neural electrodes^84^. However, these methods have notable limitations. Currently, scRNA-seq achieves modest read depth per cell so this technique can not readily detect lowly expressed genes or subtly regulated genes. Sequencing artifacts with scRNA-seq can also frequently lead to overclassification of cell subtypes, while the need for live cell dissociation, isolation, and sorting can induce reactive changes in cells prior to transcriptomic evaluation that can distort the natural *in situ* profiles^79,85,86^. Spatial techniques also suffer from low read depth, loss of mRNA quality from poor tissue processing, and a lack of cell type specificity/single cell resolution^80^. Thus, while these methods may provide some valuable insights into how distinct populations of astrocytes transcriptionally reprogram into AB cells and the heterogeneity amongst individual AB cells^87^, we saw the need to develop an alternative, more robust approach to rigorously characterize the full extent of reprogramming involved in establishing chronic AB cell states at injected biomaterials.

We turned to the RiboTag affinity purification method to achieve high quality, detailed information about the AB cell translatome. RiboTag, much like other affinity purification methods such as bacTRAP and INTACT, enables enrichment of RNA from specific cell populations via targeted immunoprecipitation to permit translational profiling via IP-enriched bulk RNA sequencing, providing enhanced read depth and overall coverage with low cost per sample compared to the other aforementioned methods^88–90^. The ABC Tool developed here employed an established and well validated AAV for RiboTag expression under an astrocyte-specific promoter. The critical feature of the ABC Tool that was indispensable to effective AB cell state profiling was the spatial confinement of the RiboTag labelling achieved by loading and releasing the AAV directly from the injected biomaterial. Thus, by using the ABC Tool such that only the astrocytes within a tissue region 250 µm from the biomaterial-tissue interface were profiled, we effectively turned a cell-specific, but bulk RNA evaluation method into a biomaterial-tissue interface-defined, spatial transcriptomic analysis with maximum possible read depth and coverage. Our data show that the enrichment of AB cell RNA permitted by this approach led to at least a 14-fold increase in detected DEGs compared to pre-labelling astrocytes within discrete neuropil volumes prior to biomaterial injection. Notably, the ABC Tool facilitated detection of nuanced, but potentially therapeutically modulatable changes in astrocytes functions, including in oxidative metabolism as explored here, that would have otherwise gone undetected using the other less sensitive approaches.

It is uncertain whether the spatially confined AAV delivery noted here is unique to the particular MC biomaterial formulation employed or to the striatal tissue site used for biomaterial injections, an important consideration when incorporating the tool into other biomaterial systems or applying it to other brain regions to profile their unique AB cell states. In our prior work, we demonstrated that specific surface chemistries presented on injected biomaterials induce recruitment and persistence of different, but well defined, non-neural niches^24^. For example, nonionic sulfoxide-based biomaterials evoked minimal immune cells, cationic amine-based biomaterials promoted acute inflammation that resolved into fibrotic scar, and sulfonium-functionalized biomaterials induced a persistent neutrophil response phenotypically like chronic granulomas^24^. These different non-neural niches appear to alter the morphology of the adjacent astrocyte borders by IHC staining, but whether these different biomaterial surface chemistries elicit different transcriptional reprogramming in astrocyte has yet to be determined, representing a logical next step for use of the ABC Tool. However, caution must be exercised when considering broader use of the ABC Tool. For example, we showed here that RiboTag expression was not sufficient until 7dpi, implying that if there are critical functional changes that are isolated to within the first week post injection, they may be missed. Additionally, local molecule ablation significantly impaired astrocyte border labeling and RNA recovery, which may mean that biomaterials that stimulate similar levels of peripherally derived inflammation or local neural damage may be ill-suited for investigation with the ABC Tool. Other biomaterial physiochemical properties that do not elicit excessive inflammation but that may influence astrocyte border states, including biomaterial mechanical properties (e.g. stiffness and viscoelasticity)^91–94^ or biomaterial microstructure and porosity^95^, may be more appropriate for future investigations with the ABC Tool.

Since the ABC Tool employs AAVs to achieve the necessary transgenic manipulations, it is inherently a modular platform, limited only by what can be packaged into AAV constructs. Therefore, the ABC Tool could be readily adapted to incorporate other recently developed molecular tools that have been packaged into astrocyte-specific AAVs such as CRISPR/Cas systems to genetically edit critical reprogramming molecules^96,97^, or proximity-dependent biotinylation (BioID2) to assess protein expression of the most critical pARGs^98,99^. Additionally, use of alternative AAV serotypes and promoters could enable variants of the ABC Tool to be employed to transcriptionally profile other important features of FBRs, including specific immune and fibrotic responses, both in the CNS and in other organs/tissue contexts. Exploring such complementary molecular evaluations will serve to further refine our understanding of the role astrocyte border formation plays in the context of CNS FBRs and enable new mechanism-based approaches for improving implant biocompatibility.

Our findings here improve our understanding of the conserved adaptive reprogramming mechanisms involved in astrocyte border formation by complementing insights previously derived from transcriptional profiling of astrocyte wound responses at CNS injuries^29,43^. Our data shows that biomaterial injections elicit strikingly similar astrocyte reprogramming as that observed at CNS injuries, including a strong congruence in transient, persistent, and delayed regulated astrocyte-enriched genes. For example, the acute astrocyte dedifferentiation and de-prioritization of critical homeostatic astrocyte-neuron interactions detected at MC biomaterials mirrored the same loss of functions previously seen in astrocytes at spinal cord injuries^29^, including persistent downregulation of astrocyte enriched molecules directing synapse maintenance and modulation. Other notable similar functional changes detected in AB cells at biomaterials and CNS injuries included the transient upregulation of proliferative and immaturity gene signatures as well as the persistent but delayed increase in molecules regulating host defense and immune regulatory mechanisms. The conserved nature of these transient and persistent gain-of-functions suggests that they are relevant targets for incorporation into future therapeutic strategies to minimize or strengthen astrocyte border responses depending on the context. A previously identified facet of the acute astrocyte dedifferentiation response at CNS injuries^29^, but whose significance was perhaps underappreciated by us until reemerging in this current work, was the dramatic metabolic reprogramming that occurs to fuel AB cell state transitions, including the persistent downregulation of fatty acid and oxidative metabolism and transient upregulation of aerobic glycolysis pathways. Previous work has demonstrated that astrocytic oxidative phosphorylation and fatty acid metabolic processes are linked, and that targeted loss-of-function of oxidative phosphorylation causes lipid accumulation and reactive changes in astrocytes that exacerbates neuroinflammation and neurodegeneration^100^. This raises the question of whether persistent suppression of these critical and astrocyte-specific metabolic programs is a major contributor to the chronic upregulation of reactivity gene programs in astrocyte borders, and whether correcting this metabolic dysfunction may be a way to attenuate reactivity.

As part of a first attempt to modulate the metabolic reprogramming associated with astrocyte border formation, we demonstrated that delivering BHB locally from biomaterials altered several reprogramming processes and attenuated chronic reactivity. Specifically, local BHB delivery prevented astrocyte-directed contraction of the biomaterial and enhanced the expression of mtDNA-encoded transcripts for molecules comprising the electron transport chain and mitochondrial biogenesis, while diminishing acute aerobic glycolysis. Together these results are consistent with a direct, BHB-dependent modulation of the natural metabolic reprogramming stimulated by biomaterial injection. Since mtDNA depletion induces reactive changes in astrocytes^101^, our findings point to a possible mechanism by which BHB directly contributes to attenuating reactivity gene expression in AB cells by enhancing necessary mitochondrial activity that could warrant further investigation. A relevant limitation of this study is that we could not establish the BHB release kinetics from the biomaterial *in vivo,* so we do not know for how long therapeutic concentrations of BHB are being delivered from the biomaterial following injection. Since *in vitro* release data suggests that BHB is likely to be depleted from the biomaterial over the first 7 days, developing engineered strategies to extend BHB release locally for several weeks or testing if incorporating frequent, systemically administered doses of BHB further reduces the reactivity signature in chronic astrocyte borders will be important future work. BHB is just one of many potential metabolic regulators that may be effective at modulating astrocyte responses at biomaterials. While 2DG is a common metabolic regulator that has been employed to alter FBRs previously^70,71^, it showed minimal effect on the astrocyte border phenotype or the induced biomaterial contraction by IHC in this study. A potential reason for why delivery of BHB had a more potent effect on astrocytes compared to 2DG could lie in the uniquely high expression of the relevant carboxylate transporter, Mct1, in both healthy astrocytes and AB cells^102–104^. Exploring other molecules with equivalently high affinity for Mct1 such as oxamic acid, an isostere of pyruvate that can attenuate glycolysis via *Ldha* inhibition ^105–107^, could bolster the effects of BHB and further attenuate reactivity in chronic astrocyte borders. The transcriptomic dataset generated here could also be used to identify additional astrocyte-specific transporters uniquely upregulated in AB cells, enabling preferential delivery of other potentially more potent metabolism-regulating molecules. Should future studies confirm that administering BHB or other metabolic regulators attenuates astrocyte border formation, incorporating such treatments into device design or postsurgical management protocols for chronically implanted neural devices would represent a practical strategy to improve long-term device performance.

## Materials and Methods

### Materials

All solvents used in this work are certified ACS grade or higher. From ThermoFisher, we purchased: B27 (no Vitamin A) (50X) (Cat # 12587010), Antibiotic-Antimycotic (100X) (Cat# 15240096), MEM Non-Essential Amino Acids Solution (100X, Cat # 11140050), DMEM (Cat# A1443001), Advanced DMEM/F12 (Cat #12-634-028), Pierce A/G Magnetic Beads (Cat # PI88803), Ultrapure DNase/RNase Free Distilled Water (Cat # 10-977-023), G-Biosciences Nonidet P-40 substitute (Cat# 50-103-7034), From Peprotech, we purchased: Epidermal Growth Factor (EGF) (Cat# AF-100-15-100UG), Fibroblast Growth Factor (FGF) (Cat# 100-18B-100UG). From Gibco, we purchased: Advanced Dulbecco’s Modified Eagle Medium/ Nutrient Mixture F-12 (DMEM/F12) (Ref# 12634-010), Phosphate Buffered Saline (PBS) pH 7.4 (1x) (Ref# 10010-023). From Addgene, we purchased pZac2.1-GfaABC1D-Rpl22HA plasmid (#111811) which then underwent Midiprep and AAV5 large scale packaging by Charles River to produce 500uL of AAV5 GfaABC1D-Rpl22HA at a titre of 1×10^13^ GC/ml. From Sigma Aldrich, we purchased: methylcellulose (Methocel® A15C, Cat# 64625-100G-F), L-2-Aminoadipic Acid (Cat. # A7275-250MG), β-hydroxybutyrate (Cat. # 54965-10G-F), 2-Deoxy-D-glucose (Cat # 25972), L-N⁵-(1-Iminoethyl)ornithine, Dihydrochloride (400600-20MG), 100X Nucleosides (EmbryoMax, Cat #ES-008-D), Potassium Chloride solution (1M) (60142-100ML-F), Magnesium Chloride solution (1M) (Cat# 63069-100ML), Cyclohexamide (Cat# C7698-1G), Protease Inhibitor Cocktail (Cat# P8340-5ML), Tris 1M, pH 7.4, (Cat# T2194-1L), KIMBLE Dounce tissue grinder set (2mL) (D8938-1SET), DL-Dithiothreitol solution (646563-10X 0.5mL), Heparin sodium salt from porcine intestinal mucosa (H3393-100KU).

### Dynamic Mechanical Rheology

Rheological testing of MC biomaterials and MC biomaterials loaded with molecules was performed using a TA Instruments DHR-2 Rheometer, at 25°C with a 20mm, 2° cone and plate geometry. For each experimental run, approximately 200µL of polymer solution was prepared. The plateau storage modulus was determined by oscillating at 10 rad/sec, 0.5% strain for 3 minutes to pre-stress material. The material was then allowed to rest for 2 minutes before performing a strain sweep from 0.5% strain to 500% strain, oscillating at 10 rad/sec. Plateau storage modulus was taken as the first value of this strain sweep. Temperature dependence of the rheological properties was determined by testing rheological properties at 40°C while oscillating at 0.5% strain and 10 rad/sec. Stress relaxation properties of MC biomaterials and PBS solution control were measured on the DHR-2 Rheometer by applying a fixed strain of 15% and measuring the change in stress over 500 seconds.

### Animals

All studies were performed using young adult female C57BL/6 mice (Cat#000664, JAX) that were 8-12 weeks old at the time of experimental procedure. Animals were housed in the Boston University vivarium with a 12 h light–dark cycle, controlled temperature (20–25 °C), and were allowed free access to food and water. All *in vivo* experiments involving the use of mice were conducted according to protocols approved by Boston University’s Institutional Animal Care & Use Committee (IACUC) under IACUC Protocol numbers: PROTO202000045-TR01 and PROTO202100013-TRO1.

### Surgical Procedures

All surgical procedures were approved by the BU IACUC and conducted within a designated surgical facility. Prior to surgery, general anesthesia was achieved through inhalation of isoflurane in oxygen-enriched air. Mice heads were shaved before being stabilized and horizontally leveled in a stereotaxic apparatus using ear bars (David Kopf, Tujunga, CA). Using an operating microscope for visual aid, a small craniotomy over the left coronal suture was performed with a high-speed surgical drill. A small rectangular section of bone was removed to expose the brain in preparation for material or solution injection.

To prepare hydrogels for injection, methylcellulose was first sterile filtered and lyophilized. 85µL of the hydrogel formulation (5 wt% MC in 1x PBS) was prepared by adding 85µL of PBS to 5mg of MC. For MC hydrogels loaded with a secondary molecule, solutions of this molecule were prepared at a higher concentration and added to bring the total concentration to the following: L-N^5^-(1-Iminioethyl)ornithine (LNIO, 30 mg/mL), L-2-Aminoadipic Acid (L2AA, 20 mg/mL), 2-Deoxy-D-Glucose (2DG, 20 mg/mL), and β-hydroxybutyrate (BHB, 20 mg/mL). The hydrogel was then allowed to hydrate in a fridge for 72 hours before surgery. On the day of surgery, 10µL of 0.97E13 gc/mL HA RiboTag AAV solution was mixed vigorously into the hydrogel using a pipette tip. The hydrogel material was then backloaded into a pulled glass pipette for injection. 1µL of hydrogel material was injected into the caudate putamen nucleus at 0.2 μL/min using target coordinates relative to Bregma: +1.0 mm A/P, +2.5 mm L/M and −3.0 mm D/V. 3 min of dwell time was allowed post injection before slowly removing the pulled glass pipette.

Infusions of HA AAV RiboTag solution were performed by injecting 1 µL of AAV solution at 0.97E12 gc/mL at 0.15 µL/min and allowing for 5 min of dwell time post injection. To induce focal ischemic strokes, 1.5 μL of LNIO (27.4 mg/ml or 130 µM in sterile PBS) was injected at 0.15 μL/min.

All injections were made using a standard micropipette injection protocol. This protocol uses a pulled borosilicate glass micropipette (WPI, Sarasota, FL, #1B100-4) that has been ground to a 35° beveled tip with 150–250 μm inner diameter. Glass micropipettes were mounted to a stereotaxic frame via specialized connectors and attached, via high-pressure polyetheretherketone (PEEK) tubing, to a 10 μL syringe (Hamilton, Reno, NV, #801 RN) controlled by an automated syringe pump (Pump 11 Elite, Harvard Apparatus, Holliston, MA). After injection, the surgical incision site was sutured closed, and the animals were allowed to recover.

### Transcardial perfusions for Immunohistochemistry

Mice were first given terminal anesthesia by overdose of isoflurane, before being perfused transcardially with heparinized saline (10 units/ml of heparin) and 4% paraformaldehyde (PFA) that was prepared from 32% PFA Aqueous Solution (Cat# 15714, Electron Microscopy Science), using a peristaltic pump (flow rate of 7 mL/min). Approximately, 10 mL of heparinized saline and 50 mL of 4% PFA were used per animal. Following perfusion, brains were immediately dissected and post-fixed in 4% PFA for 6-8 hours. After PFA post-fixing, brains were cryoprotected in 30% sucrose + 0.01% sodium azide in Tris-Buffered Saline (TBS) for at least 3 days with the sucrose solution replaced once after 2 days and stored at 4°C until further use.

### Immunohistochemistry (IHC)

Coronal brain sections (40 µm thick) were cut using a cryostat (Leica CM 1950). After cutting, tissue sections were stored in TBS buffer + 0.01% sodium azide at 4°C until retrieval for IHC. Tissue sections were processed for immunofluorescence staining using previously described free floating staining protocols^24^. Briefly, antigen retrieval was done using 1N hydrochloric acid. Then, normal donkey serum and triton X-100 were used to block and permeabilize tissue. The primary and secondary antibodies used for IHC are listed below. Secondary antibodies were diluted 1:250 prior to incubation. Cell nuclei were stained with 4’,6’-diamidino-2-phenylindole dihydrochloride (DAPI; 2 ng/ml; Molecular Probes). Tissue sections were mounted on glass slides and coverslips were secured using ProLong Gold anti-fade reagent (Cat# P36934, Invitrogen). Stained sections were imaged using epifluorescence and deconvolution epifluorescence microscopy on an Olympus IX83.

### Antibodies for Staining

Primary antibodies used in this work include: rat anti-Gfap (1:1,000, 13-0300, ThermoFisher), guinea pig anti-Gfap (1:500, 173 308; Synaptic Systems), rabbit anti-hemagglutinin (1:1,000, H6908; Sigma-Aldrich), goat anti-hemagglutinin (1:800, NB600-362; Novus Biologicals), rat anti-Lgals3 (1:200, 14-5301-82; ThermoFisher), goat anti-Sox9 (1:500, AF3075; R&D Systems), guinea pig anti-Vimentin (1:500, 172 004; Synaptic Systems), goat anti-Kirrel2 (1:200, AF2930; R&D Systems), sheep anti-BrdU (1:500, NB500-235; Novus Biologicals), rabbit anti-MCT1 (1:500, 356 003; Synaptic Systems), guinea pig anti-Iba1 (1:500, 234 004; Synaptic Systems), rabbit anti-Fibronectin (1:500, AB2033; Millipore), goat anti-Cd13 (1:500, AF2335; R&D systems), rabbit anti-Top2a (1:500, AB52934; ABCAM), rabbit anti-P2y12 (1:800, AS-55043A; AnaSpec), goat anti-Timp1 (1:100, AF980-SP; R&D Systems), guinea pig anti-S100β (1:1000, 287 004; Synaptic Systems), mouse anti-µ-crystallin (Crym) (1:200, sc-376687, Santa Cruz Biotechnology).

Secondary antibodies were purchased from Jackson ImmunoResearch Laboratories with donkey host and target specified by the primary antibody (e.g. AffiniPure™ Donkey Anti-Rabbit IgG secondary antibodies conjugated to Alexa Fluor 488 (AB_2313584), Alexa Fluor 555 (AB_3095471), Alexa Fluor 647 (AB_2492288), or Alexa Fluor 790 (AB_2340628)).

### IHC Image analysis

All staining intensity quantification was performed on tissue stained concurrently and imaged with standardized exposure time and evaluated using a standard intensity setting. Quantification of staining intensity was performed using the Radial Profile Angle plugin on NIH Image J software (Version 1.53k). Radial angle profiles (RAPs) of 1000 or 1500 µm (1538 or 2308 pixels) were used across this entire study. Quantification was performed from the center of the injection, with the analysis determining the average pixel intensity across the circumference of a circle at each radial pixel distance. Total values for staining intensity (ie. total HA) was determined by taking the integral (area under the curve) of the radial intensity profile. The deposit size was determined by normalizing the initial RAP value for DAPI to 0 and the average of the final 100 values to 100. Border location was determined by the location where the DAPI intensity crossed a threshold of +50. Measurement of percent of total HA-positive astrocytes located within the Vimentin-positive astrocyte border (**Fig. 1h**) was calculated by multiplying the % of total HA expression at 250 µm and the % of Vim+ / HA+ astrocytes in the border within the first 250 µm. Measurement of %HA expression in Vim+ region (**Supplementary Fig. 2b**) was determined by finding the end of the vimentin border using the first derivative (defined as the first derivative crossing a threshold of 20% its maximum value) and finding the fraction of the total HA intensity within that region. Analyses of shape parameters for MC deposits were performed by tracing deposit profiles and quantifying using Image J.

### Astrocyte Border Morphology and HA+ Vimentin+ Quantification

Astrocyte border morphology and HA+ / Vimentin+ quantification was assessed using a combination of modified ImageJ macros and custom MATLAB code. Briefly, channels were adjusted for brightness and contrast, thresholded, and denoised in ImageJ. Niblack and Sauvola local thresholds were used for Gfap and Vimentin, respectively, while the moments threshold was used for the Sox9 and HA channels.

Scripts adapted from the MicrogliaMorphology package^108^ were used to generate regions of interest (ROIs) of cell outlines defined by the Gfap channel and to detect the presence of either Sox9, HA, or Vimentin within the boundaries of the cell. Data output from ImageJ was processed using MATLAB. ROIs with area less than 300 px^2^ were excluded from the analysis. The number of nuclei in each ROI was given by the number of Sox9 particles within the outline. ROIs with a single Sox9 particle were classified as “single-domain” and those containing multiple Sox9 particles were classified as “overlapping-domain.” ROIs were classified with MATLAB, and the class label outputs were fed back into MicrogliaMorphology to recolor astrocytes. HA and Vimentin positivity was determined by the presence of HA or Vimentin within the bounds of the ROI. Average astrocyte area was calculated by dividing the total area of all astrocyte ROIs by the number of detected nuclei. The numerical density of astrocytes was determined as the number of detected nuclei within the specific border area.

### Second Harmonic Generation (SHG) Imaging using Two-Photon (2P) Microscopy

2P SHG microscopy was used to assess collagen content following astrocyte ablation using MC loaded with LNIO or L2AA. Briefly, tissue samples were mounted on a coverslip and kept hydrated using 1xTBS. Tissue was imaged on a Bruker Ultima Investigator two-photon microscope with a tunable Ti:Sapphire laser (Insight X3, Spectra Physics) and a 4x objective (ZeissPlan-Neofluar, NA = 0.16) with an image size of 1024 x 1024 pixels. Second harmonic generation was achieved by exciting tissue at a wavelength of 950 nm and acquiring Z stacks in 1-micron intervals. Images were processed as average intensity Z projections and analyzed using RAPs on ImageJ as previously described. Collagen thickness was calculated as the average of the width taken at the medial and lateral border. High resolution images were taken using a 20x objective (Olympus XLUMPlanFLNXW, NA = 1.0).

### Cylinder Rearing Test

Mouse motor function and basic behavior was assessed using the cylinder rearing test. This test characterizes spontaneous forelimb usage when exploring a clear, plastic cylinder oriented vertically ^109^. For this test, mice were placed in the cylinder (Clear Cast Acrylic Tube 4-1/4" OD x 4" ID - 1 ft long, McMaster-Carr 8486K943) at baseline (1-3 days before surgical intervention) and were recorded for 5 minutes in a dimly lit room. A mirror was placed behind the cylinder to allow for observation of rears in all directions. This was repeated at specified intervals following surgical intervention (1d, 3d, 7d, 14d, 28d). Subsequently, analysis of the video was performed by a single observer, who recorded the number of rears by the animal and the primary or preferential forepaw used, as determined by the highest forepaw placement. This was then collated to determine the percentage of right forepaw reaches and percent change in total activity relative to baseline to characterize both functional and behavioral changes.

### Neural Progenitor Cell (NPC) Culture and Differentiation

NPCs were cultured as previously described^42,63^. Briefly, the cells were grown in neural expansion (NE) media consisting of Advanced DMEM/F12 supplemented with B27 (no Vitamin A) (50X), Antibiotic-Antimycotic (100X), Nucleosides (100X), MEM Non-Essential Amino Acids Solution (100X) and 100 ng/ml each of Epidermal Growth Factor (EGF) and Fibroblast Growth Factor (FGF). NPC stocks were maintained so that all cell stocks used in this study were less than P30.

For all *in vitro* studies, 30,000 to 40,000 cells were seeded onto 10 mm round coverslips that were coated with 0.1% gelatin and placed into individual wells of a 24 well plate. Cells were cultured on gelatin coated coverslips for 2-6 days in 1 ml of media. At designated endpoints, the media was removed, the coverslips were rinsed with 1X PBS and then fixed with freshly prepared 4% paraformaldehyde for 30 minutes.

For astrocyte differentiation, NPCs were seeded onto coverslips with astrocyte differentiation media for 48 hrs. Astrocyte differentiation media either consisted of NE media and 2% fetal bovine serum (FBS) or DMEM (no glucose, no glutamine, no phenol red) that was supplemented with all the ingredients for NE media mentioned previously, 2% FBS, and 20 mM β-hydroxybutyrate (BHB). To test proliferative capacity under the two different fuel sources, astrocytes were exposed to 50 ng/mL E/F, while removing the 2% FBS for 4 days. To enable evaluation of proliferative capacity, bromodeoxyuridine (BrdU) (Cat# B15751G, TCI America) at 20 μM was added 6 hrs before fixation.

### Immunocytochemistry (ICC)

For ICC staining, fixed coverslips were stored in 24-well plates at 4°C under 1x PBS containing 0.01 wt% sodium azide until retrieval for staining. The staining procedure followed the same general protocol described above for IHC, without antigen retrieval, and has been described previously^63^. The primary and secondary antibodies used for ICC are included in the list above.

### Transcardial Perfusion for Transcriptomics

Mice were first given terminal anesthesia by overdose of isoflurane, before being perfused transcardially with heparinized saline (10 units/ml of heparin) prepared using DNase-free/RNase-free water and 10× PBS, using a peristaltic pump at a rate of 7 mL/min. Approximately, 14 mL of heparinized saline was used per animal. Following perfusion, brains were immediately dissected on ice, placed in a DNase-free/RNase-free 5 mL conical tube and flash frozen using liquid nitrogen. Tissue could be stored at - 80°C for 6-12 months before processing for transcriptomics.

### Tissue Processing and RiboTag immunoprecipitation

Frozen brain tissue was processed for RiboTag immunoprecipitation^90^ using established methods^8,29,42,110^. A brief overview of the steps follows: a single sample was retrieved from cold storage and allowed to warm on ice for 3-5 min. The brain was then placed in a sectioning block (Electron Microscopy Sciences, Cat. # 69090-C) for 15-30 seconds before blocking. Blocking was achieved by placing a clean razor blade at lambda and a second razor blade at the caudal side of the olfactory bulbs to hold the brain in place. Then, two chilled blades were used to isolate a 3 mm section of tissue containing the site of injection. Finally, the ipsilateral striatum, which contained the injection site, was biopsied using a 3 mm biopsy punch and deposited in a dounce containing RiboTag homogenization/lysis buffer^42,110^. Tissue was homogenized using two glass pestles and centrifuged to remove tissue debris. Isolation of hemagglutinin-positive ribosomes was performed by incubating the solution with an anti-HA.11 Epitope Tag Antibody (Biolegend, Cat #901515) for 4 hours in a microcentrifuge tube on a microtube rotator at 4 °C. Immunoprecipitation solutions were combined with Pierce A/G Magnetic Beads (Thermo Scientific, Ref. # 88803) and incubated overnight on a microtube rotator at 4 °C.

The following day, the immunoprecipitation solution was processed by placing the microcentrifuge tubes in a magnetic rack (DynaMag-2, Invitrogen, Ref. # 12321D) to separate the solution from the magnetic beads. The separated solution was processed as the “Flow through” samples, which represents the mRNA from all RiboTag-negative cells. These Flow through samples were then mixed with RLT Plus buffer with 2-mercaptoethanol (BME), purified using RNeasy Plus Mini kits (Cat. # 74134), and stored at −80°C. To recover the immunoprecipitated (IP) RNA from the RiboTag-positive cells, magnetic beads were washed three times with a high salt solution^42,110^. This unpurified RNA was separated from the magnetic beads using the magnetic rack, mixed with RLT Plus buffer with BME, and purified using RNeasy Plus Micro kits (Cat. # 74034). mRNA quantity and quality were assessed using an Agilent TapeStation 4150 and samples having both sufficient RNA quantity and an RNA integrity number equivalent (RINe) greater than 6.8 were submitted for sequencing. Sequencing was performed by GeneWiz (Azenta) using sequencing was performed on poly-A selected libraries using Illumina NovaSeq 6000 (Genewiz) using pair end reads (2×150 – 150bp length) with a minimum of average of 50M reads per sample.

### Analysis of RiboTag RNA-seq transcriptomics data

Raw RNA-seq data was processed in Galaxy using previously established workflows^29,42^. Briefly, the R1 and R2 FASTQ files obtained from the sequencing run were cleaned up using the Trimmomatic tool.

Data was then aligned to the most recent M. musculus reference genome (mm39) and a hemagglutinin (HA) reference using the HISAT2 tool. Counts of expressed genes from the aligned datasets were quantified using the featureCounts tool. Subsequently, fragments per kilobase of transcript per million mapped reads (FPKM) values were calculated using Microsoft Excel. FPKM values normalize for individual gene length and total number of counts per sample. Differentially expressed genes (DEGs) and fold-enrichment (FE) of transcripts were quantified by using the EdgeR tool in Galaxy on raw gene count data. Across all studies, a false discovery rate (FDR) cutoff of <0.1 was used to define significantly regulated DEGs. For assessing DEGs of individual samples and comparing to curated lists, DEGs were manually calculated as the log_2_FC of FPKM values + a nominal offset of 0.0005. Chronically regulated genes were defined as DEGs regulated after 7 dpi but persisting through 70 dpi. To categorize chronically upregulated genes as trending up, trending down, or not changing, we examined the change in differential expression between acute and chronic timepoints (ΔDEG), and defined genes that are trending up as having a ΔDEG > 0.2, genes trending down as having a ΔDEG < −0.2, and genes that are not changing as having a ΔDEG between −0.2 and 0.2. Additional tools used to assess or represent gene expression include: Enrichr, which was used for all gene ontology (GO) analyses (https://maayanlab.cloud/Enrichr), and NG-CHM BUILDER (https://build.ngchm.net/NGCHM-web-builder), which was used to generate heatmaps of DEG data.

### Principal Component Analysis (PCA)

PCAs were performed using XLStat (Addinsoft, Version 2025.1.3)^111^. All PCAs were performed on either the individual FPKM values or the average FPKM values (on a per condition basis) of identified DEGs. PCA plots were generated using the first two principal components and the samples were plotted using their relative factor scores in PC1 and PC2. PCA graphs were scaled to accurately represent the Euclidean distance. Analysis of PCAs to determine genes correlated with a particular principal component used a factor loading threshold of |PC Factor Loading| > 0.9. GO analysis of these genes was performed using Enrichr as described above.

### Factor Loading Vector Cartesian Angle (FLVCA) analysis

The Factor Loading Vector Cartesian Angle (FLVCA) for each expressed gene was calculated by determining the PC1 and PC2 factor loadings (FL) from the PCA for a given gene and using the following equation to solve for θ (where θ is the FLCVA) and factoring in the cartesian quadrant where the FL vector is located in PCA space: tan(θ) = PC2-FL / PC1-FL. Genes with a FL vector magnitude >0.8 were used for analysis. The angle ranges used to define the various gene categorizations for the FLVCA analysis were determined empirically and dependent on where the AB cell conditions were located in PCA space as well as the vectors formed between individual AB cell conditions relative to either flow through samples (**Fig. 2h-j** analysis) or healthy astrocytes (**Fig. 8j-k** analysis).

### Threshold criteria for gene expression in healthy astrocytes

Healthy astrocyte expressed genes (AEGs) were defined as having an FPKM value greater than 0.02, which was a conservative cutoff that accounted for genes that were no more than one standard deviation below the mean of the log10-transformed dataset and represented the upper 81% of all genes that had detectable counts in the dataset. Genes with an FPKM value at or above this threshold in healthy astrocytes were considered healthy expressed genes.

### BHB Release Assay

To measure BHB release from MC hydrogels, MC was first solubilized at 5 mg/mL in DI water and 1 mL of MC solution (5 mg MC) was added to microcentrifuge tubes, before being frozen and lyophilized to create identical 5 mg MC stocks. BHB was then solubilized at 20 mg/mL in 1x PBS and 95 µL of 20 mg/mL BHB solution was added to the 5 mg MC aliquots to make BHB-loaded 5 wt% MC gels. As a control, unloaded MC gels (ie. PBS only) were fabricated following the same process. Materials were allowed to solubilize for 72 hrs at 4°C before beginning the release assay. For the release assay, BHB-loaded and MC-only gels were added to Slide-A-Lyzer® MINI Dialysis Units (3,500 MWCO, ThermoFisher, Cat # 69550) and then were incubated in 1 mL PBS for 7 days at 37°C, with sampling on alternate days and full buffer replacement. All collected samples were assessed using a beta-Hydroxybutyrate Assay Kit (Colormetric, Novus Biologicals, Cat# NBP3-25919-96T) and read on a TECAN (Infinite M Plex) Microplate Reader using an absorbance of 450 nm as specified by the BHB Assay Kit.

### Statistical Analysis

Graph generation and statistical evaluations of repeated measures were conducted by one-way or two-way ANOVA with post hoc independent pairwise analysis via Tukey’s multiple comparison test, Fisher’s LSD or others where appropriate using Prism 11 (GraphPad Software Inc, San Diego, CA). All p-values for GO evaluations were calculated by two-sided Fisher’s exact test using the Enrichr tool. Other statistical details of experiments can be found in the figure legends, including the statistical tests used and the number of replicate samples. Across all statistical tests, significance was defined as p-value <0.05. Unless otherwise specified, all graphs show mean values plus or minus standard error of the means (s.e.m.) as well as individual values overlaid as dot plots.

## Acknowledgments

We thank the many funding sources that contributed to this work including Boston University Start-Up Funds, Boston University’s Distinguished Summer Research Fellowship (DSRF), Boston University’s Undergraduate Research Opportunities Program (UROP), and the National Institute of Neurological Disorders and Stroke (NIH NINDS) F31NS135944 to E.M.D, F31NS145754 to L.F.H. and R21NS136831 to T.M.O. This research was supported by the Biomedical Engineering Core Facilities at Boston University (BU). A special thanks to Xin Brown and the Biointerface Technologies (BIT) core, as well as the Micro and Nano Imaging (MNI) core at BU. We also thank the Boston University Neurophotonics Center for providing access to the Bruker Ultima Investigator two-photon microscope used in this work. We appreciate the support of our funding sources and note that this content is solely the responsibility of the authors and does not reflect the official views of the National Institutes of Health (NIH).

## Author Contributions

**CRediT (Contributor Roles Taxonomy) roles:**

**Conceptualization -** E.M.D. + T.M.O.

**Data curation -** E.M.D. + K.Y. + P.K. + L.F.H. + H.A. + I.H. + T.M.O.

**Formal analysis -** E.M.D. + K.Y. + P.K. + L.F.H. + I.H. + T.M.O.

**Funding acquisition -** E.M.D. + P.K. + L.F.H. + I.H. + T.M.O.

**Investigation -** E.M.D. + K.Y. + P.K. + L.F.H. + H.A. + I.H. + K.D. + T.M.O.

**Methodology -** E.M.D. + K.Y. + P.K. + L.F.H. + H.A. + I.H. + K.D. + T.M.O.

**Project administration -** E.M.D. + T.M.O.

**Resources -** T.M.O.

**Software -** P.K. + T.M.O.

**Supervision -** E.M.D. + T.M.O.

**Validation -** E.M.D. + K.Y. + P.K. + L.F.H. + I.H. + T.M.O.

**Visualization -** E.M.D. + K.Y. + P.K. + L.F.H. + I.H. + T.M.O.

**Writing – original draft -** E.M.D. + K.Y. + P.K. + L.F.H. + T.M.O.

**Writing – review and editing -** E.M.D. + K.Y. + P.K. + L.F.H. + I.H. + T.M.O.

## Competing interests

Authors declare that they have no competing interests.

## Data and materials availability

Raw FASTQ sequencing files and processed count data have been deposited at Gene Expression Omnibus (GEO), hosted by the National Center for Biotechnology Information (NCBI), and are publicly available with accession number GSE344979. All other data needed to evaluate the conclusions in the paper are present in the paper and/or the Supplementary Materials. Other data that support the findings of this study, or the materials used in the study, are available on reasonable request from the corresponding author.

## Notes

### Competing Interest Statement

The authors have declared no competing interest.

