## Supplementary Information for "Profiling and modulating astrocyte borders at injected biomaterials in mice"

#### Supplementary Tables

**See Attached Table File for all Supplementary Tables.**

### Supplementary Figures

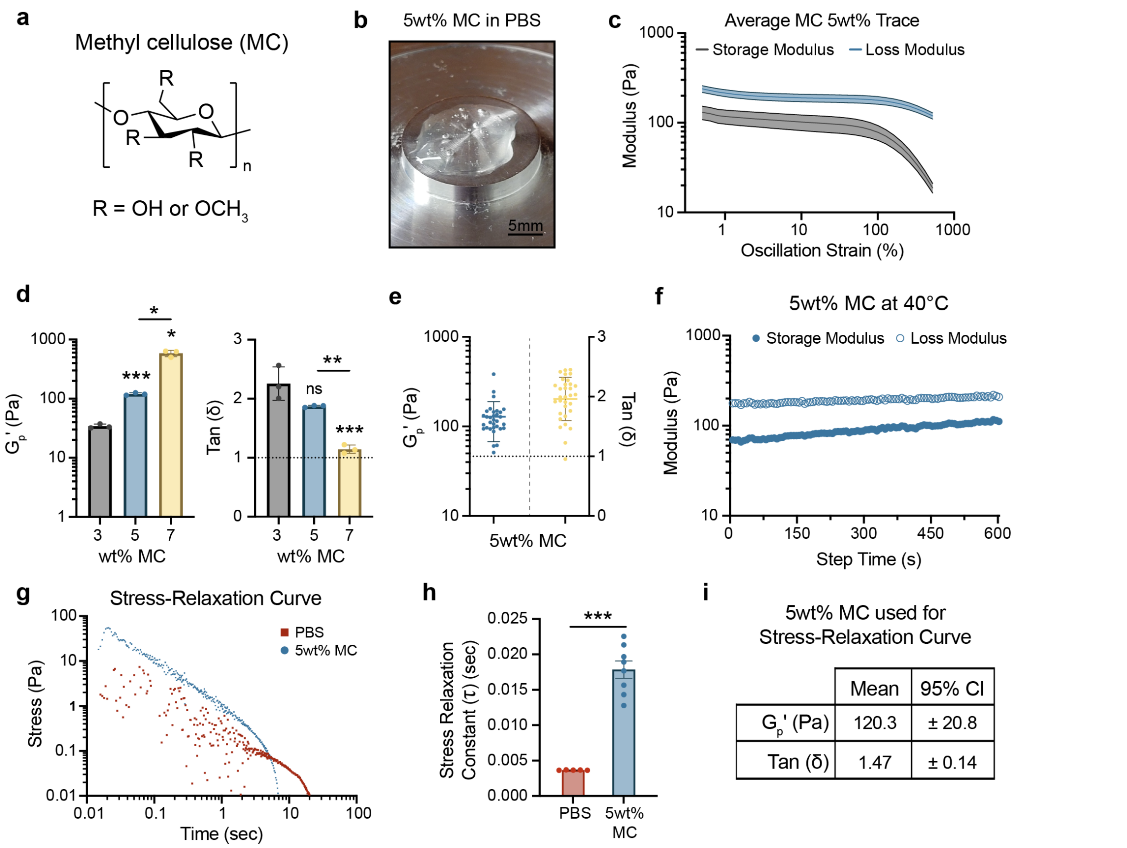


#### Supplementary Fig. 1 | MC forms a viscoelastic deposit with low variability in material properties.

**a.** Chemical structure of methyl cellulose (MC). **b.** Image showing 5 wt% MC in 1x phosphate buffered saline (PBS) deposited on rheometer baseplate. **c.** Average dynamic rheological oscillatory strain sweep measurements for 5wt% MC showing storage modulus, (G′) and loss modulus, (G″). Graphs show mean ± 95% Confidence Interval (CI), with 95% CI represented as light shaded areas (n=32). **d.** Bar chart showing plateau storage modulus ($G_p'$) and tan(∂) at 3 wt%, 5 wt%, and 7wt% MC. Graph shows mean ± standard deviation (SD) (n=3), with not significant (ns), $^*P < 0.02$, $^{**}P < 0.005$ and $^{***}P < 0.0006$ compared to 3 wt% or as designated, via Brown-Forsythe and Welch ANOVA with Dunnet's T3 Test ($G_p'$) or one-way ANOVA with Tukey's multiple comparison test (Tan(∂)). **e.** Plot showing plateau storage modulus ($G_p'$) and tan(∂) for replicates of 5 wt% MC, prepared under surgery-like conditions. Graph shows mean ± SD (n=32). **f.** Representative dynamic rheological oscillatory measurements for 5wt% MC at 40°C and 0.5% oscillation strain, showing storage modulus, (G′) and loss modulus, (G″). **g.** Mean stress relaxation curves for PBS (n=5) and 5wt% MC (n=8). **h.** Comparison of stress relaxation time constant (τ) for PBS (n=5) and 5wt% MC (n=8). Plot shows $^{***}P < 0.0001$ via Welch's t-test. **i.** Table showing characteristic properties of 5wt% MC gel used in stress relaxation test (n=8).

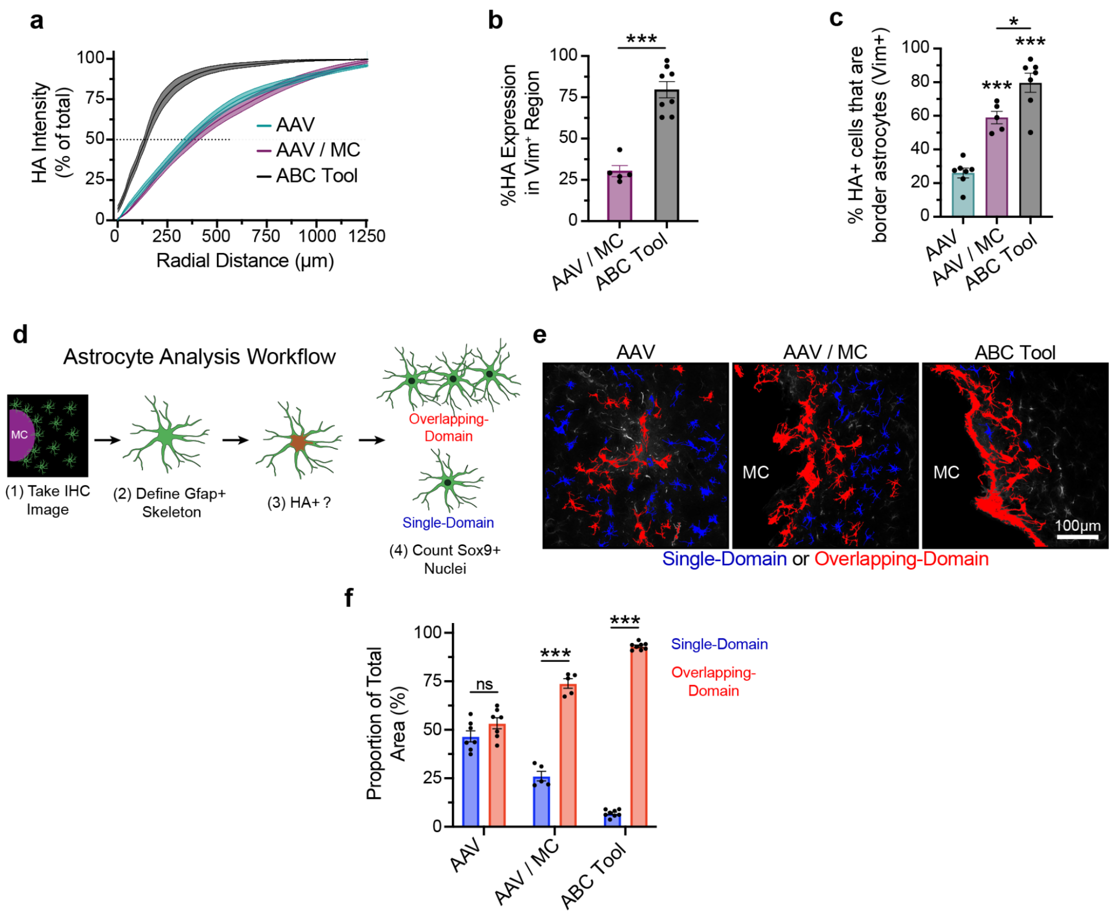


#### Supplementary Fig. 2 | MC deposits localize HA expression and create morphologically unique border phenotypes.

**a.** Quantification of HA intensity measured radially from the material surface and normalized to total HA expression. Graphs show mean ± s.e.m., with s.e.m. represented as light shaded areas. **b.** Bar chart showing the % of total HA expression located within the Vimentin (Vim)-positive Region. **c.** Percentage of HA+ astrocytes that stain positive for Vimentin (Vim) within 250 µm of MC surface. **d.** Schematic showing Astrocyte Analysis Workflow for (**e-f**), used to assess the relative contribution of single-domain and overlapping-domain astrocytes to the glial response to AAV or MC injection. Using a custom analysis workflow performed across ImageJ and MATLAB, we assessed the differences in astrocyte phenotype between AAV solution alone and MC-containing conditions. By isolating Gfap-positive skeletons of astrocytes within the first 250 µm, which also stain positive for HA-RiboTag and Sox9 (an astrocyte specific transcription factor), we were able to perform a morphological analysis. The predominant morphological characteristic that emerged from this analysis was the transition of astrocytes from single-domain (SD) to overlapping-domain (OD) cells. **e.** Detail images that have been processed using the Astrocyte Analysis Workflow and pseudocolored to show single-domain (blue) and overlapping-domain astrocytes (red) adjacent to the injection site. **f.** Quantification of proportion (by area) of single-domain and overlapping-domain HA-positive astrocytes contributing to glial response, with AAV alone having an approximate 1:1 area ratio of SD:OD cells, which shifted dramatically to 1:13 for the ABC Tool. All statistical comparisons shown compare single-domain to overlapping-domain in a given condition. All bar charts in this figure show not significant (ns), $^{*}P < 0.02$, and $^{***}P < 0.0004$ compared to AAV or as designated, via unpaired t-test (**b**), one-way ANOVA with Tukey's multiple comparison test (**c**) or two-way ANOVA with Tukey's multiple comparison test (**f**). Graph shows mean ± s.e.m. with individual data points showing AAV (n = 7), AAV / MC (n = 5), and ABC Tool (n = 8 or n=7 (**c**)).

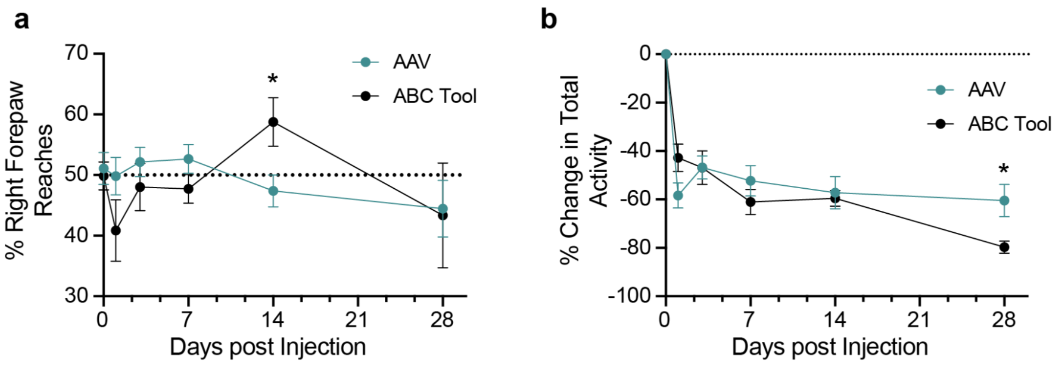


**Supplementary Fig. 3 | Behavioral and functional assessment of mice post-surgery using the cylinder test.**

**a.** Plot showing percent of forepaw reaches performed using right forepaw for AAV and the ABC Tool at 0-28 days post injection (dpi). **b.** Plot showing percent change in total mouse activity for AAV and the ABC Tool at 0-28 days post injection (dpi). All plots in this figure show *P < 0.04 with comparisons between AAV and the ABC Tool via RM two-way ANOVA with Geisser-Greenhouse correction and assessed with Tukey's multiple comparison test. Graph shows mean ± s.e.m. with individual data points showing (n = 10) for AAV and the ABC Tool.

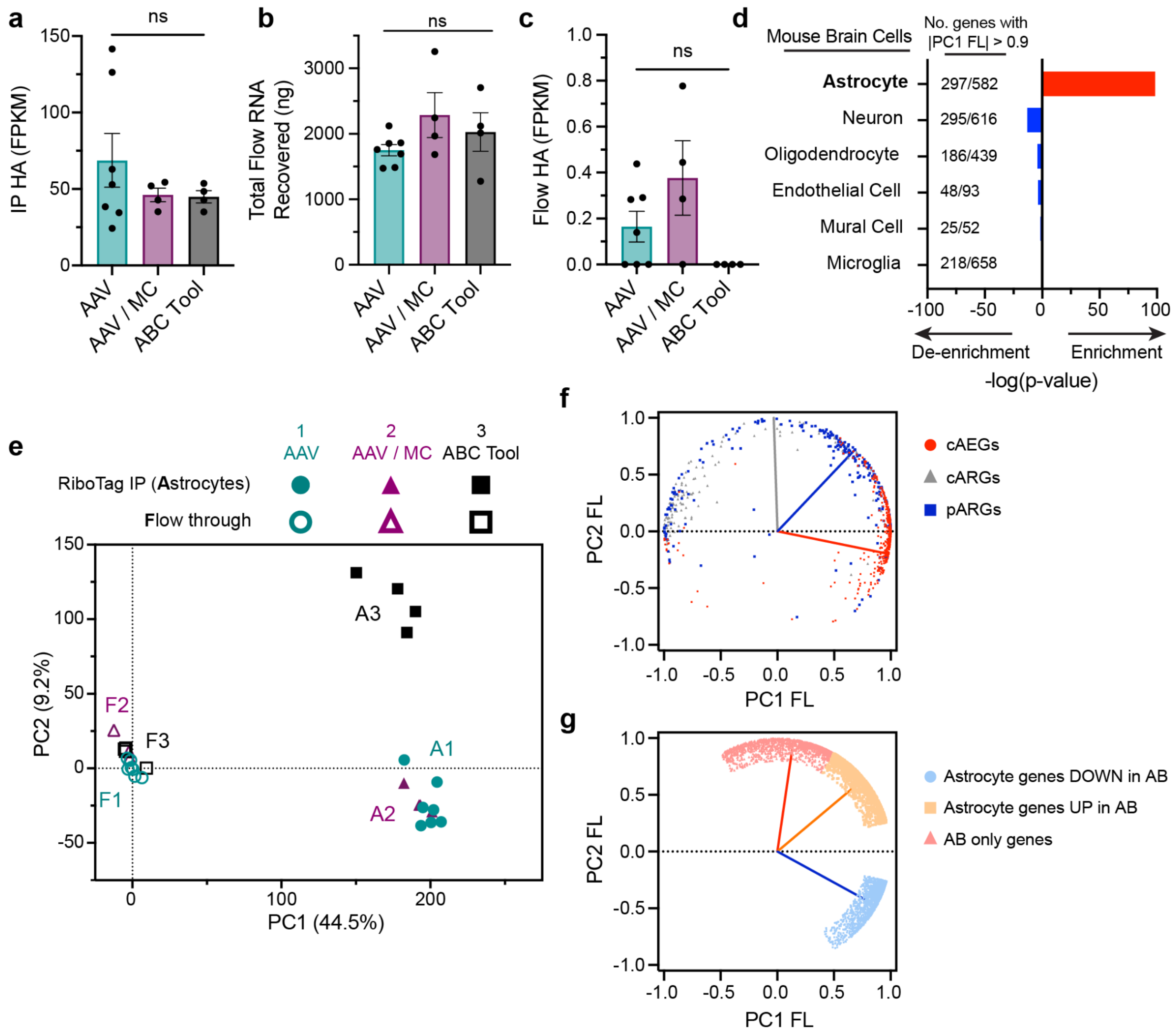


**Supplementary Fig. 4 | The ABC Tool enables high enrichment of astrocyte transcripts with enhanced border-specific gene identification.**

**a.** HA fragments per kilobase of transcript per million mapped reads (FPKM) for IP samples. **b.** Total RNA recovered from the Flow process for all conditions. **c.** HA FPKM for flow samples. **d.** Cell Marker database analysis performed on all PC1 genes with a factor loading (FL) > 0.9 or FL < -0.9. Plot shows the significance and relative overlap of identified genes with major brain cell types. **e.** PCA analysis of FPKM values of DEGs on a per animal basis. **f.** Plot of factor loading values in PC1 and PC2 for genes from previously identified, curated lists for consensus healthy astrocyte-enriched genes (cAEGs), consensus astrocyte reactivity genes (cARGs), and persisting astrocyte reactivity genes (pARGs). **g.** Plot of factor loading values in PC1 and PC2 for genes from newly identified, curated lists for astrocyte genes down in AB, astrocyte genes up in AB, and AB-only genes. All bar charts in this figure (**a-c**) show not significant (ns) compared to AAV or as designated. One-way ANOVA with Tukey's multiple comparison test. Graph shows mean ± s.e.m. with individual data points showing IP: AAV (n = 7), AAV / MC (n = 4), and ABC Tool (n = 4). Flow: AAV (n = 7), AAV / MC (n = 4), and ABC Tool (n = 4). PCA plot is generated from the same sample set.

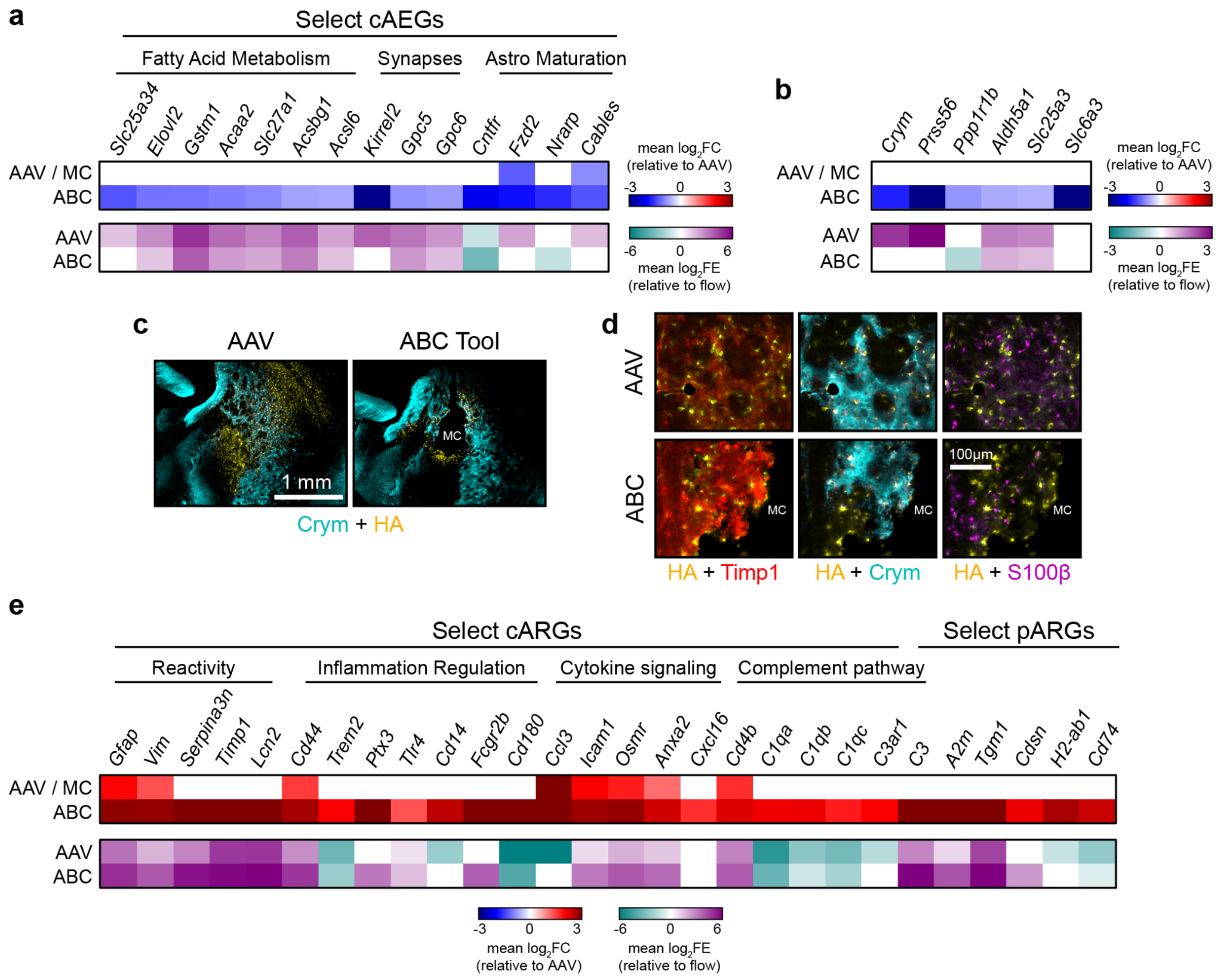


#### Supplementary Fig. 5 | DEGs for healthy and reactive astrocyte-specific molecules.

**a.** Top heatmap shows DEGs for the AAV/MC and ABC Tool condition and bottom heatmap shows mean enrichment in IP samples relative to Flow for the AAV and and ABC Tool conditions for **a.** select DEGs from the cAEGs list. **b.** Select DEGs for canonical striatal region-specific astrocyte molecules. **c.** Overview and **d.** detail images showing colocalization of Crym in Timp1+ border astrocytes **e.** Select DEGs from the cARGs and pARGs list. All transcriptomic data shown in this figure is derived from the following sequenced sample set: IP: AAV (n = 7), AAV / MC (n = 4), and ABC Tool (n = 4). Flow: AAV (n = 7), AAV / MC (n = 4), and ABC Tool (n = 4).

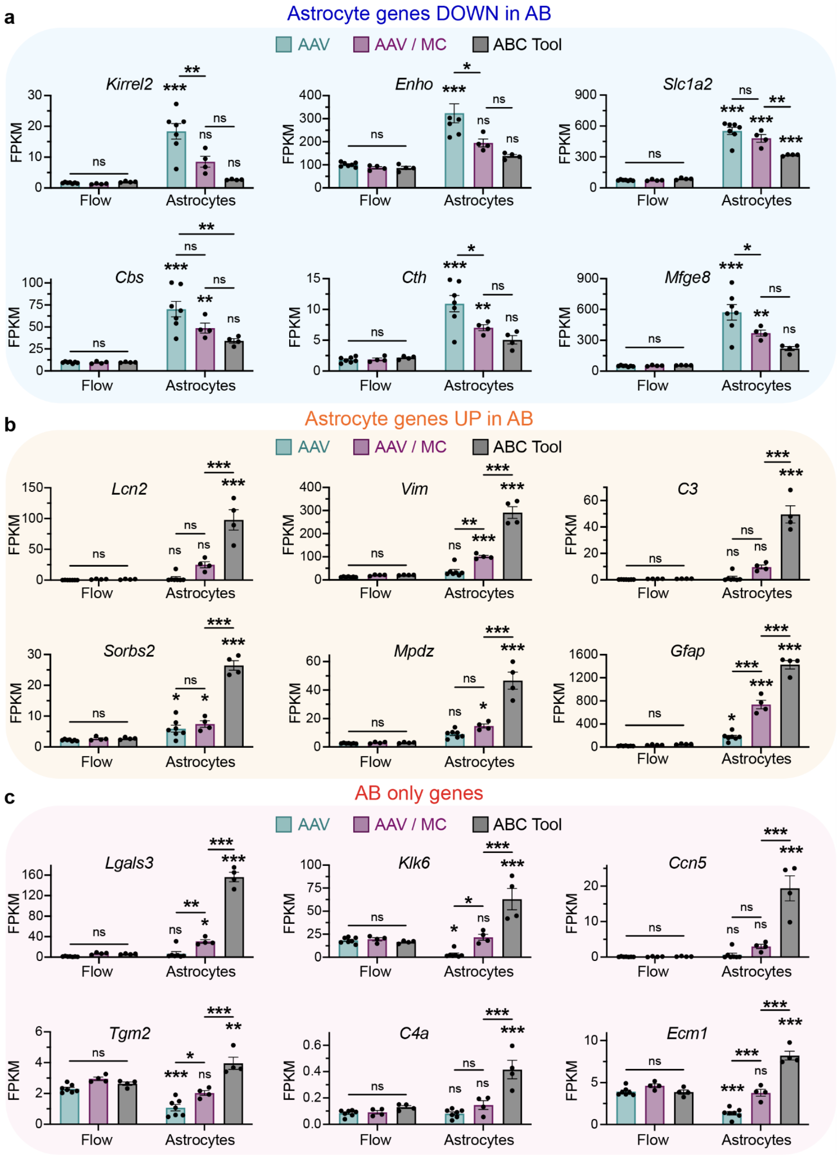


**Supplementary Fig. 6 | FPKM plots showing expression levels of astrocyte genes down in AB, astrocyte genes up in AB, and AB-only genes.**

Plots showing FPKM values for IP and flow samples for selected **a.** astrocyte genes down in AB, **b.** astrocyte genes up in AB, and **c.** AB-only genes. All bar charts in this figure show not significant (ns), *P < 0.05, **P < 0.009 and ***P < 0.0007 compared to matched flow sample or as designated. Two-way ANOVA with Tukey's multiple comparison test. Graph shows mean ± s.e.m. with individual data points showing IP: AAV (n = 7), AAV / MC (n = 4), and ABC Tool (n = 4). Flow: AAV (n = 7), AAV / MC (n = 4), and ABC Tool (n = 4).

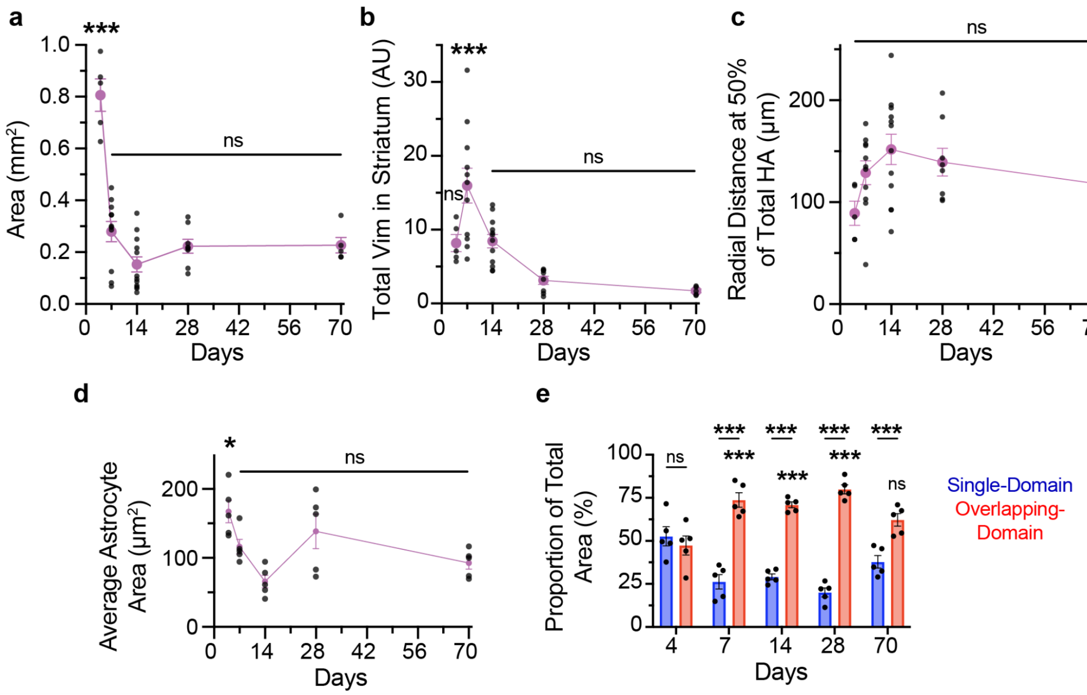


**Supplementary Fig. 7 | MC interface matures over time, resulting in less total vimentin, lower surface area, and decreased circularity.**

**a.** Trace showing MC deposit area ($mm^2$) over time as defined by 4′,6-diamidino-2-phenylindole (DAPI) staining. **b.** Trace showing total Vimentin expression over time, calculated as area under the curve (AUC) up to 1000 μm from the center of the injection. **c.** Trace showing radial distance at 50% of HA expression over time. **d.** Trace showing average astrocyte area (area of Gfap+ skeletons normalized by Sox9+ nuclei) up to 100μm into tissue from MC surface over time. **e.** Quantification of proportion (by area) of single-domain and overlapping-domain HA-positive astrocytes within 250μm of MC deposit across time. All x-y plots in this figure show not significant (ns), *$P < 0.03$, and ***$P < 0.0008$ compared to 70d or as designated via a one-way ANOVA with Tukey's multiple comparison test (**a-d**) or compared to 4d or as designated via a two-way ANOVA with Tukey's multiple comparison test (**e**). Graphs show mean ± s.e.m. with individual data points showing: (**a-c**) 4d (n = 5), 7d (n = 11), 14d (n = 12), 28d (n = 8), and 70d (n = 5). (**d-e**) n=5 for all timepoints.

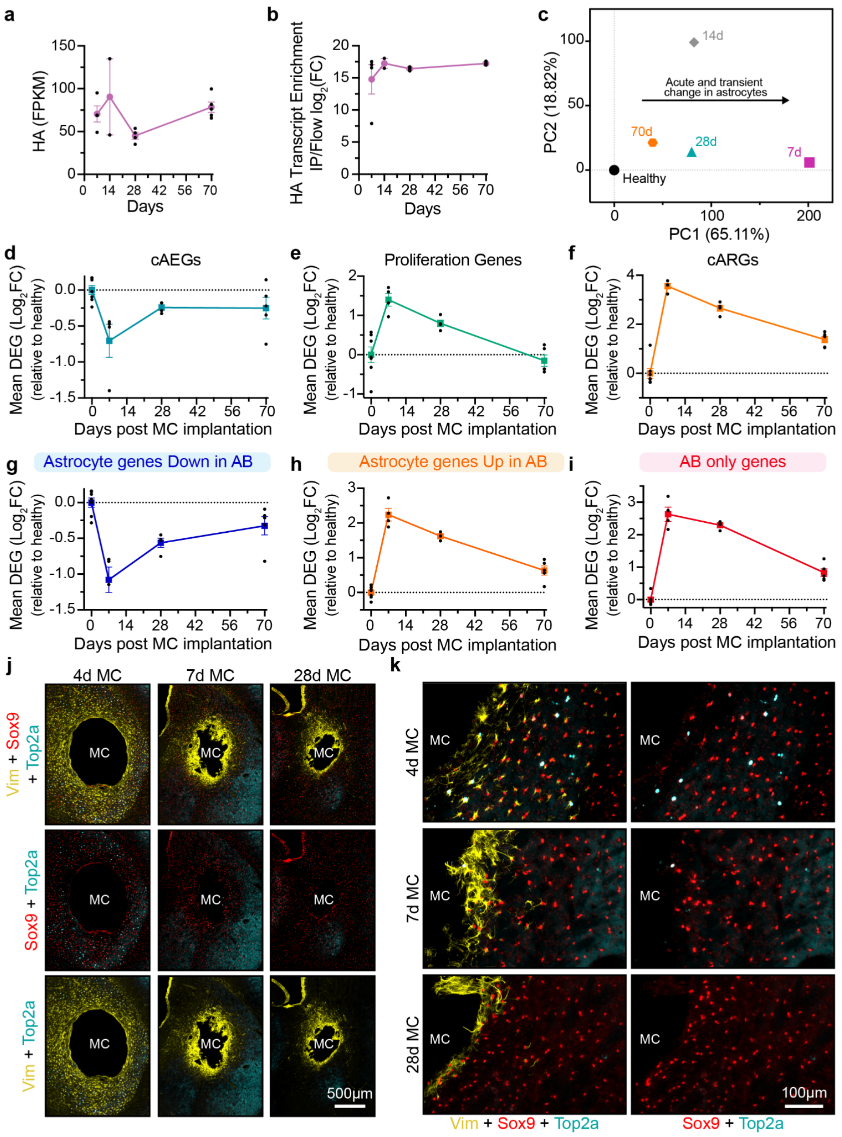


### Supplementary Fig. 8 | Assessment of temporal transcriptomics reveals constant HA expression over time and transient, temporally regulated changes in gene expression.

**a.** HA FPKM values for IP samples at all timepoints. **b.** Relative enrichment of 3xHA in RiboTag-IP samples relative to flow samples at all timepoints. **c.** PCA analysis of average FPKM values of DEGs on a per timepoint basis. **d.-f.** Mean differential expression ($log_2(FC)$) over time of a curated list of previously identified, **d.** consensus healthy astrocyte-enriched genes (cAEGs), **e.** proliferation genes, and **f.** consensus astrocyte reactivity genes (cARGs). **g.-i.** Mean differential expression ($log_2(FC)$) over time of a curated list of newly identified, **g.** astrocyte genes down in AB, **h.** astrocyte genes up in AB, and **i.** AB-only genes. **j.** Overview and **k.** Detail images showing astrocyte proliferation at 4, 7, and 28d following MC implantation. All x-y plots in this figure show mean ± s.e.m. Individual data points in (**a-b**) show 7d (n = 4), 14d (n = 2), 28d (n = 4), and 70d (n = 5). PCA plot is generated from the same sample set. Individual data points in (**d.-i.**) show 0d (Healthy) (n = 7), 7d (n = 4), 28d (n = 4), and 70d (n = 5).

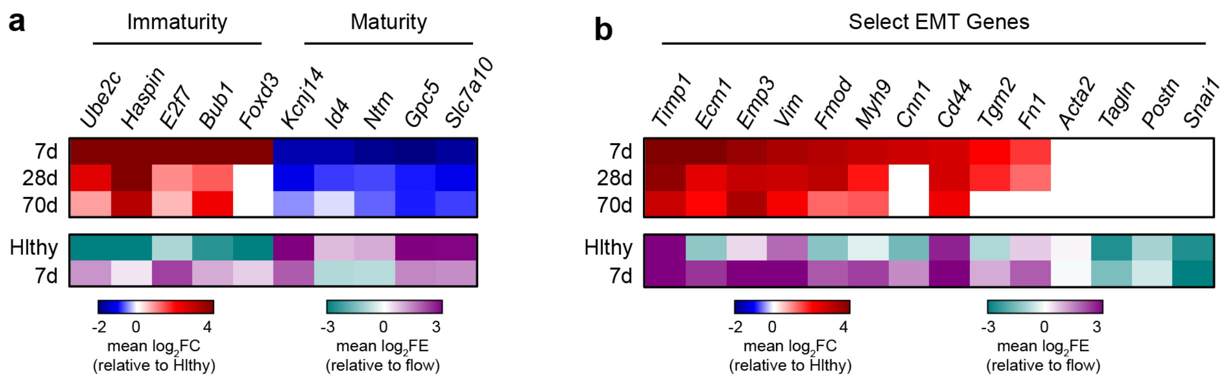


**Supplementary Fig. 9 | Temporally regulated DEGs for Immaturity/Maturity and EMT.**

Heatmaps of $log_2FC$ and $log_2FE$ for **a.** Immaturity/Maturity and **b.** EMT-related genes at designated timepoints post MC biomaterial injection. Plots show Healthy (Hlthy) (n = 7), 7d (n = 4), 28d (n = 4), and 70d (n = 5).

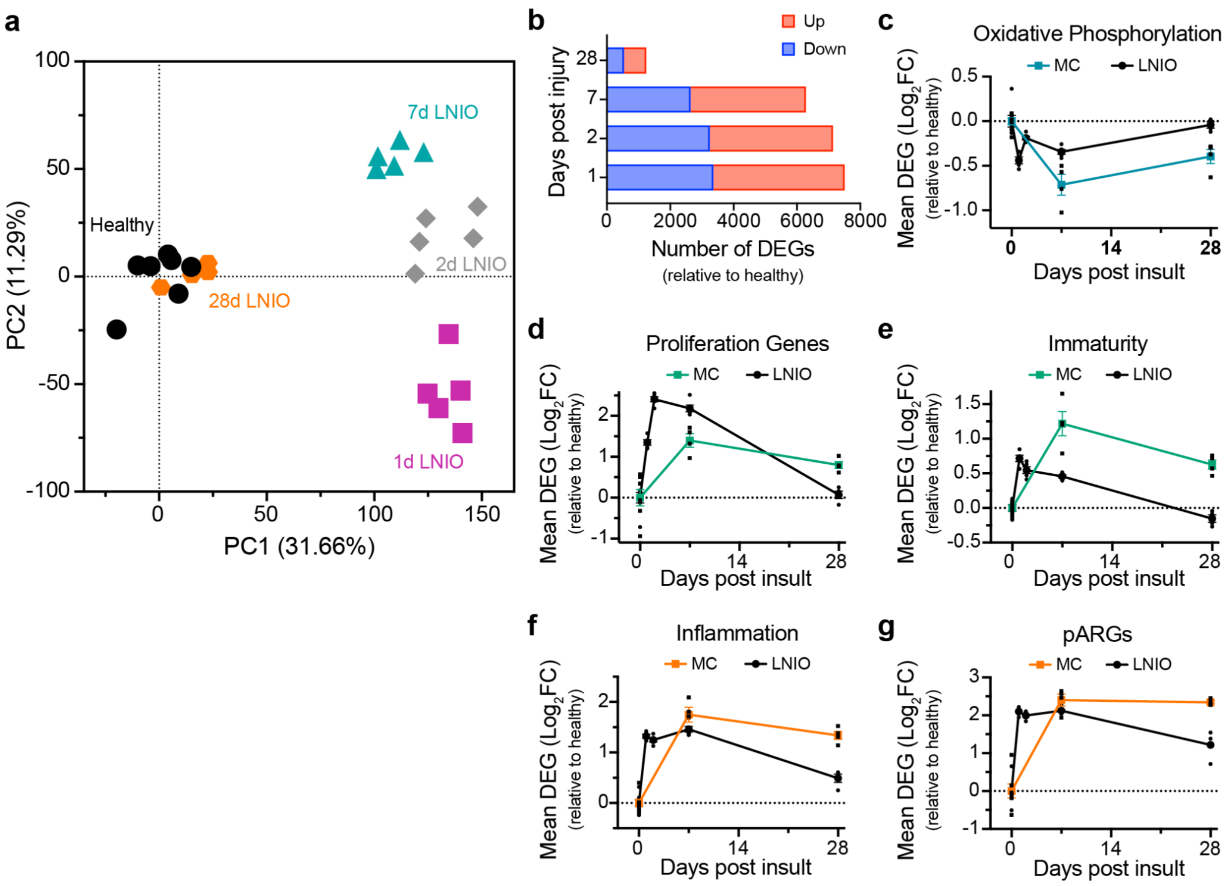


**Supplementary Fig. 10 | Ischemic stroke results in similar changes in transcriptome, causing a temporal regulation of metabolism, proliferation, and wound resolution-related genes.**

**a.** PCA analysis of FPKM values of DEGs on a per animal basis. **b.** Bar chart showing the number of DEGs at each timepoint relative to Healthy (AAV IP). **c.-g.** Mean differential expression ($log_2$(FC)) over time of a curated list of previously identified **c.** oxidative phosphorylation-related genes. **d.** proliferation genes. **e.** immaturity and maturity-related genes (immaturity score is assessed as the difference between immaturity and maturity-related genes) **f.** inflammation genes, and **g.** persisting astrocyte reactivity genes (pARGs). All x-y plots in this figure show mean ± s.e.m. with individual data points showing 0d or Healthy (n = 7), 7d (n = 4), 28d (n = 4) for MC and 1d (n = 5), 2d (n = 5), 7d (n = 5), 28d (n = 4) for LNIO. PCA plot and DEG data is generated from the same LNIO sample set.

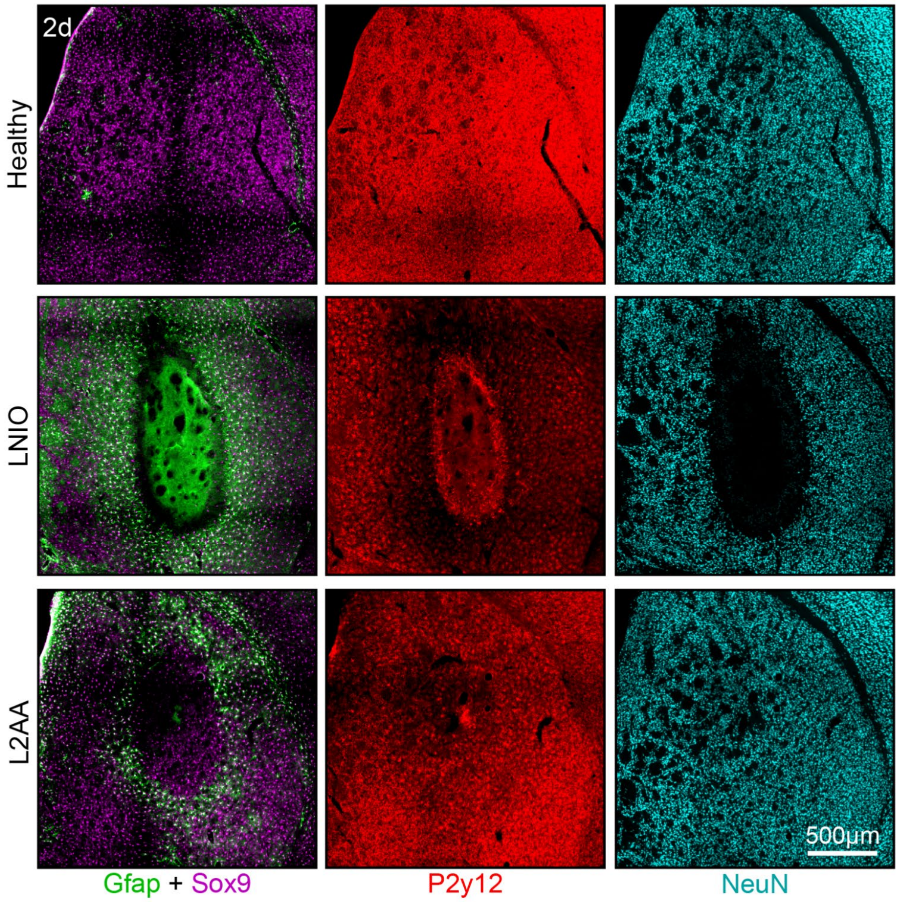


**Supplementary Fig. 11 | LNIO solution injection results in ablation of both glial cells and neurons while L2AA solution injection results in astrocyte-specific ablation.**

Overview images showing relative distribution of glial cells and neurons in healthy tissue or at 2d post-injection of an LNIO (27.4 mg/mL) or L2AA (20 mg/mL) solution.

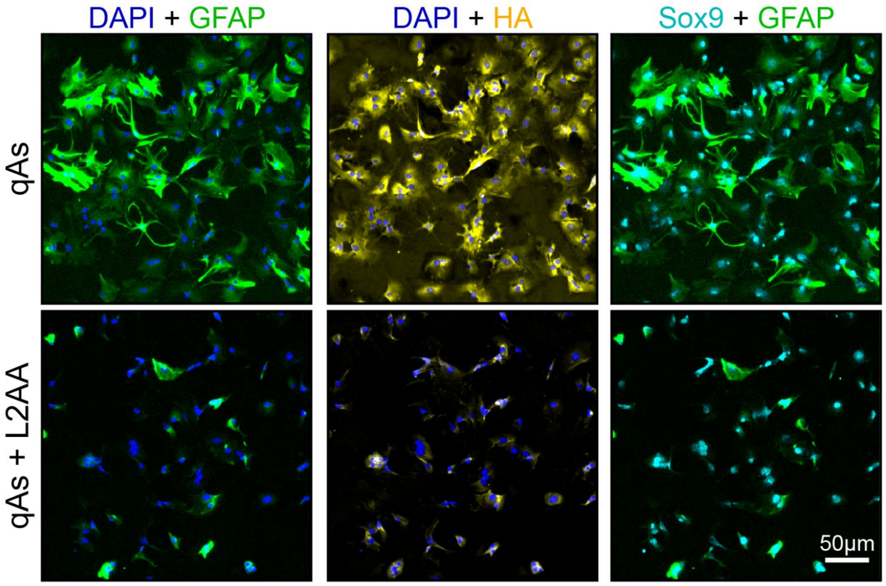


**Supplementary Fig. 12 | Astrocytes exposed to L-2-Aminoadipic Acid (L2AA) dysregulate ribosome and Gfap production.**

ICC staining showing quiescent astrocytes after 2 days in L2AA-free or L2AA-containing media stained for DAPI, Gfap, HA, and Sox9.

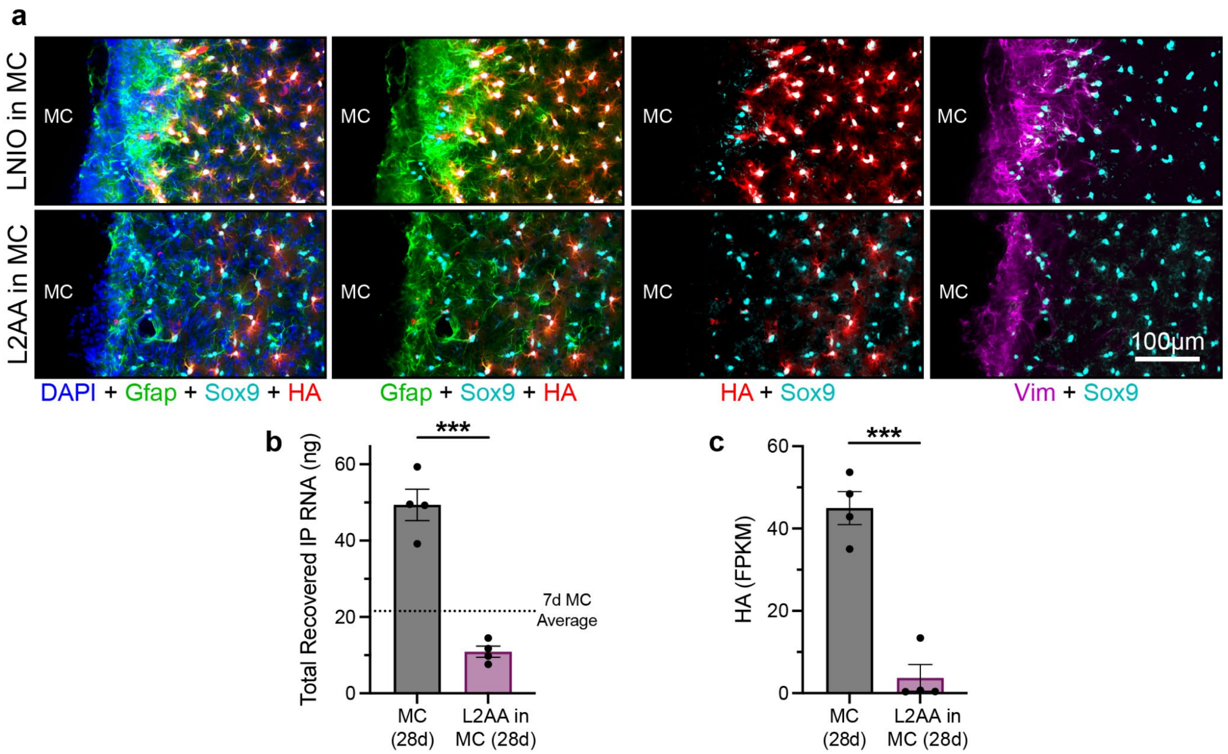


**Supplementary Fig. 13 | Release of LNIO and L2AA from MC deposits results in thick astrocyte borders and decreased HA expression at the material-tissue interface.**

**a.** Detail images showing astrocyte border formation and HA localization for LNIO- and L2AA-containing MC deposits at 28 dpi. **b.** Total RNA recovered from the IP process for MC and L2AA in MC conditions. **c.** HA fragments per kilobase of transcript per million mapped reads (FPKM) in MC and L2AA in MC IP samples. All bar charts in this figure show ***$P < 0.0003$ as designated via unpaired t-test. Graph shows mean ± s.e.m. with individual data points showing MC ($n = 4$) and L2AA in MC ($n = 4$).

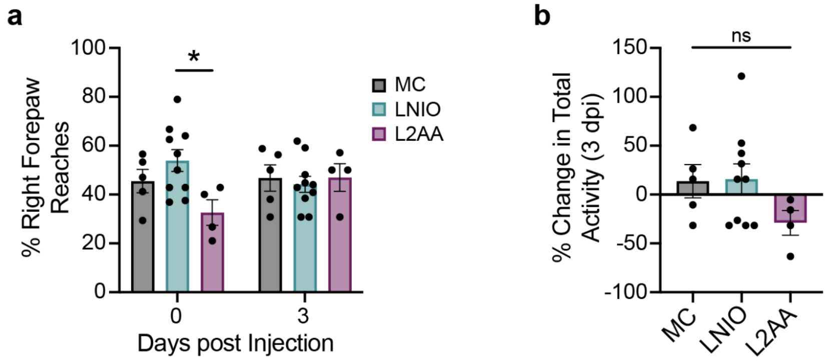


**Supplementary Fig. 14 | Behavioral and functional assessment of mice post implantation of unloaded and LNIO- or L2AA-loaded MC deposits using the cylinder test.**

**a.** Plot showing percent of forepaw reaches performed using right forepaw for MC, LNIO, and L2AA at 0 and 3 days post injection (dpi). **b.** Plot showing percent change in total mouse activity for MC, LNIO, and L2AA at 3 dpi. All plots in this figure show not significant (ns) and $^{*}P < 0.02$ with comparisons as designated by either (**a.**) repeated measures (RM) two-way ANOVA or (**b.**) one-way ANOVA with Tukey's multiple comparison test. Graph shows mean ± s.e.m. with individual data points showing MC (n = 5), LNIO (n = 10), and L2AA (n = 4).

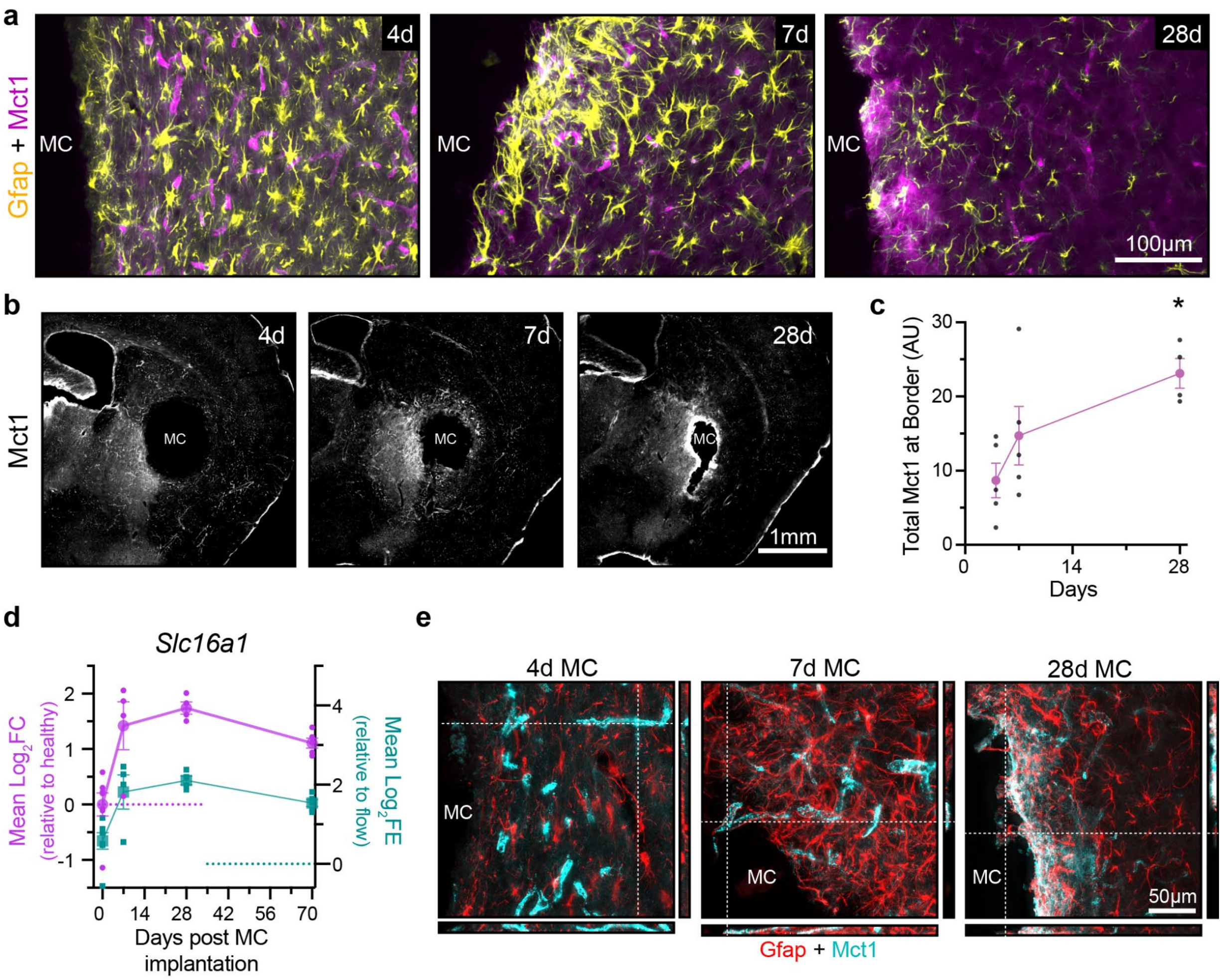


**Supplementary Fig. 15 | Astrocytes express Mct1, a transporter for ketone bodies, with expression increasing in response to MC biomaterial stimulus.**

**a.** Detail images showing the emergence of MCT1 and colocalization with Gfap over time. **b.** Overview images showing the emergence of MCT1 over time. **c.** Quantification of total MCT1 expression at the border (<250µm into tissue from the start of the deposit) over time. Plot shows *P < 0.02 compared to 4d or as designated, one-way ANOVA with Tukey's multiple comparison test. Graph shows mean ± s.e.m. with individual data points showing 4d (n = 5), 7d (n = 5), and 28d (n = 4). **d.** Plot showing mean $log_2FC$ and $log_2FE$ for *Slc16a1* expression (gene for Mct1) over time. Plot shows mean ± s.e.m. with individual data points showing 0d or Healthy (n = 7), 7d (n = 4), 28d (n = 4), and 70d (n = 5). **e.** 3D orthogonal projections showing changes in MCT1 expression in astrocytes at MC border over time.

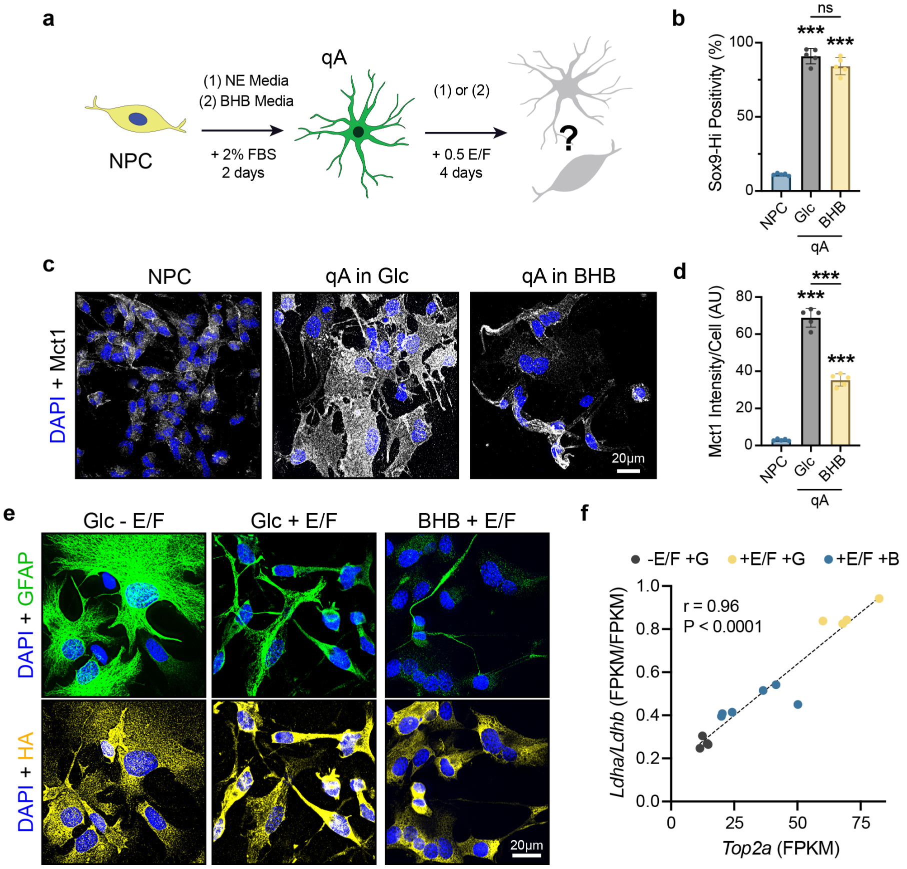


#### Supplementary Fig. 16 | Astrocytes express Mct1 *in vitro* and modulate Gfap expression and organization in response to BHB treatment.

**a.** Schematic showing conditioning scheme for converting neural progenitor cells (NPCs) into quiescent astrocytes (qAs) in either (1) neural expansion (NE) or (2) BHB-containing media and testing proliferative capacity by 4-day exposure to 50 ng/mL E/F in either NE (Glc) or BHB media. **b.** Bar chart quantifying high intensity Sox9 positivity in NPCs vs qAs derived in Glc or BHB media. **c.** Detail images showing Mct1 staining in NPCs vs qAs derived in Glc or BHB media **d.** Bar chart quantifying Mct1 staining in NPCs vs qAs derived in Glc or BHB media. **e.** Detail images showing Gfap and HA-RiboTag distribution in qAs derived in NE media vs qAs treated with glucose or BHB and E/F. **f.** xy-plot showing correlation between Top2a expression and the Ldha/Ldhb ratio. Plot shows the line of best fit for a simple linear regression. r- and P- values are derived from Pearson's Correlation Test, ±E/F +Glu (n = 4) and +E/F +BHB (n = 6). Plots **b** and **d** show not significant (ns) and ***P < 0.0001 compared to NPC or as designated. One-way ANOVA with Tukey's multiple comparison test. Graph shows mean ± s.e.m. (n=5).

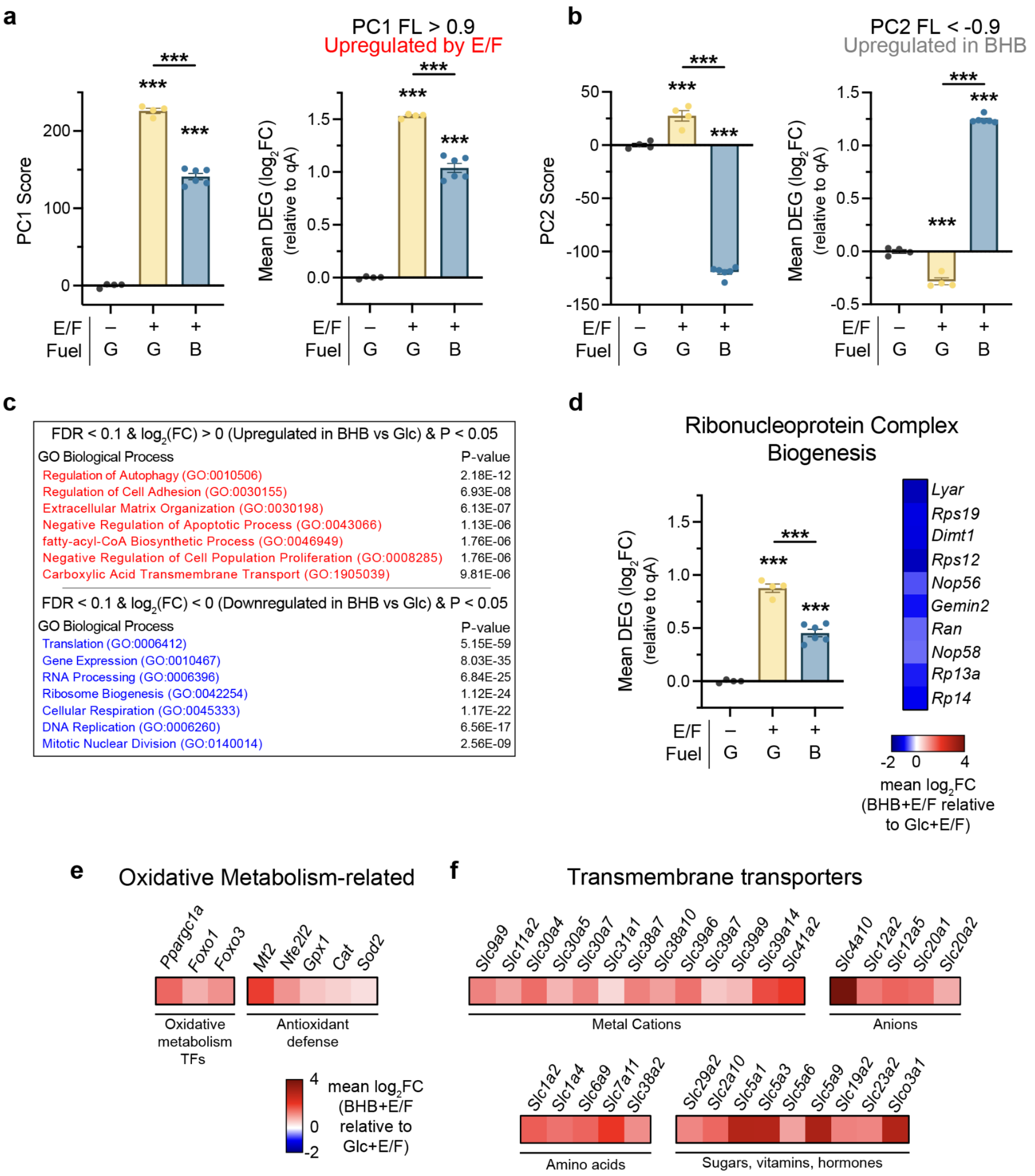


#### Supplementary Fig. 17 | Additional characterization of BHB treatment effect on astrocytes *in vitro*.

Bar charts showing PC1 Score and mean $log_2FC$ of genes defining **a.** the effect of E/F mitogen stimulation (PC1 FL > 0.9) and **b.** BHB-specific DEGs (PC2 FL < -0.9) for qA and E/F mitogen stimulated astrocytes in Glc- or BHB-media. **c.** GO-BPs identified by unbiased evaluations of DEGs defined by comparing proliferating astrocytes fueled by BHB to those fueled by glucose. DEGs must meet criteria of an FDR < 0.1, $log_2FC$ either >0 (upregulated in BHB vs Glc) or <0 (downregulated in BHB vs Glc), and a P-value of <0.05. **d**. Mean differential expression ($log_2FC$) across the 3 *in vitro* conditions with heatmaps showing representative fuel dependent DEGs for ribonucleoprotein complex biogenesis. **e**.-**f**. Heatmaps showing representative **e**. Oxidative Metabolism-related and **f**. Transmembrane transporter-associated DEGs. All bar charts show not significant (ns) and ***P < 0.0003 compared to the first column of data or as designated via one-way ANOVA with Tukey's multiple comparison test. Data is shown as mean ± s.e.m. with n=4 for ± E/F + G and n=6 for + E/F + B.

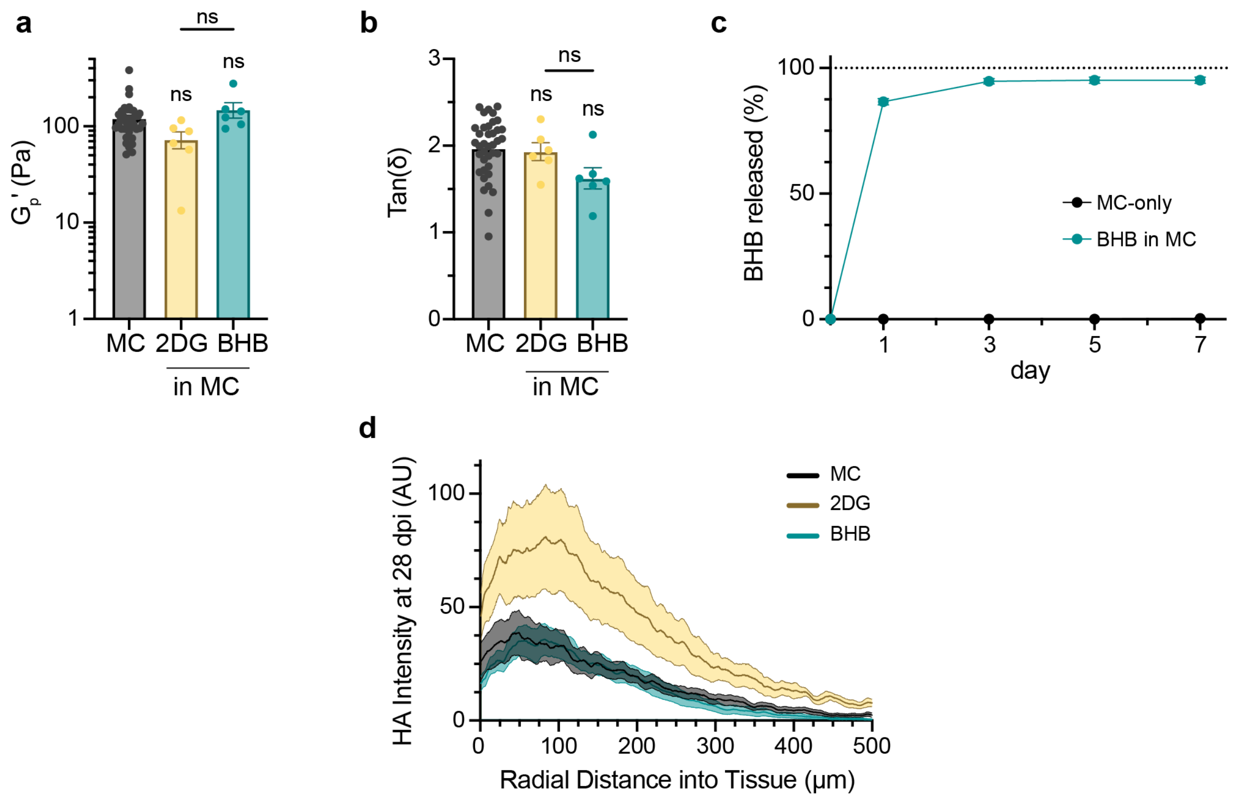


**Supplementary Fig. 18 | Incorporation of metabolic regulators into MC hydrogels results in no significant changes to material properties, rapid release, and no change in HA RiboTag distribution.**

Bar chart showing **a.** plateau storage modulus ($G_p'$) and **b.** tan($\partial$) of 5 wt% MC and 5wt% MC loaded with 20 mg/mL of 2-deoxy-d-glucose (2DG) and β-hydroxybutyrate (BHB). Plots show not significant (ns) across all samples via one-way ANOVA with Tukey's multiple comparison test. Graph shows mean ± s.e.m. (n=38 for MC, n=6 for 2DG and BHB). **c.** Graph showing 7 day cumulative release profile for BHB loaded at 20 mg/mL in a 5wt% MC gel and an MC-only control. BHB was released into PBS at 37°C. Graph shows mean ± s.e.m. (n=3). **d.** Plot of average HA intensity measured radially from the edge of the biomaterial deposit. Graph shows mean ± s.e.m. (shaded area) with individual data points showing: MC (n = 8), 2DG (n = 5), and BHB (n = 10).

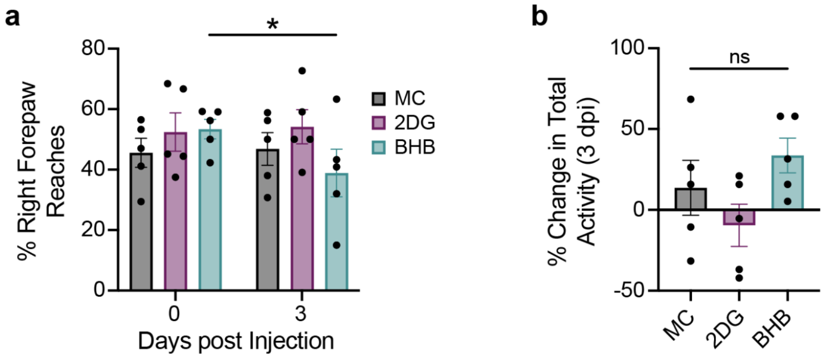


#### Supplementary Fig. 19 | Behavioral and functional assessment of mice post-implantation of unloaded and 2DG- or BHB-loaded MC deposits using the cylinder test.

**a.** Plot showing percent of forepaw reaches performed using right forepaw for MC, 2DG, and BHB at 0 and 3 days post injection (dpi). **b.** Plot showing percent change in total mouse activity for MC, 2DG, and BHB at 3 dpi. All plots in this figure show not significant (ns) and *P < 0.04 with comparisons as designated by either (**a.**) repeated measures (RM) two-way ANOVA or (**b.**) one-way ANOVA with Tukey's multiple comparison test. Graph shows mean ± s.e.m. with individual data points showing (n = 5) for MC, 2DG, and BHB.

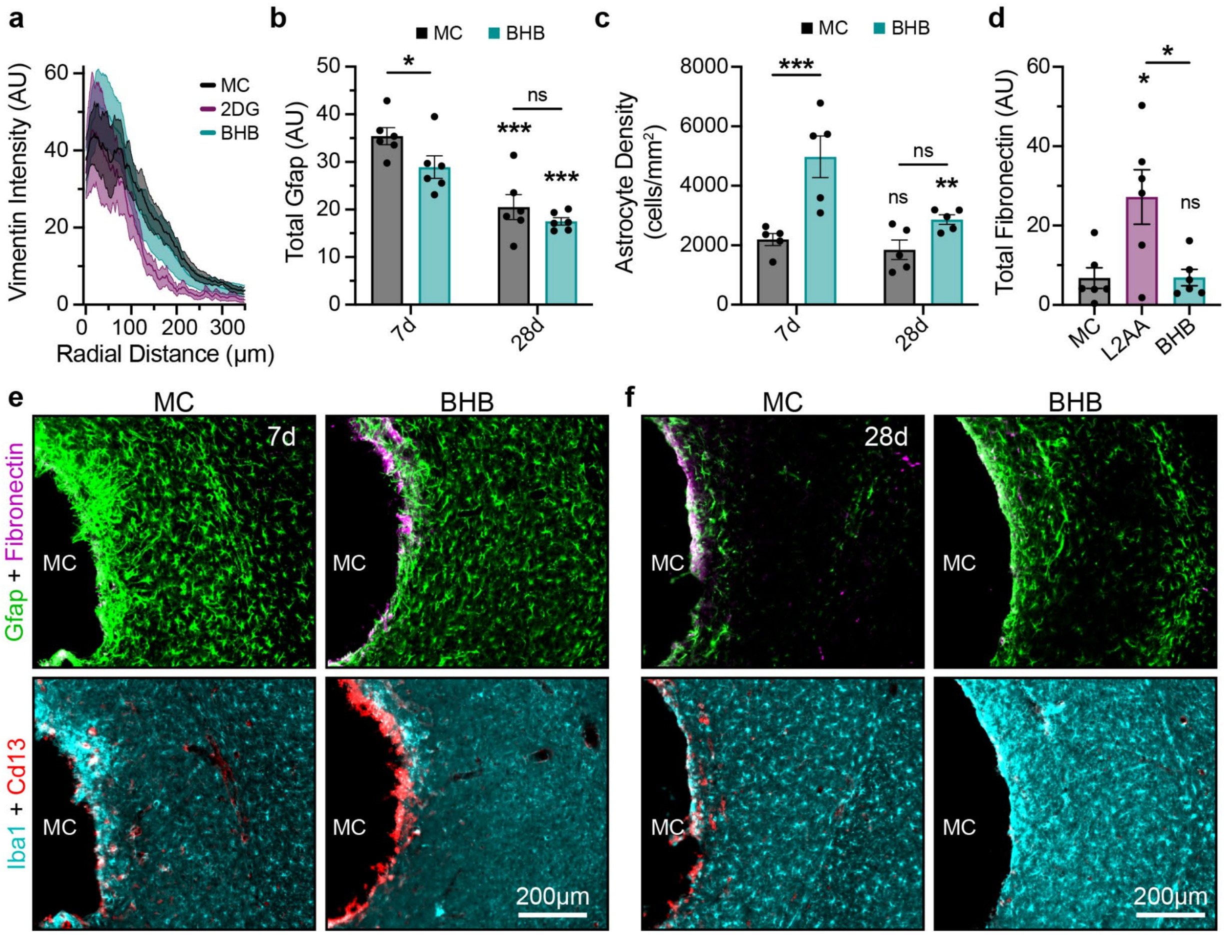


**Supplementary Fig. 20 | Releasing BHB from MC deposits *in vivo* results in astrocyte border differences, but minimal immune response.**

**a.** Vimentin intensity measured radially from the surface of the material. Graphs show mean ± s.e.m., with s.e.m. represented as light shaded areas. **b.** Quantification of total Gfap for 7 and 28 dpi MC and BHB + MC materials up to 300µm into the tissue. **c.** Plot showing Sox9+ astrocyte density up to 100µm into tissue at 7 and 28 dpi for BHB in MC and MC-only deposits. **d.** Quantification of total Fibronectin, calculated as area under the curve (AUC) up to 300µm into the tissue at 28 dpi. **e.-f.** Overview images showing fibrosis and inflammatory response at the astrocyte border at **e.** 7 dpi and **f.** 28 dpi for MC and BHB + MC materials. All bar charts in this figure show not significant (ns), *$P < 0.04$, **$P < 0.003$, and ***$P < 0.0008$. Either with comparisons made between timepoints within a group or as designated (**b-c**) via two-way ANOVA with Fisher's LSD test, or when compared to MC or as designated (**d**) via one-way ANOVA with Tukey's multiple comparison test. Graph shows mean ± s.e.m. with individual data points showing: 7d/28d MC (n = 6), 7d/28d BHB (n = 6) (**b, d**) or 7d/28d MC (n = 5), 7d/28d BHB (n = 5) (**c**).

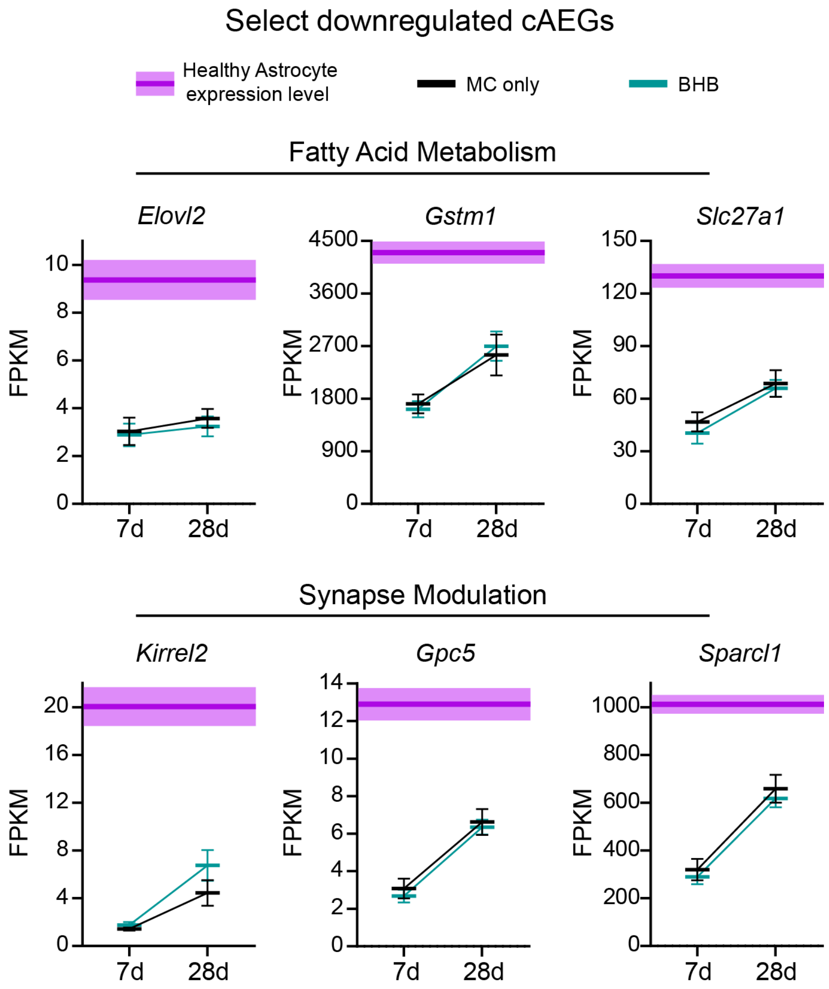


**Supplementary Fig. 21 | Releasing BHB from MC deposits *in vivo* results in conserved downregulation of cAEGs.**

FPKM values for select DEGs from the cAEG list across the MC-only and BHB-loaded biomaterial conditions at 7 and 28 dpi showing downregulation of Fatty Acid Metabolism- and Synapse Modulation-related functions. All graphs in this figure show mean ± s.e.m. with: Healthy (n=4), 7d/28d MC (n = 4), 7d BHB (n = 5), and 28d BHB (n = 4).

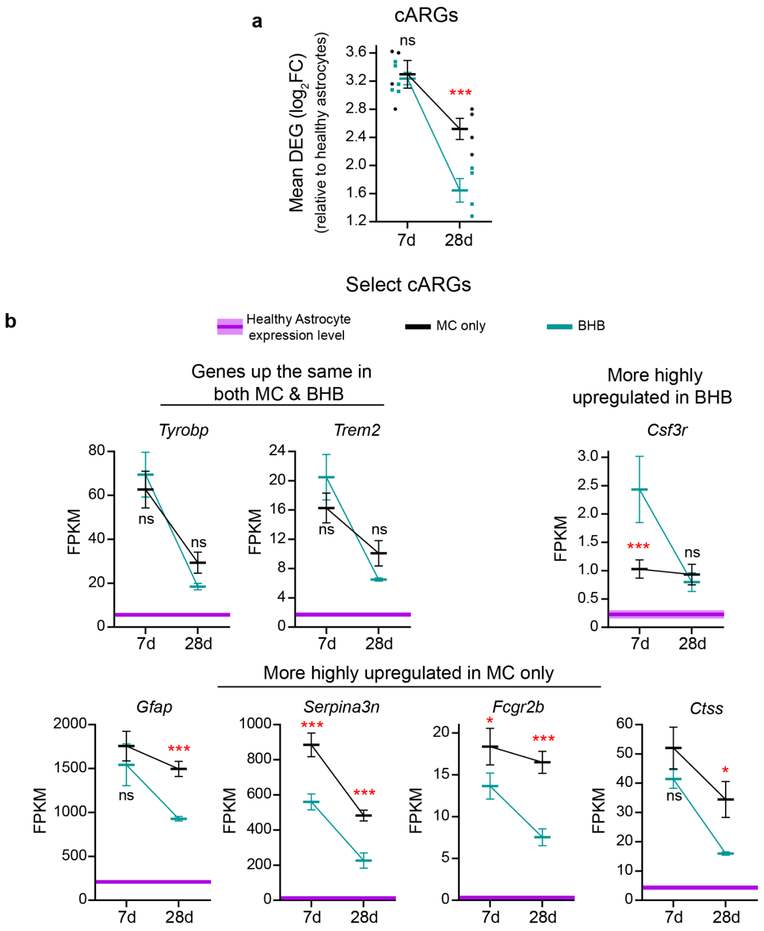


#### Supplementary Fig. 22 | Releasing BHB from MC deposits *in vivo* results in differential regulation of cARGs at 28 dpi.

**a.** Mean differential expression ($log_2FC$) across the MC-only and BHB-loaded biomaterials at 7 and 28 dpi showing DEGs for cARGs gene list. **b**. FPKM values for select DEGs from the cARG list across the MC-only and BHB-loaded biomaterial conditions at 7 and 28 dpi showing differential regulation of genes that are similarly up in both MC-only and BHB-containing conditions, more highly upregulated in BHB-containing conditions, and more highly upregulated in MC-only conditions. All graphs in this figure show not significant (ns), $^*P < 0.05$, and $^{***}P < 0.005$. Comparisons are made between groups at a single timepoint via two-way ANOVA with Tukey's multiple comparison test. Graph shows mean ± S.E.M (**a.-b.**). with individual data points (**a.**) showing: Healthy (n=4), 7d/28d MC (n = 4), 7d BHB (n = 5), and 28d BHB (n = 4).

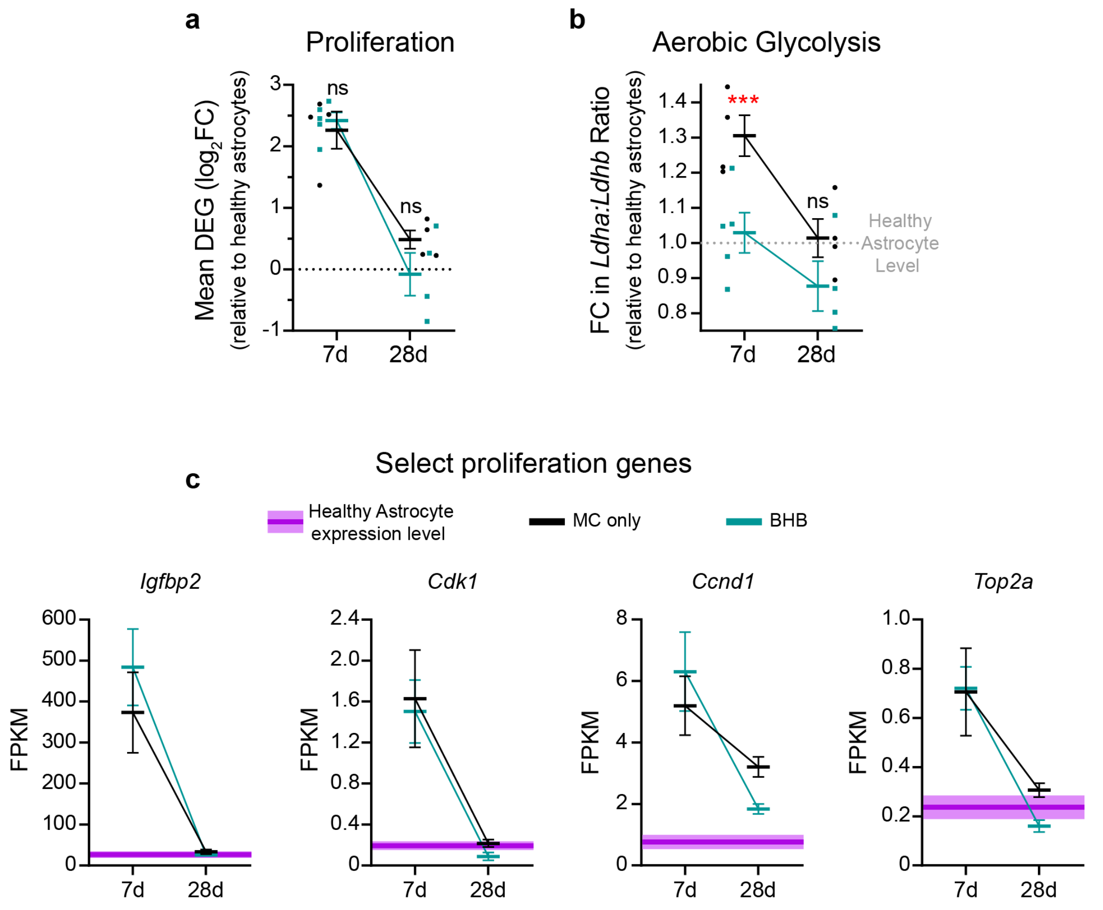


#### Supplementary Fig. 23 | Releasing BHB from MC deposits *in vivo* results in no change in proliferation-related genes, but differential regulation of Aerobic Glycolysis-associated genes at 7 dpi.

**a.** Mean differential expression ($log_2FC$) across the MC-only and BHB-loaded biomaterials at 7 and 28 dpi showing DEGs for the Proliferation-associated gene list. **b.** FC of select Aerobic Glycolysis-associated genes (ratio of Ldha/Ldhb expression). **c**. FPKM values for select DEGs from the Proliferation-associated list across the MC-only and BHB-loaded biomaterial conditions at 7 and 28 dpi. All graphs in this figure show not significant (ns), ***$P < 0.006$. Comparisons are made between groups at a single timepoint via two-way ANOVA with Tukey's multiple comparison test. Graph shows mean ± S.E.M (**a.-c.**). with individual data points (**a.-b.**) showing: Healthy (n=4), 7d/28d MC (n = 4), 7d BHB (n = 5), and 28d BHB (n = 4).

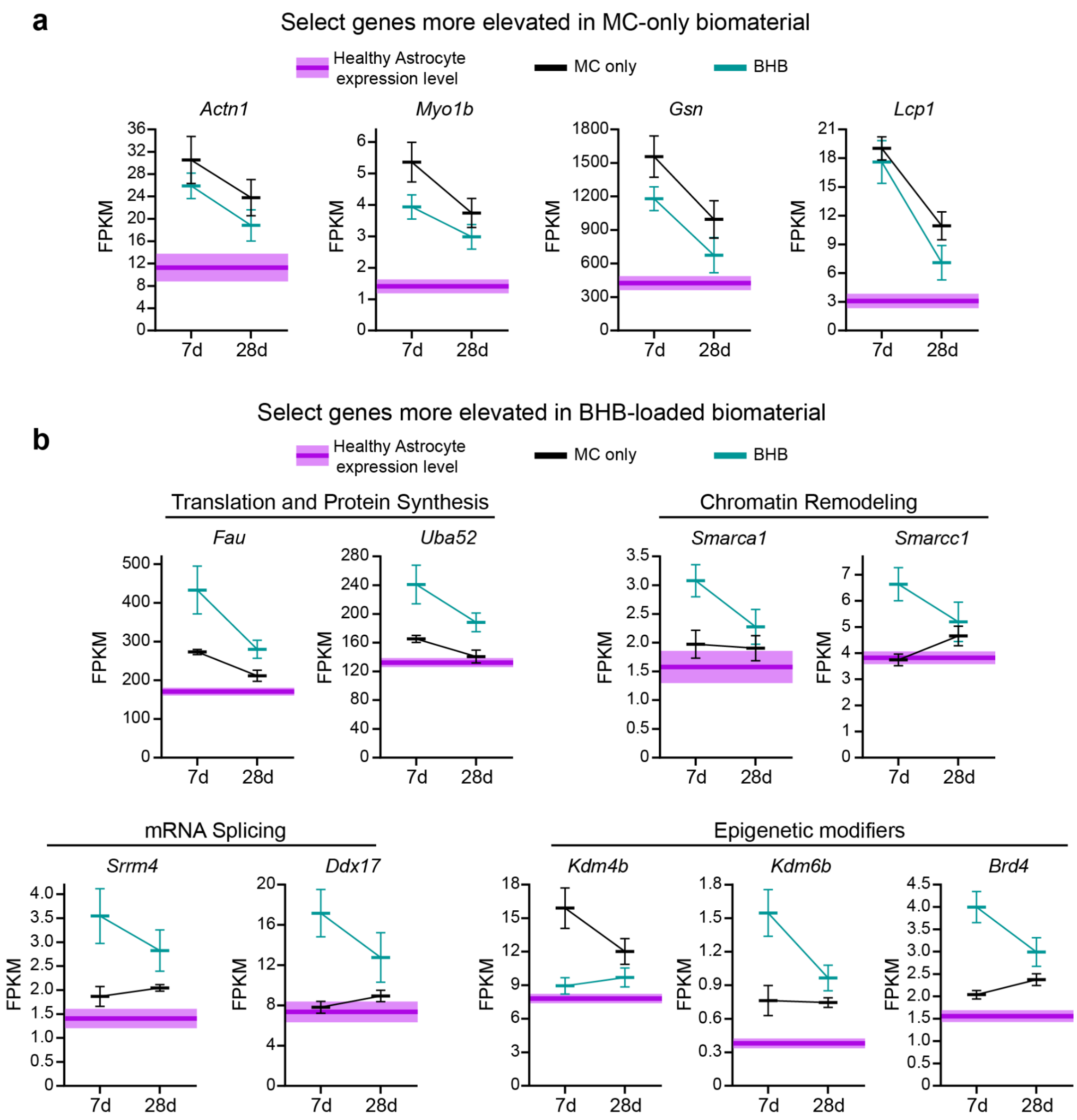


**Supplementary Fig. 24 | Comparison of select genes that are more highly upregulated in either MC-only or BHB-containing materials.**

**a.-b.** FPKM values for select DEGs representing **a.** genes that are more highly upregulated in MC-only biomaterials and **b.** genes that are more highly upregulated in BHB-containing biomaterials across the MC-only and BHB-loaded biomaterial conditions at 7 and 28 dpi. All graphs in this figure show mean ± s.e.m. with: Healthy (n=4), 7d/28d MC (n = 4), 7d BHB (n = 5), and 28d BHB (n = 4).

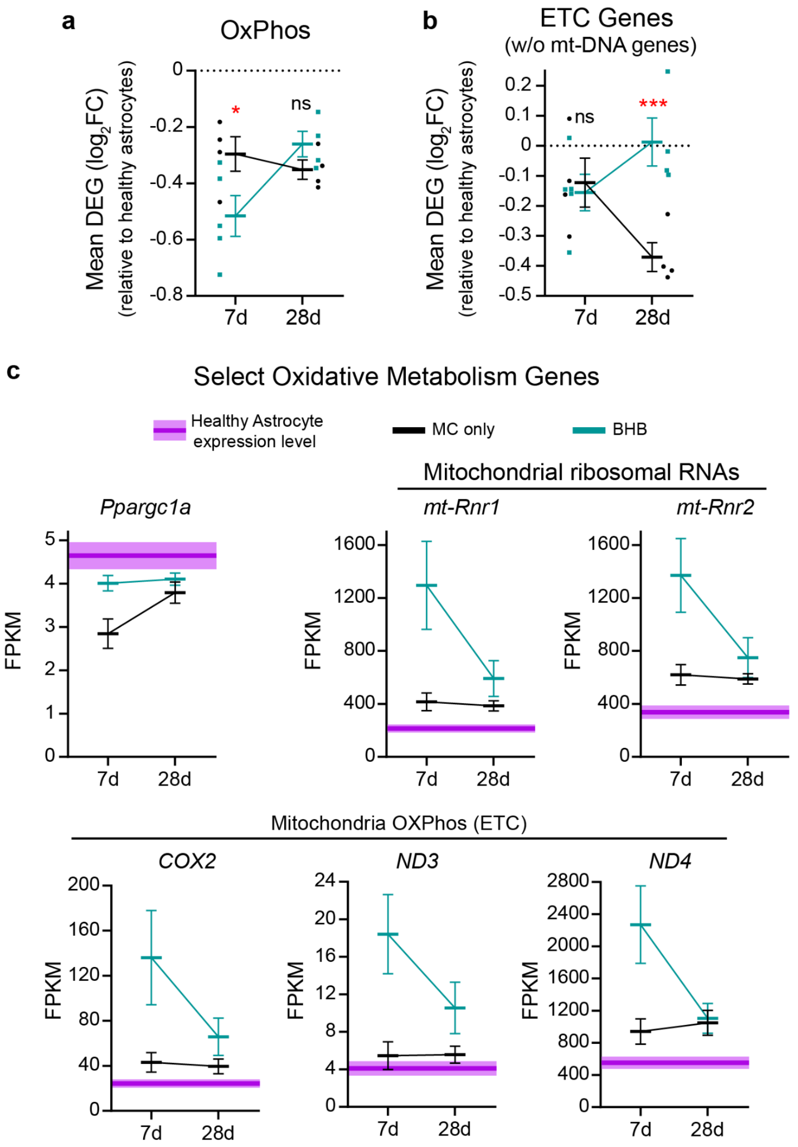


#### Supplementary Fig. 25 | Releasing BHB from MC deposits *in vivo* results in differential regulation of Oxidative Phosphorylation and ETC-associated genes.

**a.-b.** Mean differential expression ($log_2FC$) across the MC-only and BHB-loaded biomaterials at 7 and 28 dpi showing DEGs for the **a.** Oxidative Phosphorylation- and **b.** Electron Transport Chain (ETC, without mt-DNA genes)-associated gene lists. **c**. FPKM values for select Oxidative Metabolism-related DEGs across the MC-only and BHB-loaded biomaterial conditions at 7 and 28 dpi. All graphs in this figure show not significant (ns), $*P < 0.05$, and $***P < 0.002$. Comparisons are made between groups at a single timepoint via two-way ANOVA with Tukey's multiple comparison test. Graph shows mean ± S.E.M (**a.-c.**). with individual data points (**a.-b.**) showing: Healthy (n=4), 7d/28d MC (n = 4), 7d BHB (n = 5), and 28d BHB (n = 4).

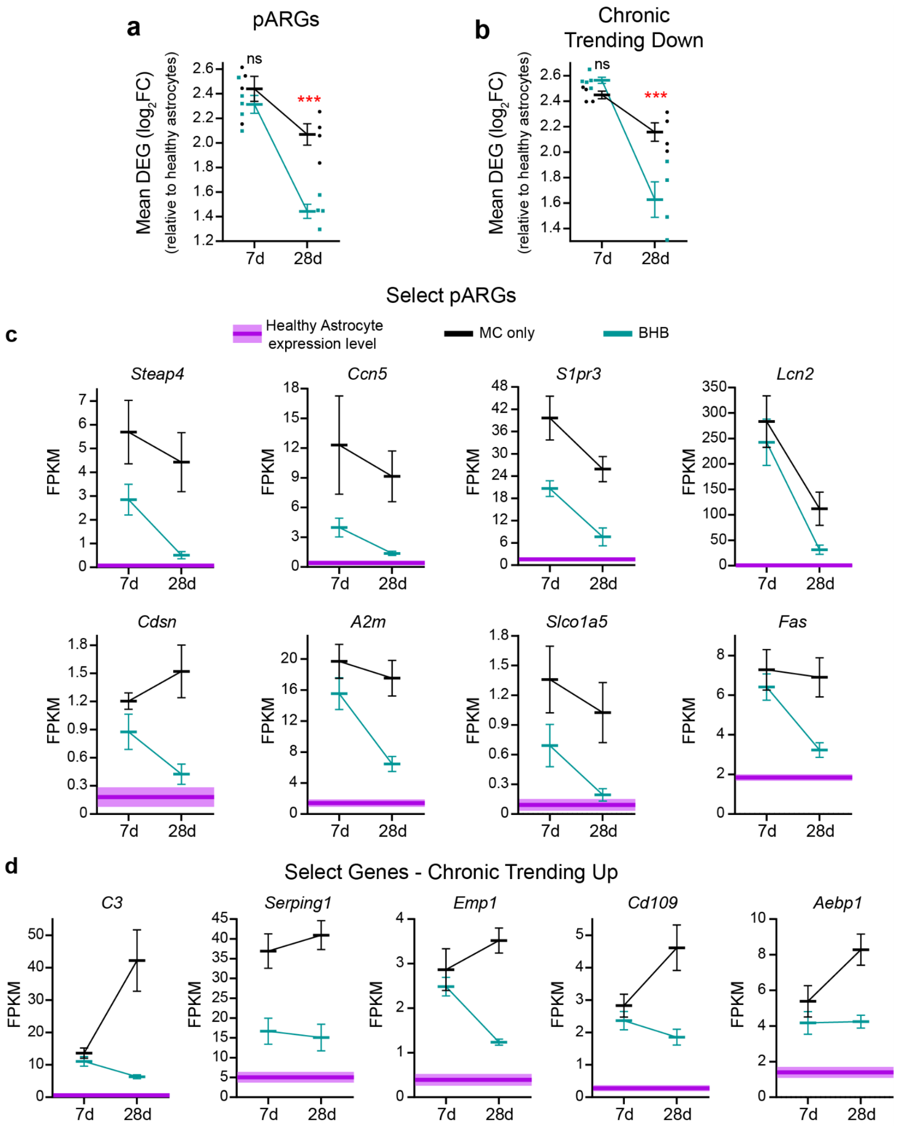


**Supplementary Fig. 26 | Releasing BHB from MC deposits *in vivo* results in downregulation of pARGs and Chronic Trending Down genes at 28 dpi.**

**a.-b.** Mean differential expression ($log_2FC$) across the MC-only and BHB-loaded biomaterials at 7 and 28 dpi showing DEGs for **a.** pARG and **b.** Chronic Trending Down gene lists. **c.-d**. FPKM values for select **c.** pARG-related DEGs and **d.** Chronic Trending Up genes across the MC-only and BHB-loaded biomaterial conditions at 7 and 28 dpi. All graphs in this figure show not significant (ns), ***$P < 0.0007$. Comparisons are made between groups at a single timepoint via two-way ANOVA with Tukey's multiple comparison test. Graph shows mean ± S.E.M (**a.-d.**) with individual data points (**a.-b.**) showing: Healthy (n=4), 7d/28d MC (n = 4), 7d BHB (n = 5), and 28d BHB (n = 4).
